# Comprehensive Profiling of Age- and Immune Cell–Specific Signaling Activation Using Multiplex Phosphoflow

**DOI:** 10.64898/2026.05.24.727113

**Authors:** Petra Hadlova, Michael Svaton, Katerina Kochmannova, Jakov Korzhenevich, Franziska Schmidt, Stefan F.H. Neys, Marei-Theresa Bott, Petra Vrabcova, Julian Staniek, Marketa Bloomfield, Tomas Kalina, Marta Rizzi

**Author notes:** Corresponding authors: Marta Rizzi, Tomas Kalina. Contributed equally (Tomas Kalina, Marta Rizzi).

## Abstract

Immune phenotyping represents a pillar in diagnostics, characterization of new genetic defects, and understanding mechanisms of diseases. Cell population distribution often does not cover the intrinsic function changes that may contribute to disease. Outcome of signaling activation can be used as proxy for cell function. To overcome the limitation of sample availability and standardization of signaling assays, we developed a multiplex full spectrum cytometry phosphoflow assay allowing the study of 6 phospho-proteins representing BCR/TCR, MAPK, PI3K/Akt/mTOR and canonical NF-κB signaling pathways in 18 immune cell subpopulations. Maximal stimulation and temporal dynamics were studied in response to pan-stimuli, activating cells regardless of receptor, and targeted stimuli for T, B, and innate immune cells. We studied healthy individuals between 1-69 years and discovered subpopulations-specific responses. Furthermore, pediatric donors showed broad differences in B cell and T cell function compared to adults. Hence, we established a tool to assess multiple signaling pathways at once and provide age- and subpopulation-specific references for signaling outcome.

**Summary:** Multiplex full spectrum flow cytometry-based phosphoflow assay across 18 immune cell subpopulations, 6 phospho-proteins in response to 6 stimuli at 4 time points in individuals aged 1-69 years, reveals distinct age- and subpopulation-associated signaling patterns in magnitude and dynamics of pathways activation.

## Introduction

Changes in the composition of the peripheral blood cell compartment are often used to corroborate diagnosis, to assess disease status or to understand cellular contribution to pathogenesis. For example, B cells are absent in agammaglobulinemia, an immune deficiency caused by a mutation in the Bruton’s tyrosine kinase (*BTK*) gene (*1,2*). Aged-associated or extrafollicular B cells are expanded in active systemic lupus erythematosus (SLE) (*3–6*), in human immunodeficiency virus (HIV) infection (*7*), and in a subset of common variable immunodeficiency (CVID) patients (*8,9*). Reduced counts of T regulatory cells are observed in several autoimmune diseases (*10–12*), inflammatory bowel disease (*13*), but also in rare conditions such as *FOXP3* mutation (*14,15*). The phenotyping of the peripheral blood mononuclear cell (PBMC) compartment is based on the expression of markers that identify both cell type and differentiation stage. This approach, while informative and even diagnostic (*16*), fails to capture the cellular function. In fact, changes in cell activation and signaling can be associated with autoimmune disease, immunodeficiency, and cancer (*17–21*). For example, dysregulated PI3K/Akt/mTOR pathway is associated with enhanced extrafollicular B cell responses and the formation of autoantibodies (*22–25*). Also dysregulated B cell receptor (BCR) signaling is associated with disease activity in complex autoimmune diseases such as systemic sclerosis, rheumatoid arthritis and antineutrophil cytoplasmic antibody (ANCA)-associated vasculitis (*26–28*). Furthermore, the study of signaling activation can drive the therapy management or the identification of new therapeutic targets (*29,30*). Hence, the study of signaling pathways complements phenotypic analysis, adding functional information that helps to understand disease status or pathogenesis.

The challenge in functional phenotyping is to define the cellular model, the type and timing of stimulation, and the relevant signaling pathway, which will provide information about the specific diagnosis. Additionally, assay standardization and the definition of reference controls is critical to the generation of robust data. The pFlow assay allows for a broad comparison across cell types and simultaneous analysis of multiple signaling pathway outcomes, overcoming the challenge of selecting the optimal molecules and time points. Age influences immune function, in fact, beside changes in cell composition (*31,32*), with children having more naïve T and B cells (*33,34*), the response to stimuli is different, as suggested by accelerated dynamic of activation in cord blood B cells (*35*), and the differential response to a similar novel infection cue, as shown by COVID19 infection studies (*36*). Hence, the study of age-related outcome of stimulation across populations is crucial to understanding outcome of immune responses.

We present a multiplexed in vitro pFlow assay to study the magnitude and temporal dynamics of the response to six different stimuli by analyzing 4 signaling pathways: the B and T cell receptor (BCR/TCR), Mitogen-Activated Protein Kinase (MAPK), phosphoinositide 3-kinase/ AKR mice strain thymoma/Mechanistic Target of Rapamycin (PI3K/Akt/mTOR), and canonical Nuclear Factor-kappa B (NF-κB) signaling pathway by the phosphorylation of 6 proteins: BTK/interleukin-2-inducible T-cell kinase (ITK), Akt, p38, S6, and p65) in 18 different cell populations from PBMCs at a single cell level. Applying this assay in 54 healthy individuals spanning the age range between 1-69 years old we highlight distinct characteristics in signaling responses across ages and distinct cell populations.

## Results

### Multiplex assay to assess multiple signaling pathways at a single cell level

The activation dynamics of distinct cell populations depends on the type of stimulus, cell responsiveness, the stimulation time, and the signaling pathway analyzed. Moreover, crosstalk between signaling pathways during activation can also influence the outcome. Full spectrum flow cytometry enabled a holistic analysis of cell activation and study of signaling outcomes in 18 distinct cell populations (Supplementary Figure 1). Additionally, barcoding allowed to multiplex the assay, analyzing all the cells stimulated with the same stimulus at all timepoints together (Figure 1A). The barcoding approach minimized differences in staining among samples across time and facilitated the comparison of phosphorylation induction along the time axis in response to defined stimuli (*37*). For detailed information about antibodies included in the panel and the used stimuli, see Supplementary Table 1.

**Figure 1.**
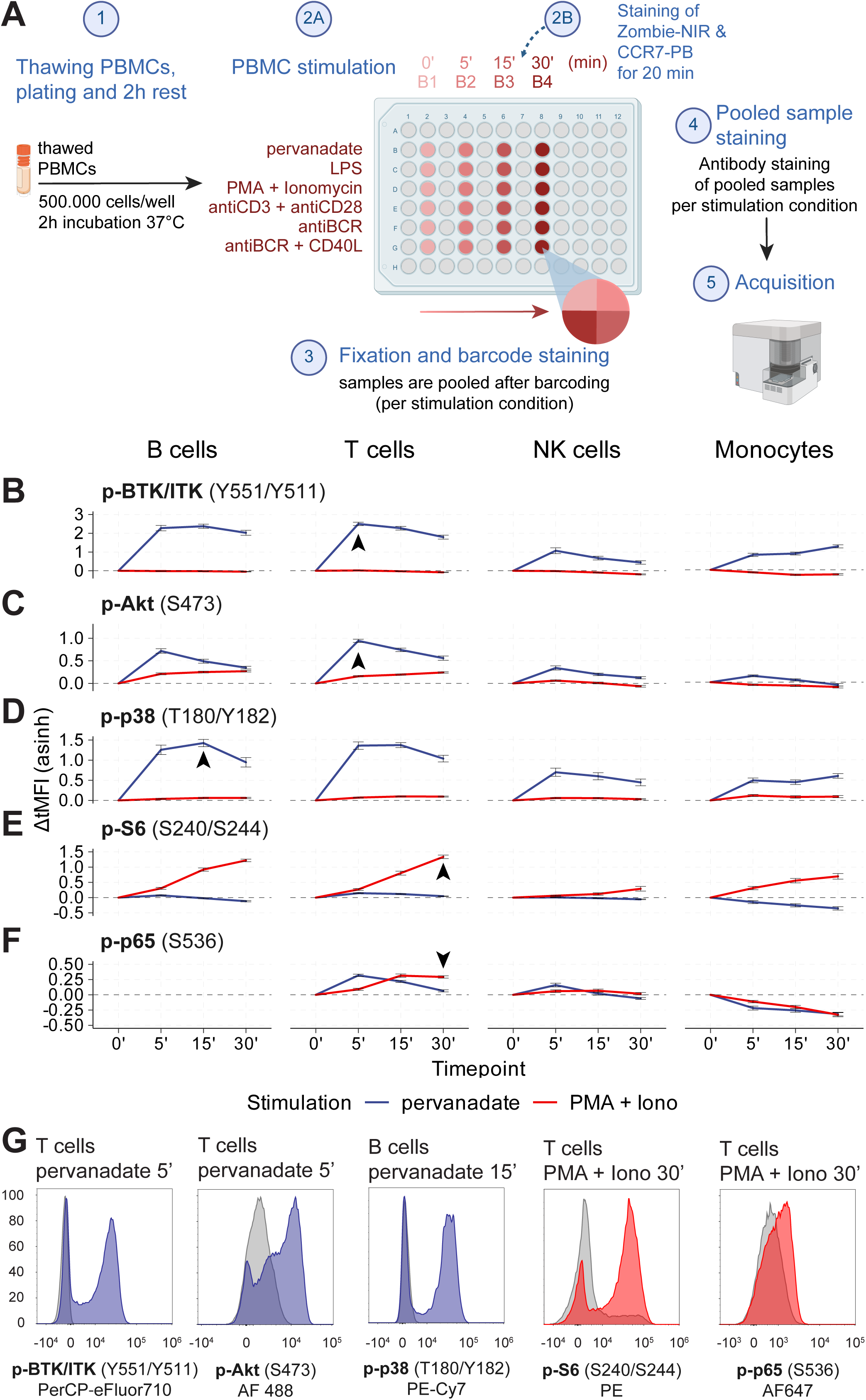
Pervanadate triggering proximal signaling events and PMA+Ionomycin distal signaling events in B, T, NK cells and monocytes. **(A)** Scheme of the pFlow assay. PBMCs isolated from peripheral blood. The plate depicts a representative experimental layout used for each donor. The timepoints (columns) from left to right – unstimulated (0‘), 5-minute (5‘), 15-minute (15‘) and 30-minute (30‘) stimulation. Each timepoint has one barcode assigned for all stimulations. Barcodes are indicated above each column by their code (B1-B4). Each row represents a different stimulus (indicated to the left of the plate). After barcoding, cells from one stimulus are pooled into one well (symbolized by a circle split into equal quarters). All subsequent surface and intracellular staining is performed on the pooled cells = 1 well per stimulus (6 total per donor). Cells are measured using the 5-laser full-spectrum cytometer, Cytek Aurora. Maximum inducible signal for each phospho-protein using pan-stimuli, pervanadate (blue) and PMA + Ionomycin (red). Signal is shown as transformed MFI change (ΔtMFI) on the y-axis, with the timepoint of stimulation indicated on the x-axis. Each row is one phospho-protein (phosphorylation site indicated in parentheses) **(B)** p-BTK/ITK, **(C)** p-Akt, **(D)** p-p38, **(E)** p-S6, **(F)** p-p65 each of the four major populations T and B cells (adaptive immune cells) and NK cells and monocytes (innate immune cells) is shown in columns. The black arrow always indicates the maximum induction reached across cell types for each phospho-protein. **(G)** Representative healthy donor histogram plots showing the maximum signal induction for each phospho-protein with population, time and stimulus indication above the plots. pervanadate (blue), PMA + Ionomycin (red). The phospho-proteins are indicated below individual histogram plots together with the phosphorylation site and fluorochrome.

As pan-stimuli, we selected the protein tyrosine phosphatase (PTP) inhibitor pervanadate (*38*) and phorbol 12-myristate 13-acetate (PMA), a strong activator of protein kinase C (PKC), in combination with Ionomycin, an ionophore, which triggers the Ca^2+^-calmodulin-dependent signaling pathway (*39*). Pervanadate blocks the activity of PTPs, which results in accumulation of phosphorylated proteins and induction of proximal signaling events (*40,41*). These include phosphorylation of BTK/ITK (Y551/Y511), Akt (S473) and p38 (T180/Y182) in both innate and adaptive cell subpopulations (*42–45*) PMA + Ionomycin induce more distal signaling events such as phosphorylation of S6 (p-S6, S240/S244) (*46*) and p65 NF-κB activation (S536) (*47,48*).

To compare the effects of stimulation across all timepoints, cell subpopulations and individuals, we used a standardized pipeline to analyze the data (see Methods). For the analysis of individual phospho-proteins, we applied an inverse hyperbolic sine (arcsinh) transformation to the data and calculated the median fluorescent intensity from the transformed values (tMFI) for each cell subpopulation and stimulation at different timepoints (Supplementary Figure 2). The induction of the signal after stimulation, representing the change relative to the unstimulated state at each timepoint, was calculated as the difference (ΔtMFI) between the individual tMFI values and the median of the tMFI values from the unstimulated samples. The overall activation of each signaling pathway was quantified as the area under the curve (AUC) from the ΔtMFI across all timepoints.

We assessed stimulation responses to pan-stimuli in 54 healthy donors (Supplementary Table 1). When looking at phosphorylation of the signaling intermediates, the maximum intensity of signal was different in distinct populations (Supplementary Figure 2) and showed a different pattern depending on the stimulus. To compare across stimuli, we analyzed the induction of signaling. The maximum ΔtMFI for p-BTK/ITK (Figure 1B), p-Akt (Figure 1C), and p-p38 (Figure 1D) was reached after 5-and 15 minute-stimulation with pervanadate across all cell types. The highest mean ΔtMFI for p-BTK/ITK and p-Akt was measured after 5 minutes in T cells reaching 2.5 and 0.94 respectively. The highest mean ΔtMFI for p-p38 was measured after 15 minutes in B cells reaching 1.43. For all other ΔtMFI values, see Supplementary Table 2. While B and T cells showed persistent activation over 15 and 30 minutes, NK cell rapidly returned to baseline (Figure 1B-D). These differences could be due to different availability of BTK/ITK, Akt, and p38 proteins as signaling intermediates in the different cell types, or to a difference in the activity of the phosphatases targeted by pervanadate such as SHP-1, SHP-2 and PTPN22 (*49,50*).

The stimulation with PMA + Ionomycin resulted in a slower induction of S6 phosphorylation with the maximum signal detected after 30 minutes in T cells, B cells, and monocytes (Figure 1E). T cells showed the maximal ΔtMFI mean for p-S6 at 1.34. p-S6 was only minimally induced in NK cells. NF- κB was not strongly induced by either of the pan-stimuli (Figure 1F). Nonetheless, PMA+Ionomycin induced the peak NF-κB p65 phosphorylation in T cells, after 30 minutes (mean ΔtMFI 0.39). Signal induction in B and T cells was greater than in NK cells and monocytes in all five phospho-proteins (Figure 1B-F). As expected, stimulation with PMA+Ionomycin did not induce proximal signaling and pervanadate had little effect on the distal signaling phospho-proteins. We excluded p-p65 expression in CD19^+^ B cells from the analysis because of spillover of the fluorochromes.

The stimulation inducing maximum phosphorylation (Figure 1G) resulted in bimodal distribution in most cell subpopulations (Supplementary Figure 3). Major cell populations, such as B or T cells, include distinct subpopulations (Supplementary Figure 1), that have been reported to respond differently to stimuli, and may therefore influence the outcome of activation (*51,52*). Hence, when pervanadate and PMA+Ionomycin are used as positive controls, it is crucial to focus on the selection of cell population, the phospho-residue of target proteins, and the temporal dynamics to correctly assess the assay outcome and performance.

### Cell-type analysis reveals subpopulation-specific activation

It has been previously shown that distinct subpopulations within a cell type may respond differently to stimuli (*53–55*). Therefore, we compared the dynamics of signal induction relative to the unstimulated cells in CD27^-^IgD^+^ naïve and CD27^+^IgD^-^ switched memory (SM) B cells and in naïve and memory T cells defined by the differential expression of CD45RA and CCR7 (Supplementary Figure 1).

The analysis of the dynamic of stimulation with the calculation of the AUC of the ΔtMFI showed a similar outcome of stimulation of p-BTK, p-Akt and p-p38 in naïve and SM B cells in response to pan-stimuli, except for p-S6 induced by PMA + Ionomycin (Figure 2A). SM B cells showed a significantly higher p-S6 induction at 15 and 30 minutes compared to naïve B cells. In response to pervanadate CD4^+^ (Figure 2B) and CD8^+^ (Figure 2C) naïve T cells showed a higher response in p-Akt and p-p38 compared to memory cells, as depicted by the AUC, except for p-ITK that was increased only in naïve CD4^+^ T cells. CD4^+^ naïve T cells showed higher p-ITK induction at 5 minutes that persisted up to 30 minutes. In contrast to B cells, the response (AUC) to PMA + Ionomycin in p-S6 induction was similar across T cell subpopulations. The induction of p65 phosphorylation was higher in naïve CD4^+^ and CD8^+^ T cells both at 15 minutes and 30 minutes of stimulation with PMA + Ionomycin compared to CD4^+^ and CD8^+^ memory T cells. For specific ΔtMFI values, see Supplementary Table 2.

**Figure 2.**
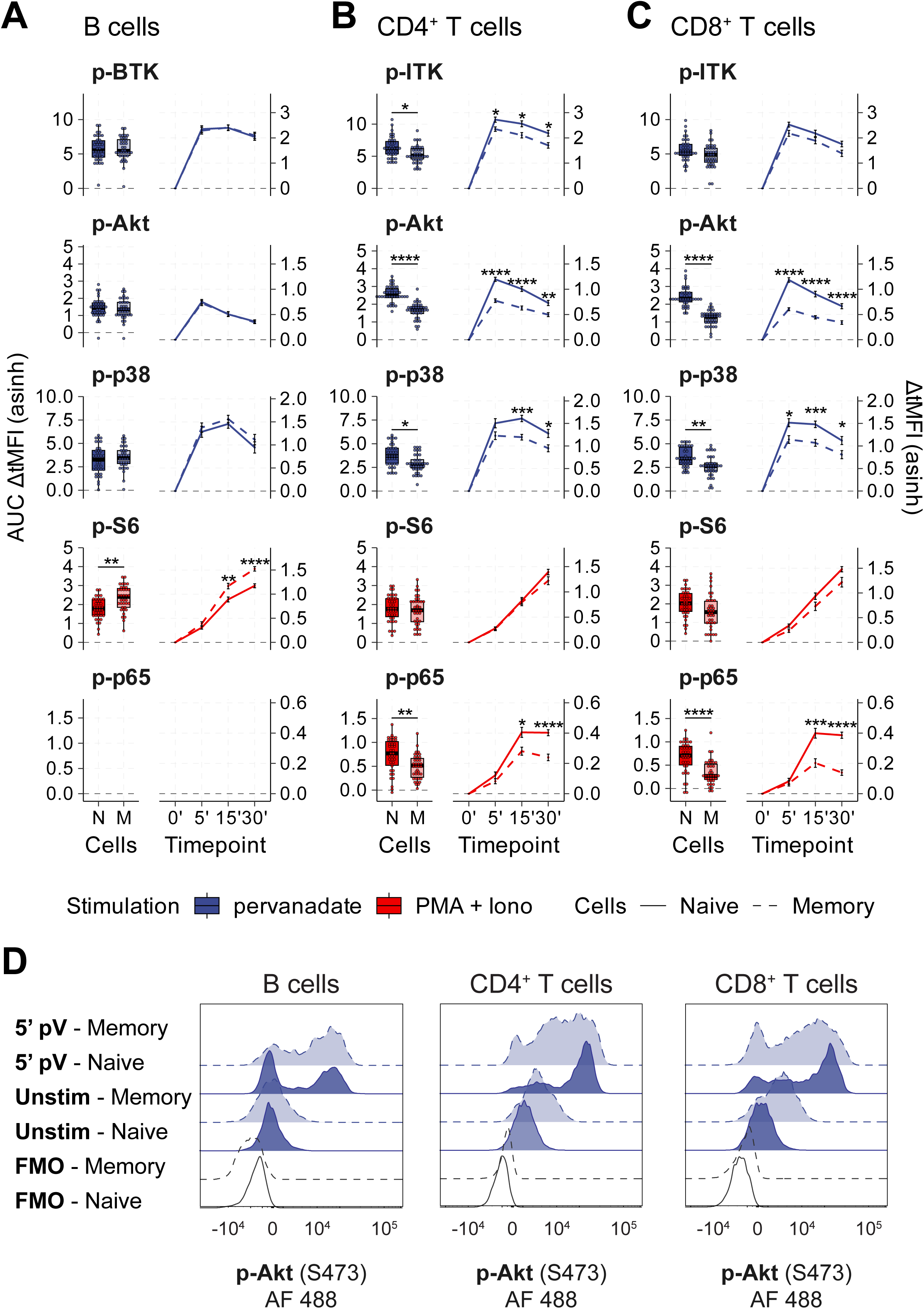
Cell subpopulation-specific activation. Pan-stimuli induced subpopulation-specific signal induction. Pervanadate stimulation is depicted in blue and PMA + Ionomycin in red. Each column represents one major subpopulation, from left to right, **(A)** B cells, **(B)** CD4^+^ and **(C)** CD8^+^ T cells. Each row represents one phospho-protein. Within each population naïve (full line) vs SM B cells and memory T cells (dashed line) are to be distinguished. Two parts of each graph represent the area under curve (AUC) for each phospho-protein and population (left side) and curve of ΔtMFI for kinetics (right side). **(D)** Illustrative healthy donor histogram plots of p-Akt signal induced by 5-minute stimulation with pervanadate (5’ pV) in comparison to unstimulated cells (Unstim) and FMO controls (FMO). Memory (light blue, dashed line) and naïve (dark blue, full line) cell subpopulations are depicted for B cells, CD4^+^ and CD8^+^ T cells (columns from left to right). The statistical analysis was performed using unpaired Welch’s t test with Benjamini-Hochberg correction. P-value is indicated by asterisks above graphs for significant observations only, P<0.05 (*), P<0.01 (**), P<0.001(***), P<0.0001 (****).

The amplitude of the induction of activation is influenced by the intrinsic level of phosphorylated molecules for each signaling pathway. In line with previous reports (*27,56*), memory cells showed higher basal levels of phosphorylation of BTK/ITK in B and CD4^+^ T cells, Akt in CD4^+^ and CD8^+^ T cells, p38 in CD4^+^ T cells, S6 in B and CD4^+^ T cells compared to their naïve counterparts (Supplementary Figure 4). Additionally, upon activation with pan-stimuli, the memory populations did not reach tMFI as high as the naïve populations. This observation explains the higher induction levels in naïve in contrast to memory T cells. Of note, as observed for the major cell populations, the activation with pan-stimuli resulted in a bimodal signal distribution as demonstrated for p-Akt after 5-minute stimulation with pervanadate (Figure 2D). The observed bimodal distribution indicates the presence of subpopulations exhibiting distinct sensitivities to pan-stimulation within the naïve or memory compartments.

### Enhanced response to BCR stimulation in memory B cells

The similar dynamic of activation between naïve and SM B cells in response to pervanadate prompted us to analyze the signal induction upon anti-IgA/M/G (anti-BCR) stimulation. Indeed, memory B cells are known to promptly react to activation (*26,52,54*). In response to anti-BCR stimulation, SM B cells showed a stronger overall induction of phosphorylation of four proteins BTK, Akt, p38, and S6 as demonstrated by the AUC plots (Figure 3A) in line with previously published data (*28,57*). Signaling dynamics showed higher and faster induction of p-BTK, p-Akt, p-p38, and p-S6 in SM B cells that persisted over all timepoints compared to naïve B cells (Supplementary Figure 5A).

**Figure 3.**
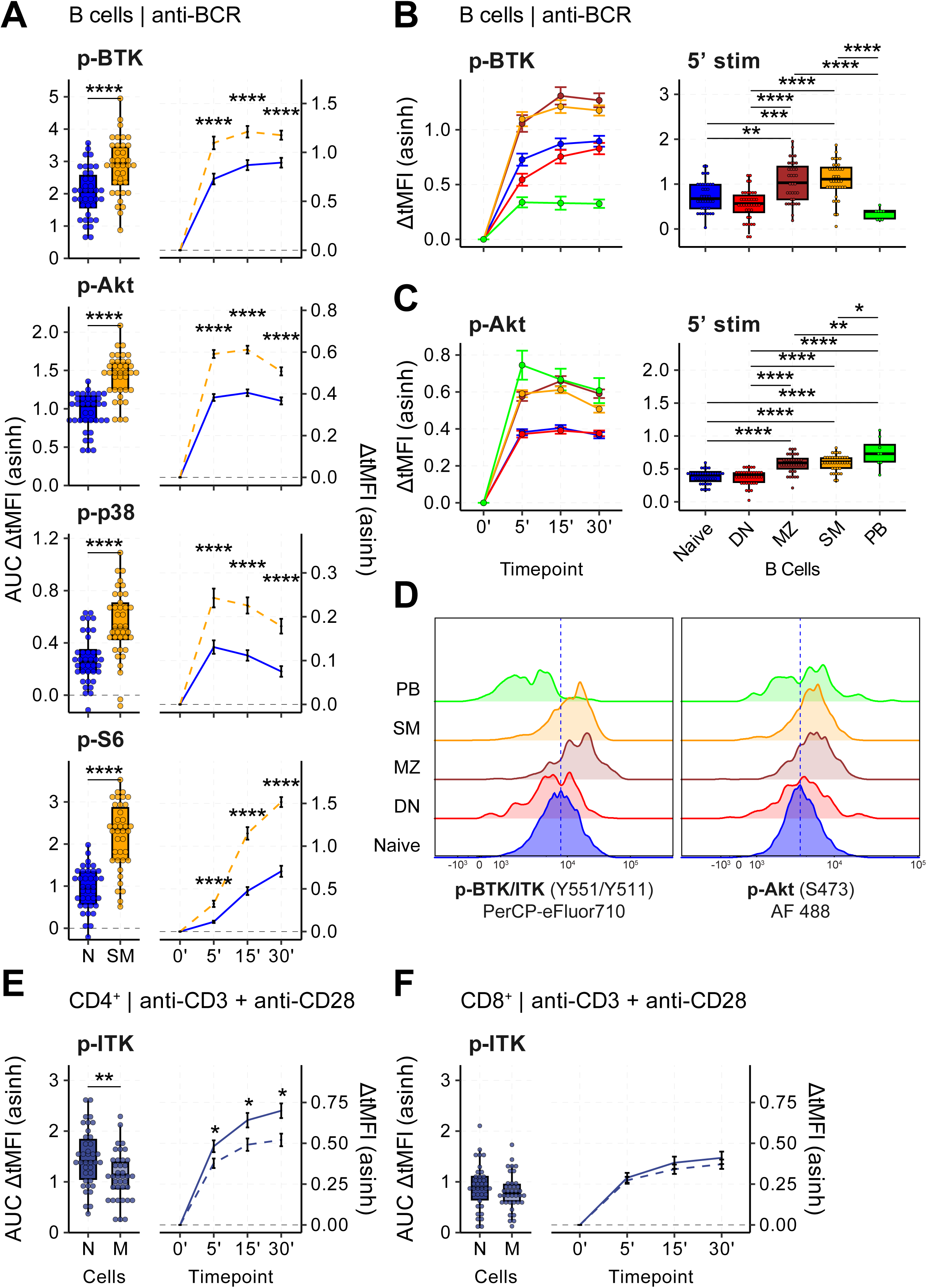
Cell type-specific stimulation shows differences in B and T cell subpopulation activation. Signal induction by cell-specific stimulation in B cells (by anti-BCR stimulation) and T cells (by anti-CD3 + anti-CD28) and their naïve and memory subpopulations. AUC representation for each phospho-protein and population on the left and ΔtMFI kinetics of signaling on the right. **(A)** B cells stimulated by anti-BCR stimulation of naïve (blue) and switched memory (yellow) cells. **(B, C)** Identification of distinct activation levels across the B cell subpopulations stimulated with anti-BCR for 5 minutes as observed in **(B)** p-BTK and **(C)** p-Akt. Each B cell subpopulation is depicted in a different color (naïve-blue, SM - yellow, DN-red, MZ - dark red, PB - green). **(D)** Representative healthy donor histogram plots of p-BTK (left) and p-Akt (right). **(E, F)** p-ITK signal after anti-CD3 + anti-CD28 stimulation as measured in naïve (dark blue, full line) and memory (light blue, dashed line) (**E)** CD4^+^ and **(F)** CD8^+^ T cell naïve and memory subpopulations. The statistical analysis was performed using unpaired Welch’s t test with Benjamini-Hochberg correction. P-value is indicated by asterisks above graphs for significant observations only, P<0.05 (*), P<0.01 (**), P<0.001(***), P<0.0001 (****).

Our multiplex panel allowed the distinction of further B cell differentiation stages beside naïve and SM B cells, in particular double negative-(DN) B cells, which contain non-conventional memory cells, marginal zone (MZ)-like B cells that respond to T-independent antigens and plasmablasts (PB) (Supplementary Figure 1). In response to anti-BCR stimulation, MZ and SM B cells showed the highest intensity and dynamics of ΔtMFI for p-BTK at 15 minutes (Figure 3B). The induction of signaling in naïve and DN B was slower reaching the highest values after 30 minutes and lower to about half of the SM and MZ B cells (Figure 3B). PB showed the lowest response reaching their maximum induction after 5 minutes.

p-Akt also showed distinct patterns of induction (Figure 3C). In response to BCR stimulation, MZ and SM B cells and PB, showed higher ΔtMFI values than naïve B cells and DN B cells. PB were faster and reached their maximum after 5 minutes compared to the other four subpopulations, which reached their peak at 15 minutes (Figure 3C). The analysis of single cell subpopulations contained within the CD19 gate showed homogeneous distribution of the activation, as indicated by the histogram plots (Figure 3D). This supported the idea that bimodal distribution of activation may often be driven by distinct dynamic of activation in different cells included in the analysis gate.

In the T cell compartment, prior to stimulation, ITK phosphorylation level was higher in memory, compared to naïve CD4^+^ T cells (Supplementary Figure 5B), as were pAkt, p-p38 and p-S6. p-ITK was strongly induced by anti-CD3 + anti-CD28, both in CD4^+^ and in CD8^+^ T cells, but none of the other signaling molecules (Supplementary Figure 5B). Naïve CD4^+^ T cells showed a higher overall signal of p-ITK when stimulated via anti-CD3 + anti-CD28 compared to memory CD4^+^ T cells, as shown by the AUC (Figure 3E). Significantly higher p-ITK induction was detected at all timepoints for naïve CD4^+^ T cells over memory CD4^+^ T cells. There was no significant difference neither in the overall signal nor in the dynamics for the CD8^+^ T cell subpopulations (Figure 3F). Hence, different subpopulations show specific responsiveness patterns with unique dynamics, intensities, and (potentially) functional outcome.

### Maximum induction and signal across all cell compartments

To study signaling responses in diseases or in response to signaling inhibitor therapies, choosing the level of phosphorylation, the kind of stimulus used, and the specific cell type is crucial. To this end, we compared the level of phospho-protein induction, (ΔtMFI) related to unstimulated cells (Figure 4A) and the maximum tMFI (Figure 4B) across 14 immune cell populations at all timepoints after all types of stimulation. We selected the stimulation and timepoint at which the median of the ΔtMFI reached the highest value for each phospho-protein and each cell population.

**Figure 4.**
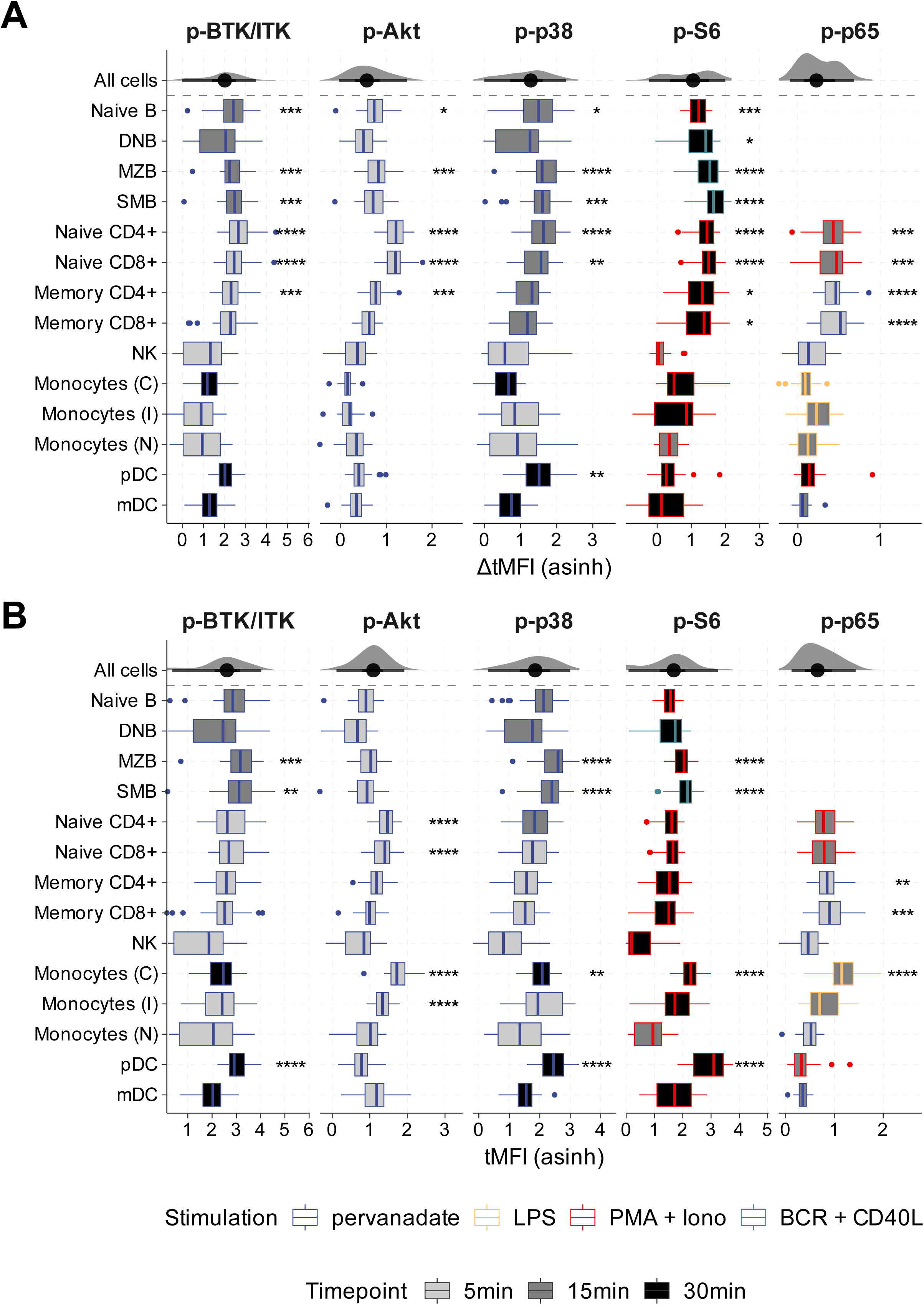
Maximum signal induction across cell populations and stimuli. Maximum **(A)** ΔtMFI and **(B)** tMFI for all subpopulations (rows) for each phospho-protein (column) with indicated stimulation. Each box plot contains the information on stimulation (outline color; pervanadate – blue, PMA + Ionomycin – red, LPS-yellow, BCR+CD40L-green) and selected timepoint (fill color; 5min - light grey, 15min - dark grey and 30min - black). The statistical analysis was performed using unpaired Welch’s t test, comparing each cell population to the combined data for selected stimulations and timepoints of all cells (top row). P-value is indicated by asterisks above graphs for significant observations only, P<0.05 (*), P<0.01 (**), P<0.001(***), P<0.0001 (****).

As expected, the proximal signaling events were all maximally induced by pervanadate. The highest ΔtMFI for p-BTK/p-ITK was reached in naïve, SM and MZ B cells, and in naïve, and memory CD4^+^ T cells and naïve CD8^+^ T cells (Figure 4A). The maximum tMFI for BTK was observed in SM and MZ B cells, and plasmacytoid dendritic cells (pDCs) (Figure 4B). Induction of p-Akt was highest in naïve and memory CD4^+^ and naïve CD8^+^ T cells, in naïve and MZ B cells (Figure 4A), while the maximal tMFI was observed in naïve CD4^+^ and CD8^+^ T cells, and in classical and intermediate monocytes (Figure 4B). p-p38 induction was highest in MZ, SM and naïve B cells, naïve CD4^+^, and CD8^+^ T cells, and pDCs (Figure 4A). Peak tMFI for p-p38 was observed in MZ and SM B cells, classical monocytes and pDCs (Figure 4B).

Of note, when analyzing the distal signaling events, PMA + Ionomycin stimulation did not result in the highest induction in all cell subpopulations. Maximum detected p-S6 ΔtMFI was reached after 30 minutes of stimulation in T and B cell subpopulations (Figure 4A). However, the highest tMFI in DN, MZ and SM B cells was observed after anti-BCR + CD40L stimulation (Figure 4B) as opposed to the PMA + Ionomycin, which maximally induced T cell subpopulations. The highest tMFI of p-S6 was reached in SM and MZ B cells, classical monocytes and pDCs in response to PMA + Ionomycin.

Canonical NF-κB signaling, represented by p-p65, was induced the highest in both naïve and memory CD4^+^ and CD8^+^ T cell subpopulations (Figure 4A). The maximum ΔtMFI in naïve T cell subpopulations was reached using PMA+Ionomycin while in memory T cell populations the maximum signal was induced by pervanadate, and in innate immune cells by LPS. The highest tMFI was reached in classical monocytes in response to 15 minutes of LPS stimulation and in memory CD4^+^ and CD8^+^ T cells after pervanadate stimulation. Notably, despite the 2-hour resting time in our protocol, monocytes had a very high background in the measured phospho-proteins, as shown by tMFI in Figure 4B, corresponding with a relatively lower ΔtMFI (Figure 4A), indicating the limited capacity for further induction. Together, our analysis provides a powerful resource to inform future studies in the selection of phospho-proteins, the immune cell populations, timepoints, and types of stimuli for the optimal analysis of signaling responses in the model of interest.

### Naïve cells of children show higher induction of signaling

Aging may influence the intrinsic ability of immune cells including lymphocytes and myeloid cells to respond to stimuli. Indeed, cord blood-derived naïve B cells are more prone to react to BCR stimulation compared to adult naïve B cells as demonstrated by higher cytokine secretion and surface marker expression (*35,58,59*). We assessed changes in PBMC composition and signaling induction across all ages. We distinguished pediatric donors (0-18 years old) and divided the adults into two groups: 18-55 years old and above 56 years old. With age, we observed a progressive decrease in naïve B, CD4^+^ and CD8^+^ T cell frequencies, while increased percentages of SM B cells (Figure 5A), as previously described (*33,34,60*). The frequency of naïve B cells decreased from 79.2% (median) in the pediatric cohort to 61.7% in the elderly, while SM B cells increased from 7% to 15.1%, respectively. Similarly, the frequency of naïve T cell subpopulations decreased from 73.2% to 44.1% in the CD4^+^ subpopulation, and from 69.6% to 31.9% in the CD8^+^ subpopulation.

**Figure 5.**
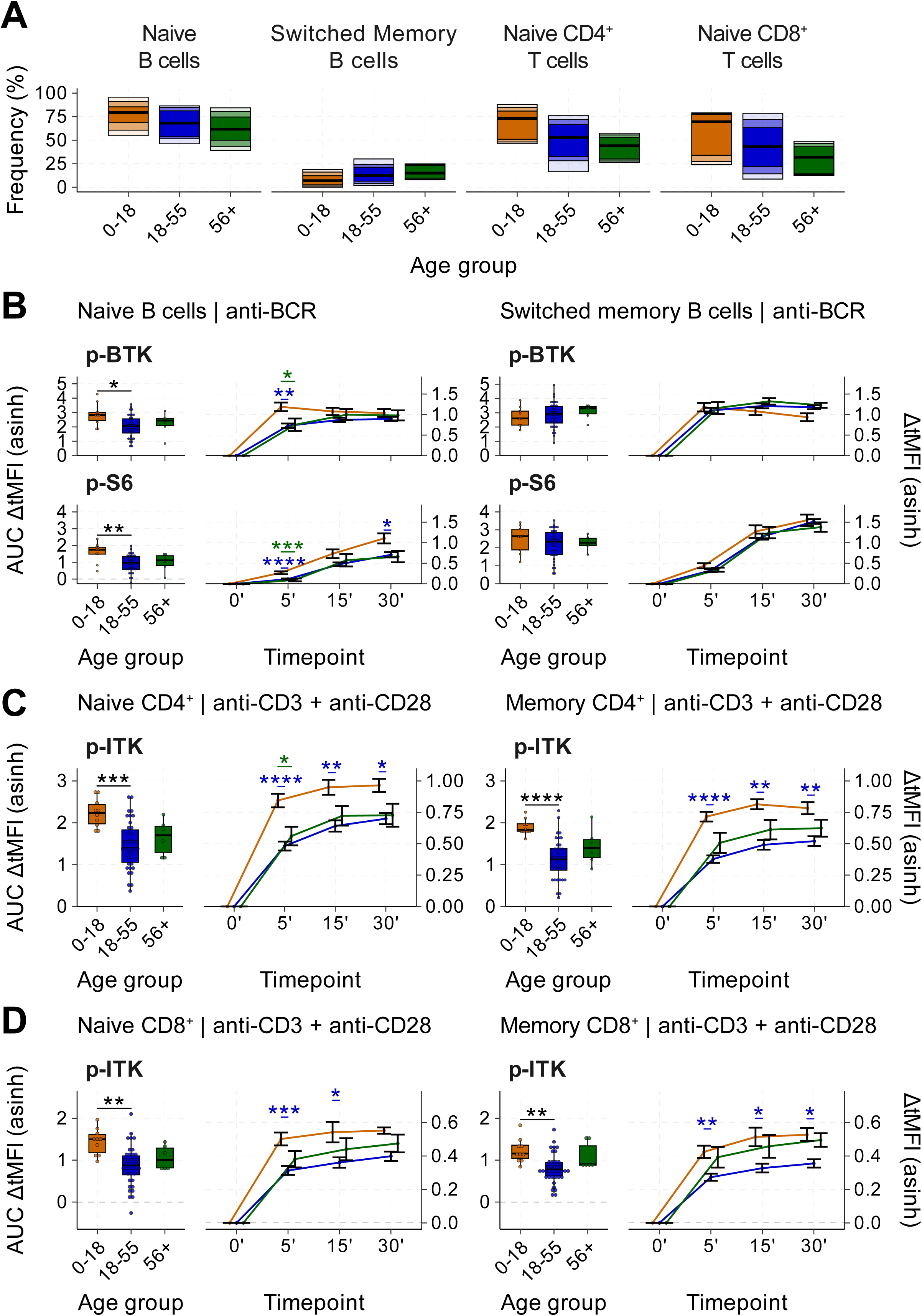
Age affects signal induction by pan-stimuli. **(A)** Boxplots depicting distribution of cells according to age of donor; 0-18yo (orange), 18-55yo (blue) and >56yo (green) as a relative frequency of total B cells for naïve and SM B cell subpopulations and for naïve T cells of total CD4^+^ and CD8^+^ T cells subpopulations. Mean frequency was calculated for each donor for all unstimulated samples. **(B)** Age differences in signaling induction of phospho-proteins downstream of BCR (p-BTK, p-S6) in naïve (left) and SM (right) B cells. Cells were stimulated with anti-BCR. AUC representation for each phospho-protein and population is shown on the left side and curve of ΔtMFI showing kinetics on the right side. Age differences in signaling induction of p-ITK as a representative of a phospho-protein downstream of TCR specific stimulation using anti-CD3 + anti-CD28 in naïve (left) and memory (right) **(C)** CD4^+^ **(D**) CD8^+^ T cells. Boxplots show the AUC (left y-axis) and the lines mean ΔtMFI with SEM at each timepoint (right y-axis). The statistical analysis was performed using One-way ANOVA with a Tukey’s post-hoc test in B–D. P-value is indicated by asterisks above graphs for significant observations only, P<0.05 (*), P<0.01 (**), P<0.001(***), P<0.0001 (****). For individual timepoints, differences between the 0-18 and 18-55 group are shown in blue, differences between 0-18 and 56+ groups are shown in green.

We studied the induction of signaling, focusing on differences within the same cell population among age groups in response to anti-BCR and anti-CD3 + anti-CD28 stimulation. Naïve B cells from pediatric donors showed a significantly increased induction of p-BTK compared to the adult cohort, as shown in the AUC box plots (Figure 5B). Pediatric naïve B cells respond faster, reaching maximal induction already at 5-minute stimulation with anti-BCR. At 15 and 30 minutes, the level of p-BTK induction in naïve B cells of pediatric donors and the adults reached same ΔtMFI levels. For p-S6 we observed a significantly increased signal in the pediatric cohort compared to the adults, with ΔtMFI significantly higher at 5 and 30 minutes. SM B cells showed similar induction patterns across all age groups with no significant differences.

Naïve and memory CD4^+^ T cells from pediatric individuals showed a significantly higher p-ITK induction both in the AUC and in the ΔtMFI for individual timepoints (Figure 5C) compared to the adults. Similar dynamics were observed in CD8^+^ T cells from pediatric donors (Figure 5D). Unlike for SM B cells, memory T cells from the pediatric cohort showed a significantly higher p-ITK induction at all timepoints. In summary, pediatric B and T cells not only have a significantly higher proportion of naïve subpopulations, but these also respond with higher intensity to targeted stimulation. Hence, age of donor is an important factor in signaling studies, which should be considered when comparing responses to different stimuli.

## Discussion

Immunophenotyping of peripheral immune cell subpopulations is often limited to defined populations, their relative frequency, and marker characteristics (*16,61–63*). Even though such information is valuable, it does not account for functional changes. Cell function is commonly tested by performing assays focused on proliferation (*64,65*), the changes in expression of surface activation markers (*64,66*), cytokine release (*67–69*), cytotoxicity (*70–72*) and cell signaling (*73–76*). Signaling studies are often done on adult donors, with cohort sizes of 10 to >160 (*33,77,78*). We presented a cohort of 54 individuals that contains healthy pediatric donors, allowing the direct comparison of signaling responses across ages. We showed increased activation levels in cells from pediatric individuals, with specific differences depending on the cell type.

The assessment of signaling provides crucial information in different disease and biological contexts. It is relevant in the study of cell development, differentiation, and activation in context of health (*57,73,74,79,80*) and disease (*8,9*), in identification of biomarkers in cancer using cytometry by time-of-flight (CyTOF) (*81*) or flow cytometry (*17,28,82,83*), in inborn errors of immunity and autoimmune diseases (*84–87*). Furthermore, it has been shown that the mTOR signaling pathway is a disease modulator in COVID-19 disease (*88,89*) and NF-κB signaling is important in disease progression (*90*). To exploit the power of signaling analysis in these contexts, enabling a holistic view on activation of distinct pathways, we introduced a full-spectrum cytometry multiplexed pFlow assay, presenting a valuable resource that enables the analysis of six phospho-proteins in response to six different stimuli, at four timepoints, in up to eighteen populations.

The activation with pan-stimuli (pervanadate and PMA + Ionomycin) is commonly used as positive controls in signaling studies (*91–93*). We show that the phosphorylation of proximal signaling molecules such as BTK/ITK, Akt, and p38 was induced by pervanadate, while the distal signaling proteins like S6 and p65 required activation with PMA + Ionomycin across all cell types. Our results highlight the importance of correct selection of stimulation and phospho-protein readout.

We found that naïve T cells show a higher induction of signaling compared to memory T cells in response to pan-stimuli. Pervanadate inhibits phosphatases. Activity and expression of phosphatases can control threshold of activation. Indeed, central memory T cells exhibit increased expression of protein tyrosine phosphatases non-receptor types (PTPN), more specifically PTPN2 (TC-PTP), PTPN12 (PTP-PEST), and PTPN22 (*94*), potentially contributing to lower signaling induction. The phosphatase PP2A regulates T cell activation (*95,96*) and is simultaneously essential for activation and germinal center formation in B cells (*97*), and plays a role in regulation of T-bet expression in NK cells (*98*). Hence, there is a cell-specific role of phosphatases in regulating activation, whereby levels of expression are essential for their function (*99,100*). Response to PMA + Ionomycin depends on sensitivity of cells to PKC activation and calcium signaling, commonly associated with T cell activation (*64,101,102*). We observed an overall higher signal induction in the distal phosphorylated - proteins induced by PMA + Ionomycin (i.e. S6 and p65) in T cells than in B cells. These data suggest that the availability of individual signaling intermediates is key for the strength of the induced signal (*46,103,104*).

Our work reveals differences not only in the response to pan-stimuli but also to targeted stimulation in different cell types and their subpopulations. Hence, when analyzing the response of total CD19^+^ B cells or total CD3^+^ T cells, the measured response will strongly depend on the relative proportion of each subpopulation in the studied individual. In fact, factors that contribute to the higher reactivity of SM B cells to anti-BCR are increased expression of BTK (*26*), Grb2 (*105,106*) and PTEN (*107*). Of note, our panel does not distinguish between IgA^+^ and IgG^+^ SM B cells, which are known to differ significantly in signaling pathway activation (*27,108*).

Even though naïve T cells require longer antigen stimulation and costimulatory signals to get activated (*109*), it has been shown that the threshold of activation via the TCR is higher in memory T cells (*94*). In line with this, our data showed that naïve T cells had higher signal induction than memory T cells and faster kinetics in response to pervanadate as well as antiCD3 + anti-CD28 stimulation. However, we observed that the maximum expression of phospho-proteins tMFI was higher in the memory and effector cells than in the naïve subpopulation. The increased baseline tMFI of the memory subpopulations is likely due the increased levels of signaling intermediates in effector and memory T cells, as previously reported (*110–112*)

Children show higher frequencies of naïve cell subpopulations, particularly in the naïve CD8^+^ T cells (*32*), in line with our data. Conversely, effector memory CD8^+^ T cells gradually increase throughout childhood into adult age (*59*) along with the accumulation of granzyme K-positive CD8^+^ T cells with an exhausted phenotype (*113*), accompanied by a decreased thymic output (*114,115*). As for B cells, their count as well as relative frequency decreases with age (*116*). On the other hand, in aging adults and in autoimmune disease such as SLE, increases in age-associated B cell frequencies are observed, which show constitutively activated BCR signals, but reduced or even absent responses upon BCR engagement (*3,117*).

Signaling pathway activation differed in maximum intensity and kinetics between children and adults. We observed that B cells in children have higher signaling induction upon anti-BCR stimulation, indicated by the higher ΔtMFI in all monitored phospho-proteins. In line with that, a 20-40% reduction in induction of kinases was previously shown in B cells from elderly donors (*118*). Additionally, the study of p38-induced inflammation measured in elderly adults (>65 years old) upon antigen challenge showed that p-p38 was significantly decreased in the aged donors compared to the younger ones (*119*). Our data also showed decreased phosphorylation of p38 in adults upon anti-BCR stimulation both in naïve and SM B cells. The PI3K/Akt/mTOR pathway activity has been shown to be significantly increased in elderly cohorts (*120,121*). This was further supported by the report of mTOR inhibition by rapamycin leading to decreased infection rates in the elderly (*122*), restoration of immune response to antigen (*123*), and improved responses to vaccination (*124*).

Careful selection of fluorochromes at the initial stages of panel design, sample handling, well-performed cryopreservation, and cell thawing are instrumental for assay performance. One of the limitations of the study is that due to spread between fluorochromes, it was not possible to reliably evaluate p-p65 expression in CD19^+^ B cells. Therefore, we suggest an improvement to the panel by moving the CD19 marker to RB744. Furthermore, the study is limited in assessing functionality of monocytes. This requires conducting the experiments on fresh whole blood as previously shown (*125–128*).

In summary, our newly established multiplex pFlow assay allows the assessment of complex signaling pathway activation, and its dynamics in response to six different stimuli. Multiplexing analysis enables capturing differences in signaling induction across different cell types and subpopulations. This comparative analysis can facilitate the choice of a specific cell population and/or stimulation to study a specific signaling pathway, for example in the frame of studying the effect of targeted therapies or to study the effect of a genetic mutation on defined signaling pathways. In addition, our assay may provide a tool to screen for lymphocytic and monocytic cell function that can be applied in disease settings to define disease activity or predict progression and prognosis. Age-specific differences in cellular signaling response provide the basis to understand differences in the immune response, such as memory formation and maintenance, and underline the importance of selecting matching controls for pediatric studies.

## Methods

### Donors

Peripheral blood (PB) samples were collected by venipuncture into EDTA coated vacuettes in Freiburg, Germany (Ethic approval of the Albert-Ludwig University of Freiburg 20-1109) and in the Motol University Hospital, Czechia (Ethic approval of Ethics Committee for Multi-Centric Clinical Trials of the University Hospital Motol and Second Faculty of Medicine, Charles University, Prague, EK-602.28/22). Samples from 54 healthy donors split into 3 age groups (<18 yo, 18-55 yo, ≥56 yo) were collected (Supplementary Table 1). All adult healthy donors signed an informed consent before PB donation, the consent for the minors was obtained from their parents or legal guardians, in line with the Declaration of Helsinki.

### PBMC isolation

PBMCs were isolated by density gradient centrifugation using Ficoll. The pellet was gently resuspended and aliquoted at no more than 20 million cells per mL of freezing medium (90 % fetal bovine serum (FBS) with 10 % DMSO) and stored in liquid nitrogen.

### Sample-quality criteria to perform pFlow

To perform the pFlow assay, samples needed to fulfill the following quality criteria. The minimum viability of thawed PBMC must be 70% of live cells. The minimum cell count to perform a complete pFlow must be 3 million live cells.

### pFlow assay

The method for analyzing the activation and kinetics of individual signaling pathways is based on detection of phosphorylated proteins representing the pathways of interest as described above. Antibodies for research use are available for detection of phosphorylated forms of the proteins.

Cryopreserved PBMCs were thawed in a water bath, and washed in Iscove’s Modified Dulbecco’s Medium (IMDM) (Gibco, USA) with 10 % FBS pre-warmed to 37 °C. After centrifugation, cells were counted using the Luna-FL Automated Fluorescence Cell Counter (BioCat, Germany). Cells were washed again and plated in a final concentration of 500,000 cells in 200 µL of IMDM+10% FBS. Cells were rested in an incubator at 37 °C, 6.5 % CO2 for 2 hours. Following, 150 µL of medium was removed and 5 µL of stimulation suspension was added.

Six different stimuli were used (Supplementary Table 1). The cells were stimulated at 3 different timepoints (5, 15, and 30 minutes). 20 minutes before the end of stimulation, a staining mix containing viability dye (Zombie NIR) and one antibody staining for surface chemokine receptor 7 (CCR7) was added to all wells. To stop the activation after the maximum timepoint, cells were fixed using the FoxP3 kit (eBioscience, Invitrogen, USA) at room temperature (RT) according to manufacturer’s instructions.

### Cytometry panel

After the cells were fixed, the plates were washed and barcoded based on the timepoints (unstimulated, and stimulated for 5, 15, and 30 minutes) for 30 minutes in the dark at RT, allowing for multiplexing of the assay (Figure 1A)(*77,129*). After barcoding, cells were washed and pooled based on the stimulus (thus pooling 4 wells into 1 well). Barcoded pooled cells were washed and stained with a freshly prepared surface staining premix (Supplementary Table 1). The cells were incubated for 30 minutes in the dark at RT.

After incubation the cells were washed and stained with a freshly prepared intracellular antibody mix resuspended in a permeabilization buffer (Supplementary Table 1). Cells were incubated for 30 minutes in the dark at RT. After the incubation, they were centrifuged twice. The cells were resuspended in 100 µl of antibody staining buffer.

### Full spectrum flow cytometry

Samples were acquired on Cytek Aurora with SpectroFlo (version 3.3.0, Cytek Biosciences, Fremont, CA, USA) and unmixing performed with the same reference controls from single-stained PBMCs from healthy donors, processed according to the protocol described above.

### Sample-quality criteria to proceed with analysis of pFlow

After de-barcoding, each sample was checked for cell count in individual cell populations. To include the sample into the healthy donor reference set, 70% of gated cell populations needed to contain a minimum of 50 cells. Next, major cell populations representing adaptive and innate immune cells, namely T cells, B cells, and NK cells were checked for response to the pan-activators, which served as positive control. Signal induction needed to be observed after pervanadate stimulation for p-BTK/p-ITK, p-Akt, and p-p38, and after PMA + Ionomycin for p-S6 and p-p65 NF-κB. If this criterion was not met, the samples were excluded from further analysis.

### Analysis

The unmixed FCS files were manually debarcoded and all cell populations were gated using FlowJo (v10.10.0, Beckman Coulter, USA) software. For the analysis of individual phospho-protein in all subpopulations, debarcoded FCS files and FlowJo workspaces were analyzed using R Statistical Software (v4.4.2)(*130*), and the following packages: data.table (*131*), dplyr (*132*), CytoML (*133*), flowCore (*134*), flowWorkspace (*135*), ggplot2 (*136*), ggdist (*137*), ggpubr (*138*), patchwork (*139*) and tidyr (*140*) using a custom pipeline. Individual cells were assigned a population identity based on imported gates and the data from phospho-protein markers were transformed using the inverse hyperbolic sine function (arcsinh) with cofactors selected based on the average magnitude of the signal in unstimulated samples (p-BTK/ITK: 3,000, p-Akt: 6,000, p-p38: 1,500, p-p65: 2,000, p-S6: 20,000). The median fluorescence intensity (MFI) was then calculated from the transformed data (tMFI) for each phospho-protein per timepoint, stimulus, and population. All subpopulations with 50 events or fewer were excluded from further analysis. ΔtMFI was calculated as the delta from the median of unstimulated values for each phospho-protein and population.

## Acknowledgements

The authors would like to thank all donors for their participation in the study, Jan Stuchly for his insight and support in the data analysis and the IR and Freeze Biobank Freiburg at the University Medical Center Freiburg. Figure 1A was created with BioRender.com

## Author contributions

**Petra Hadlova**: Conceptualization, Investigation, Data curation, Methodology, Visualization, Writing – original draft; **Michael Svaton**: Investigation, Data curation, Formal analysis, Software, Visualization, Writing – original draft; **Katerina Kochmannova**: Investigation, Validation; **Jakov Korzhenevich**: Methodology, Writing – review & editing; **Franziska Schmidt**: Methodology, Writing – review & editing; **Stefan F.H. Neys**: Methodology, Writing – review & editing; **Marei-Theresa Bott**: Investigation, Resources; **Petra Vrabcova**: Investigation, Resources; **Julian Staniek**: Methodology, Writing – review & editing; **Marketa Bloomfield**: Investigation, Resources, Data curation, Writing – review & editing; **Tomas Kalina**: Conceptualization, Supervision, Funding acquisition, Resources, Writing – original draft; **Marta Rizzi**: Conceptualization, Supervision, Funding acquisition, Resources, Writing – original draft

## Data availability statement

Underlying data for plots in Figures and Supplementary Figures can be found in Supplementary Table 2. The complete data are available from the corresponding author upon reasonable request.

---

**Supplementary Figure 1.**
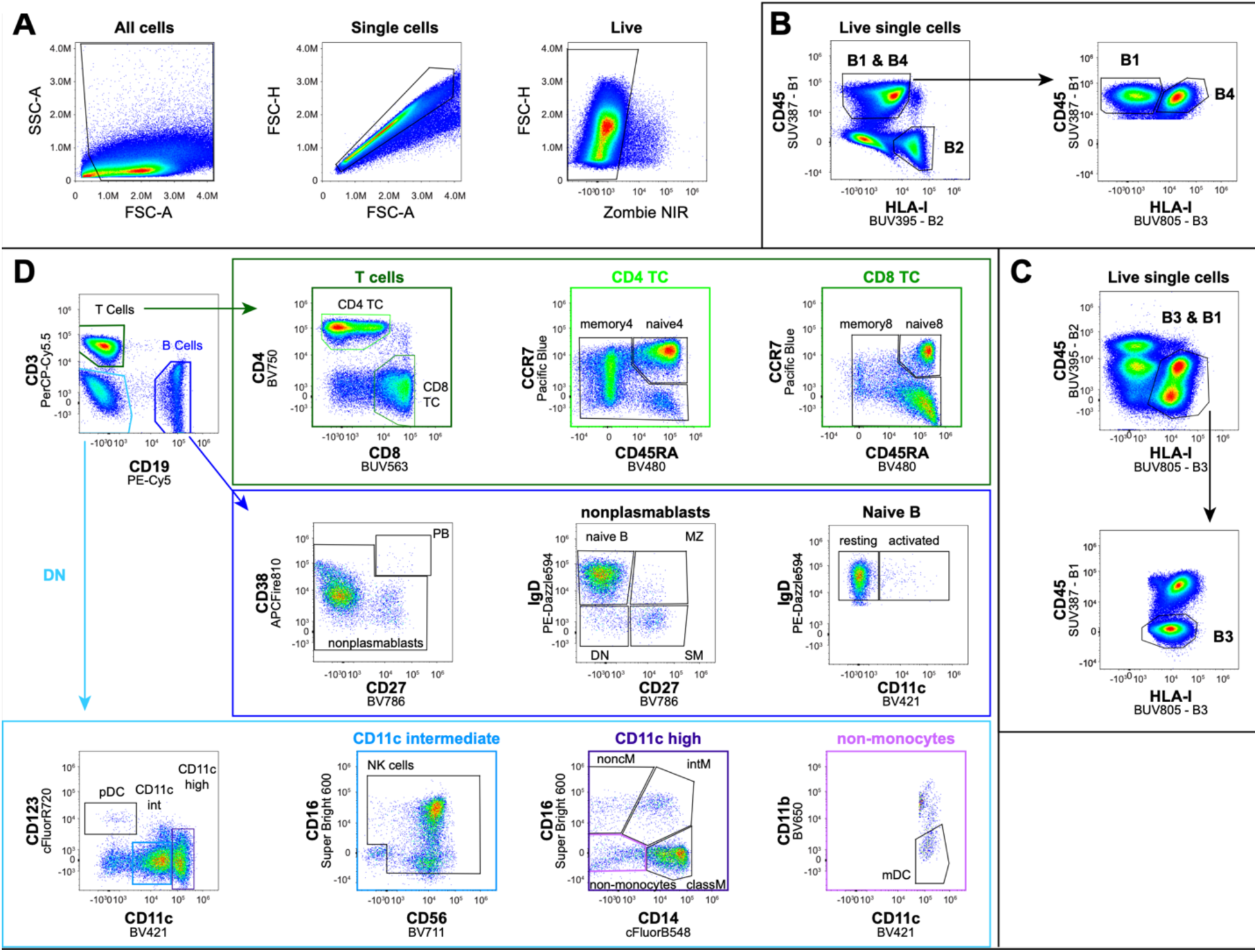
Gating strategy for pFlow analysis. **(A)** Gating strategy used for analysis in FlowJo. After elimination of debris by granularity and size, live single cells are debarcoded and analyzed further. Barcodes were assigned to each timepoint of stimulation, allowing the combining and simultaneous measuring of four timepoints in one well, B1 cells are single stained with CD45 in SUV387 and B4 cells are double stained with CD45 SUV387 and HLA-I in BUV805 **(B)**. The B2 cells are single stained with CD45 in BUV395 and B3 is single stained with HLA-I in BUV805 **(C)**. **(D)** Representative gating of individual cell populations from the B1 sample. First, the de-barcoded, live, single cells are divided into T cells (CD3^+^, CD19^-^), B cells (CD19^+^, CD3^-^) and innate immune cells (CD3^-^, CD19^-^, DN). T cells are subdivided into CD4^+^ T helper cells (CD4^+^CD8^-^) and CD8^+^ cytotoxic T cells (CD8^+^, CD4^-^). Each major subpopulation is further gated into naïve (CCR7^+^ CD45RA^+^) and memory cells (CCR7^+^ CD45RA^-^, CCR7^-^CD45RA^+^, CCR7^-^ CD45RA^-^). Within the B cells, plasmablasts (PB) are defined as CD38+CD27+. Non-plasmablasts are further divided into naïve B cells (IgD^+^ CD27^-^), marginal zone like B cells (MZB), IgD^+^ CD27^+^), switched memory B cells (SM, IgD^-^, CD27^+^) and double negative B cells (DNB, IgD^-^ CD27^-^). Innate immune cells (labeled DN) are further gated based on their expression of CD123 and CD11c. CD123^+^ CD11c^-^ are pDC, CD11c ^high^ are myeloid lineage cells. Within the cells with CD11c intermediate expression (CD11c ^inter^) CD56^-^CD16^-^ cells are excluded, while the rest are labeled and analyzed as NK cells. CD11c^high^ cells are subdivided into CD14^+^CD16^-^ classical monocytes (classM), CD14^+^CD16^+^ intermediate monocytes (intM) and CD14^-^CD16^+^ non-classical monocytes (noncM). Within the CD14^-^CD16^-^ non-monocytes, CD11c^+^CD11b^-^ cells are myeloid dendritic cells (mDCs).

**Supplementary Figure 2.**
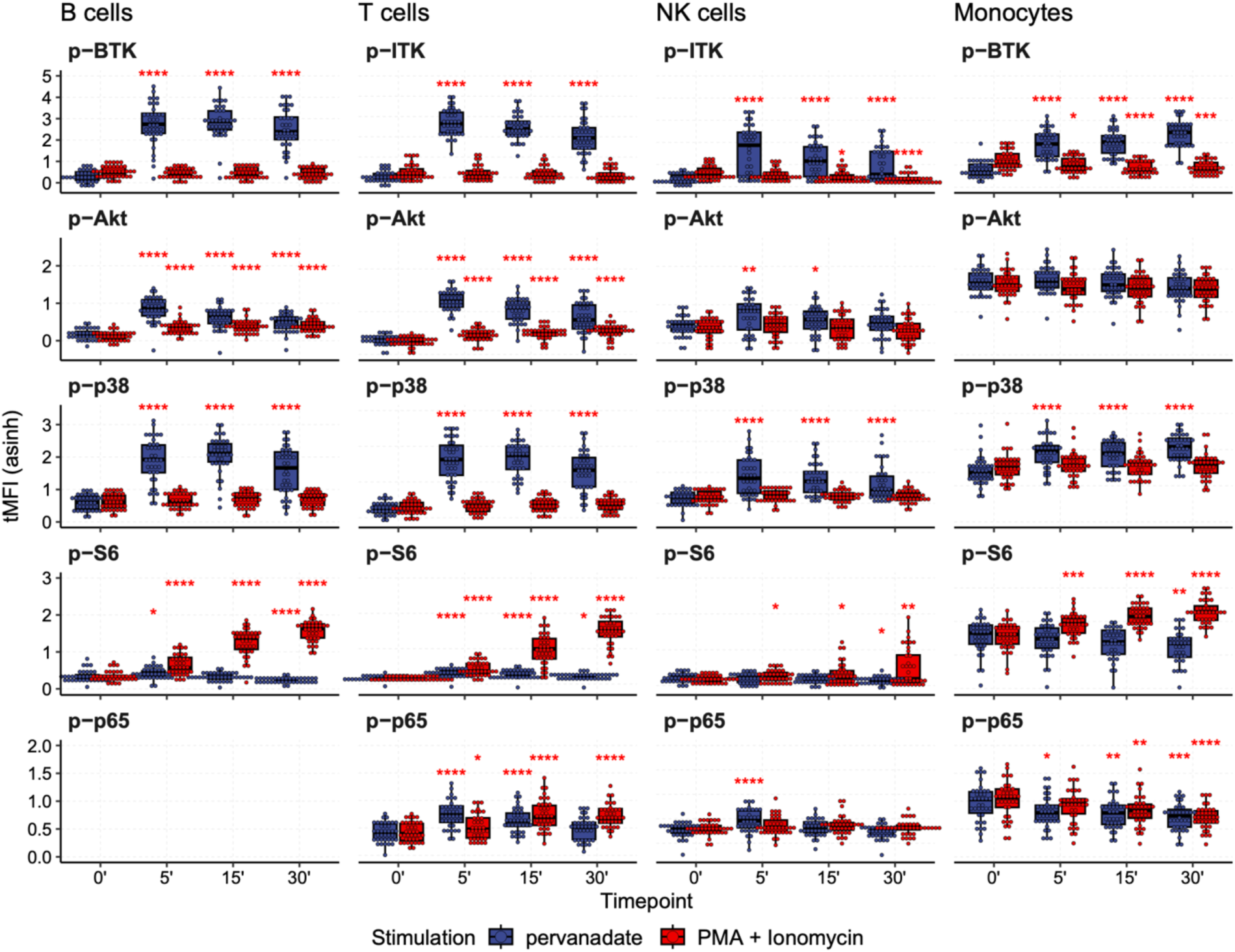
Response of major populations to pan-stimuli. Boxplots of transformed MFI values (tMFI, y-axis) for B, T and NK cells and monocytes in response to pan-stimuli, pervanadate (blue) and PMA + Ionomycin (red) at all timepoints (x-axis). Each cell population is depicted in one column. Each phospho-protein is depicted in one row. The statistical analysis was performed using an unpaired Welch’s t test between stimulated samples (5’,15’ and 30’) and the unstimulated (0’) with Benjamini-Hochberg correction. P-value is indicated by asterisks above graphs for significant observations only, P<0.05 (*), P<0.01 (**), P<0.001(***), P<0.0001 (****).

**Supplementary Figure 3.**
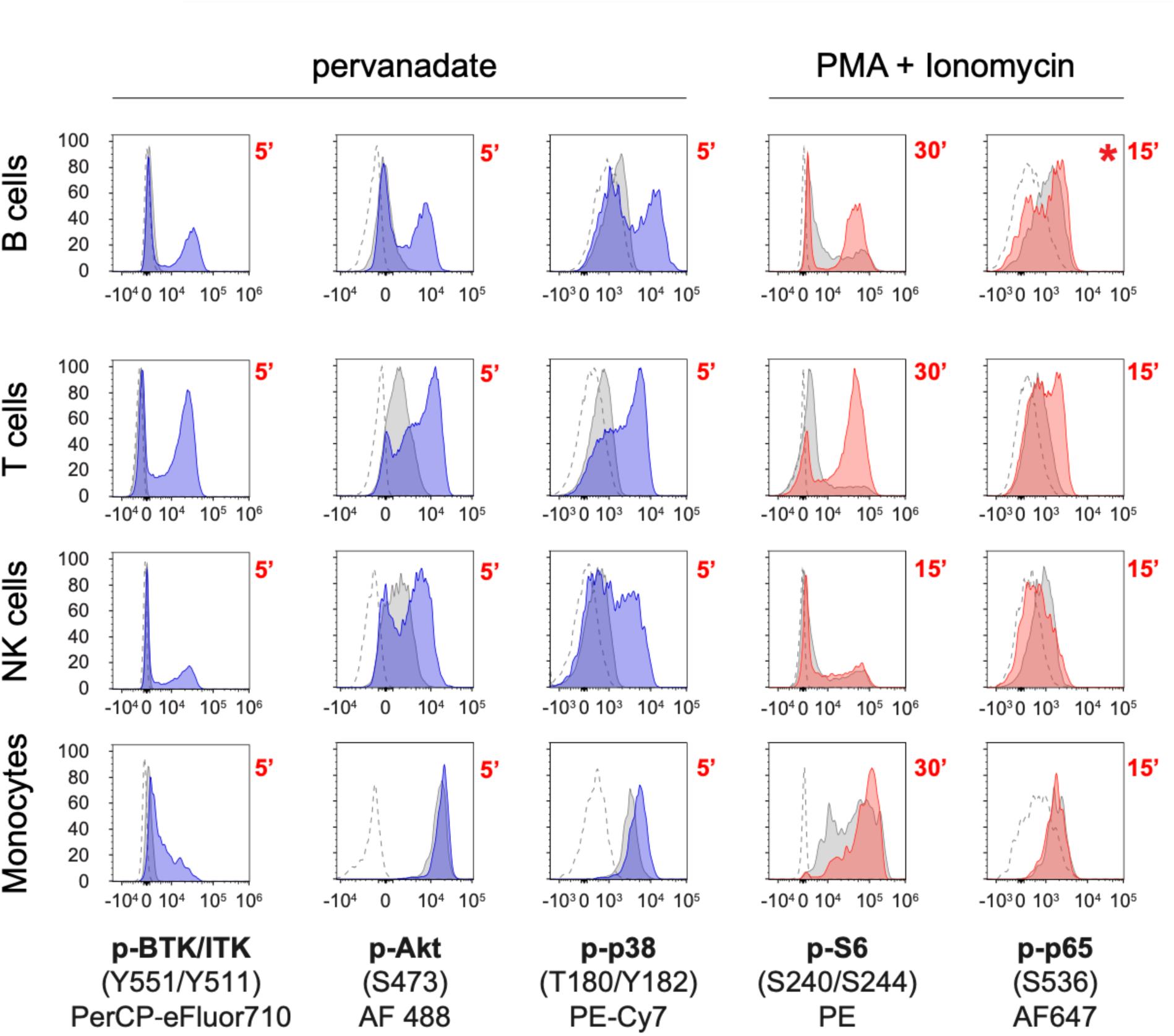
Illustrative raw data depiction of maximum induction across major cell populations. Raw data from one adult donor representative of the healthy cohort. Each row represents one cell population (indicated on the left) from top to bottom: B, T, NK cells and monocytes. Each column represents one phospho-protein (fluorescence intensity on x-axis). The proximal phospho-proteins, which are maximally induced by pervanadate, are depicted in blue (BTK/ITK, Akt, p38). The distal phospho-proteins, which are maximally induced by PMA + Ionomycin, are depicted in red (S6 and p65). The number in red indicates the duration of stimulation in minutes. Fluorochromes are indicated under the phospho-proteins below x-axis.

**Supplementary Figure 4.**
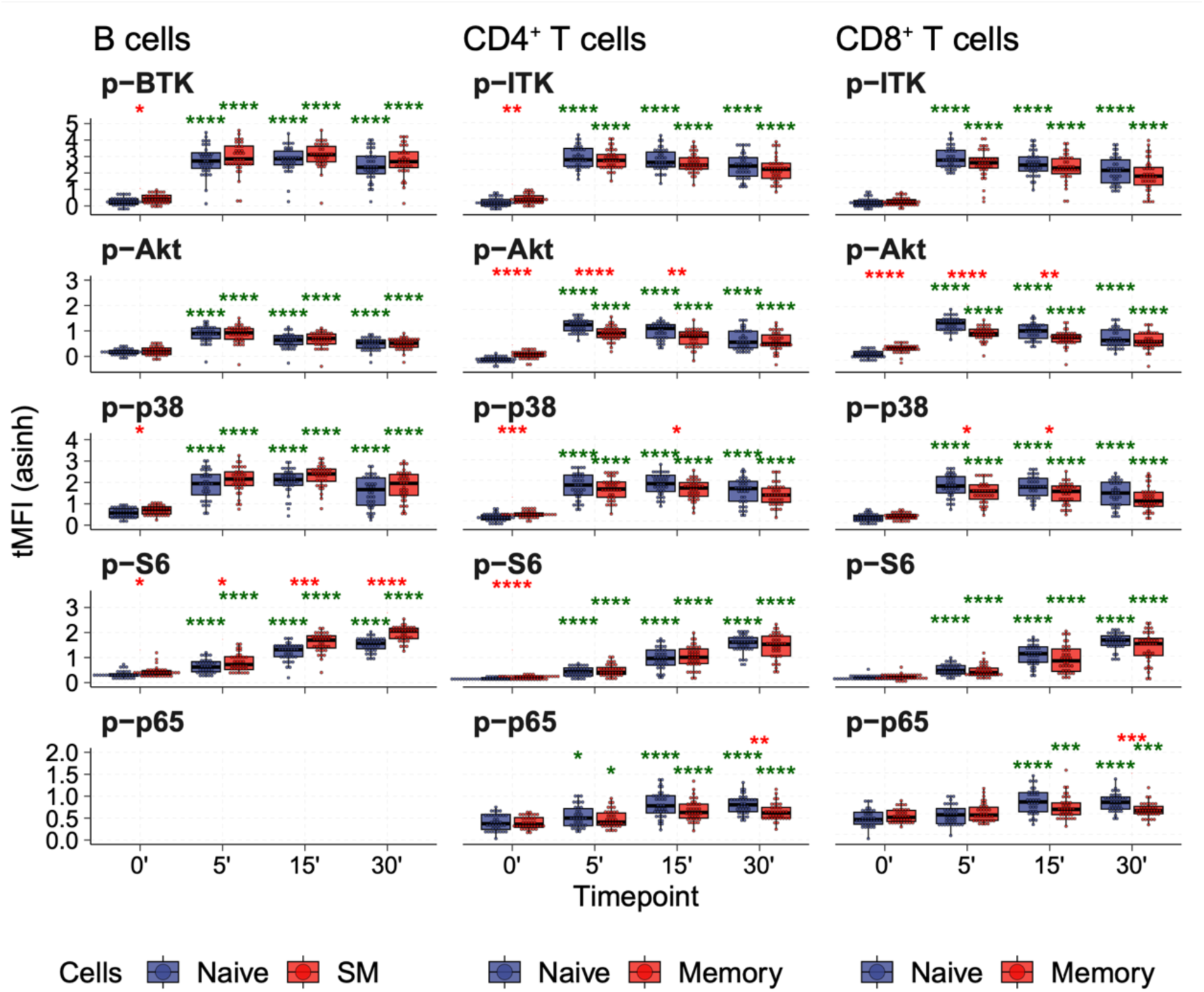
Response of subpopulations of B, CD4+ and CD8+T cells to pan-stimuli. Boxplots of transformed MFI values (tMFI, y-axis) for B cells, CD4^+^ T and CD8^+^ T cell subpopulations in response to pan-stimuli inducing the maximum signal for each respective phospho-protein as shown in previous figures (pervanadate for BTK/ITK, Akt and p38) and (PMA + Ionomycin for S6 and p65). Each phospho-protein is represented in one row. Each cell population (B cells, CD4^+^ T and CD8^+^ T cells) is depicted in one column and the subpopulations are distinguished by color naïve (blue) and memory T cells or switched memory (SM) B cells (red). The statistical analysis was performed using an unpaired Welch’s t test between between the selected subpopulations at respective timepoints (red asterisk) or between the stimulated (5’, 15’ and 30’) and the unstimulated (0’) samples (green asterisk).P-values were adjusted using Benjamini-Hochberg correction and only significant comparisons are indicated by asterisks above graphs, P<0.05 (*), P<0.01 (**), P<0.001(***), P<0.0001 (****).

**Supplementary Figure 5.**
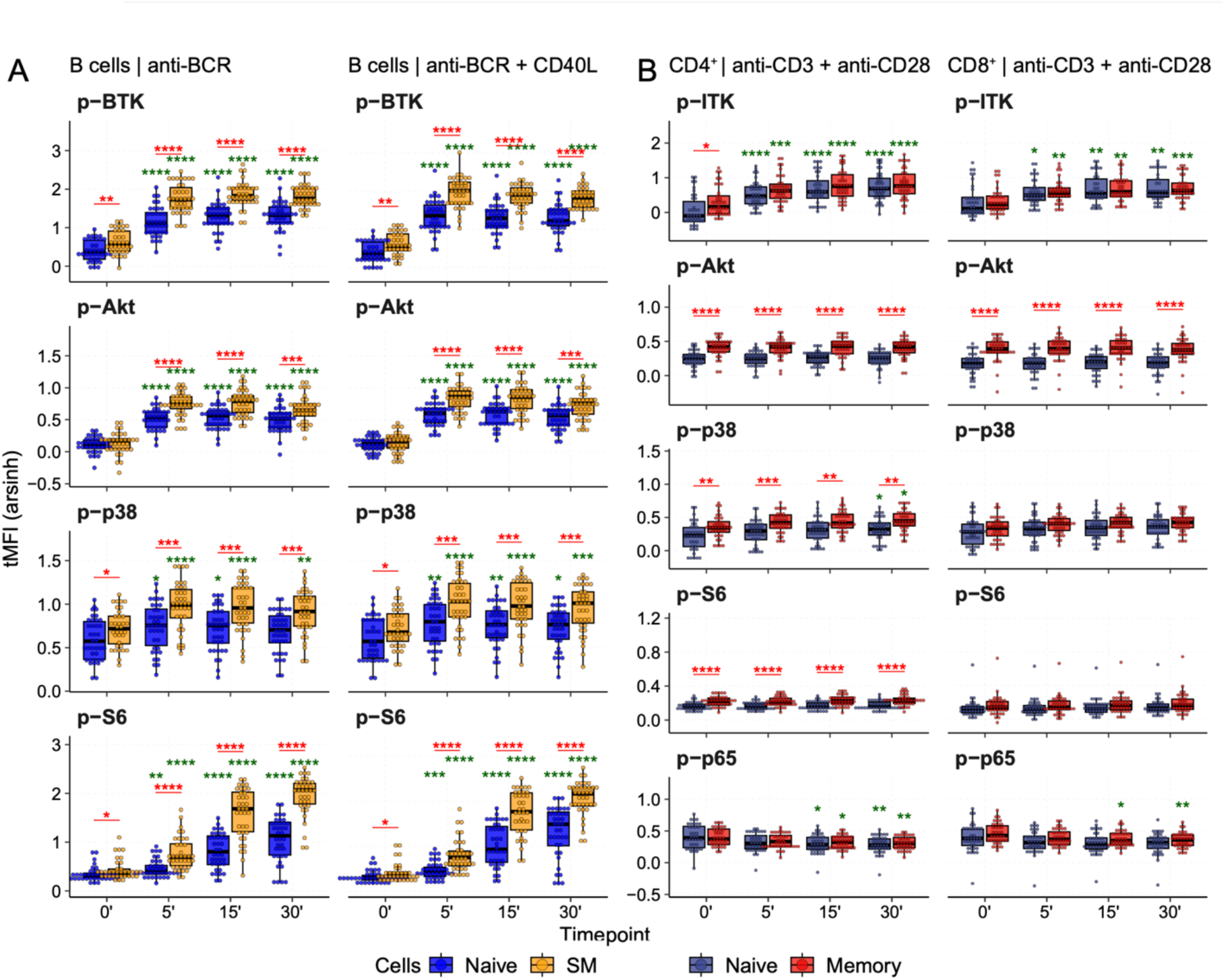
Response of subpopulations to targeted stimulation MFI. Boxplots of transformed MFI values (tMFI, y-axis) for B cell, CD4^+^ T and CD8^+^ T cell subpopulations in response to targeted stimulation. Each phospho-protein is represented in one row. Each cell population (B cells, CD4^+^ T and CD8^+^ T cells) stimulated with a targeted stimulation is depicted in one column and the subpopulations are distinguished by color. (A) B cell subpopulations naïve (blue) and switched memory (yellow) B cells stimulated with anti-BCR (left) and anti-BCR+CD40L (right). (B) From left to right naïve CD4+ T cells (blue) and memory CD4+ T cells (red) and naïve CD8+ T cells (blue) and memory CD8+ T cells (red) stimulated with anti-CD3 + anti-CD28. The statistical analysis was performed using an unpaired Welch’s t test between the selected subpopulations at respective timepoints (red asterisk) or between the stimulated (5’, 15’ and 30’) and the unstimulated (0’) samples (green asterisk).P-values were adjusted using Benjamini-Hochberg correction and only significant comparisons are indicated by asterisks above graphs, P<0.05 (*), P<0.01 (**), P<0.001(***), P<0.0001 (****).

**Supplementary Table 1A:** BARCODES.

| **marker** | **fluorochrome** | **clone** | **company** | **catalogue number** | **dilution 1 to** |
| --- | --- | --- | --- | --- | --- |
| CD45 | SUV387 | HI30 | BD Biosciences | 563792 | 200 |
| CD45 | BUV395 | HI30 | BioLegend | B405046 | 1000 |
| HLA-A,B,C | BUV805 | W6/32 | BD Biosciences | 752788 | 500 |

**Supplementary Table 1B:** SURFACE STAINING.

| **marker** | **fluorochrome** | **clone** | **company** | **catalogue number** | **dilution 1 to** |
| --- | --- | --- | --- | --- | --- |
| CD8a | BUV563 | RPA-T8 | BD Biosciences | 612914 | 1000 |
| CD20 | BUV737 | 2H7 | Invitrogen | 367-0209-42 | 200 |
| CD11c | BV421 | S-HCL-3 | BioLegend | B357126 | 300 |
| CD197 | Pacific Blue | G043H7 | BioLegend | 353210 | 200 |
| CD45RA | BV480 | HI100 | BD Biosciences | 566114 | 1500 |
| CD16 | Super Bright 600 | CB16 | Invitrogen | 63-0168-42 | 100 |
| CD11b | BV650 | M1/70 | BioLegend | 101239 | 100 |
| CD56 | BV711 | HCD56 | BioLegend | 318336 | 400 |
| CD4 | BV750 | SK3 | BD Biosciences | 566356 | 400 |
| CD27 | BV786 | L128 | BD Biosciences | 563327 | 100 |
| CD14 | cFluor B548 | 63D3 | Cytek | R7-20115 | 800 |
| CD3 | PerCP-Cy5.5 | UCHT1 | Tonbo biosciences | 65-0038-T100 | 200 |
| IgD | PE-Dazzle 594 | IA6-2 | BioLegend | 348240 | 400 |
| CD19 | PE-Cy5 | HIB19 | Tonbo biosciences | 55-0199-T100 | 200 |
| CD123 | cFluor R720 | 6H6 | Cytek | R7-20013 | 400 |
| CD38 | APC-Fire 810 | HB-7 | BioLegend | 356644 | 100 |
| Zombie | Near Infrared (NIR) |  | BioLegend | 423106 | 800 |

**Supplementary Table 1C:**
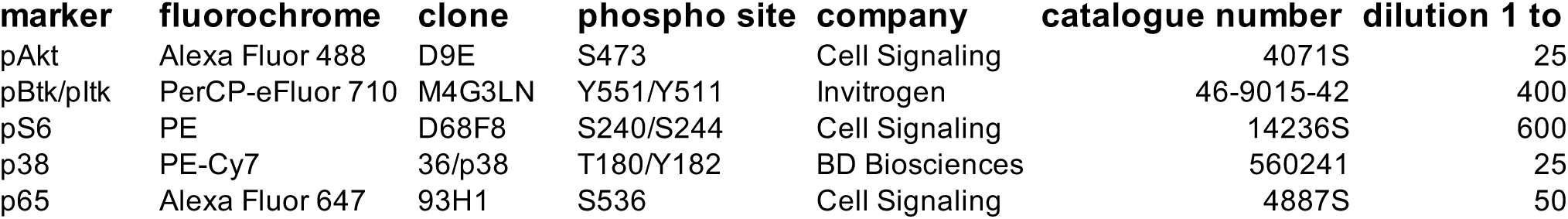
INTRACELLULAR STAINING.

| marker | fluorochrome | clone | phospho site | company | catalogue number | dilution 1 to |
| --- | --- | --- | --- | --- | --- | --- |
| pAkt | Alexa Fluor 488 | D9E | S473 | Cell Signaling | 4071S | 25 |
| pBtk/pltk | PerCP-eFluor 710 | M4G3LN | Y551/Y511 | Invitrogen | 46-9015-42 | 400 |
| pS6 | PE | D68F8 | S240/S244 | Cell Signaling | 14236S | 600 |
| p38 | PE-Cy7 | 36/p38 | T180/Y182 | BD Biosciences | 560241 | 25 |
| p65 | Alexa Fluor 647 | 93H1 | S536 | Cell Signaling | 4887S | 50 |

**Supplementary Table 1D:** REAGENTS.

| **Reagent** | **company** | **catalogue number** |
| --- | --- | --- |
| eBioscience FOXP3/Transcription factor staining buffer set | Invitrogen, eBioscience | 00-5523-00 |
| *component 1.* Permeabilization Buffer 10x | Invitrogen, eBioscience | 00-8333-56 |
| *component 2.* Fixation/Permeabilization concentrate | Invitrogen, eBioscience | 00-5123-43 |
| *component 3.* Fixation/Perm diluent | Invitrogen, eBioscience | 00-5223-56 |
| Brilliant Stain Buffer Plus | BD Biosciences | 566385 |
| FBS Superior | Sigma | S0615-500ML |
| Iscove' s Modified Dulbecco' s Medium (IMDM 1x) | Gibco | 21980-032 |
| Penicilin Streptomycin | Gibco | 15140-122 |
| Acridin Orange/ Propidium Iodide Stain Cell Viability Kit | BioCat | F23001-LG |

**Supplementary Table 1E:** STIMULI.

| Stimulus | company | catalogue number | final concentration |
| --- | --- | --- | --- |
| Fab' 2 anti Ig A, G, M | Jackson ImmunoResearch | 109-006-064 | 9.23ug/mL |
| anti CD3 (clone OKT3) | BioLegend | 302934 | 1,25 µg/ml |
| anti CD28 (clone CD28.2) | BioLegend | 317326 | 1,25 µg/ml |
| cell stimulation cocktail (w/o protein transport inhibitors) 500x | Tonbo Biosciences | TNB-4970-UL 100 | 1x |
| LPS | Sigma | L4005 | 100 ng/ml |
| orthovanadate | Sigma | S6508-10G |  |
| CD40L | AG Rizzi, produced in laboratory |  |  |

**Supplementary Table 1F:** SAMPLES.

| **sample_code** | **status** | **sex** | **age** |
| --- | --- | --- | --- |
| HD_001 | HD | F | 25.1 |
| HD_002 | HD | F | 30 |
| HD_003 | HD | M | 27 |
| HD_004 | HD | F | 23.5 |
| HD_005 | HD | M | 24.8 |
| HD_006 | HD | F | 24.9 |
| HD_007 | HD | F | 24.5 |
| HD_008 | HD | F | 22.8 |
| HD_009 | HD | F | 38 |
| HD_010 | HD | F | 38.9 |
| HD_011 | HD | F | 22.5 |
| HD_012 | HD | F | 30.2 |
| HD_013 | HD | F | 22.2 |
| HD_014 | HD | F | 54.25 |
| HD_015 | HD | F | 26.25 |
| HD_016 | HD | F | 53.2 |
| HD_017 | HD | F | 56.2 |
| HD_018 | HD | F | 61.2 |
| HD_019 | HD | F | 60.2 |
| HD_020 | HD | F | 61.2 |
| HD_021 | HD | F | 32 |
| HD_022 | HD | F | 23 |
| HD_023 | HD | M | 10 |
| HD_024 | HD | F | 38 |
| HD_025 | HD | M | 1 |
| HD_026 | HD | M | 23 |
| HD_027 | HD | M | 3 |
| HD_028 | HD | M | 33 |
| HD_029 | HD | F | 39 |
| HD_030 | HD | F | 53.2 |
| HD_031 | HD | M | 6 |
| HD_032 | HD | M | 11 |
| HD_033 | HD | M | 11 |
| HD_034 | HD | M | 7 |
| HD_035 | HD | M | 38 |
| HD_036 | HD | M | 25 |
| HD_037 | HD | F | 14 |
| HD_038 | HD | M | 46 |
| HD_039 | HD | M | 43 |
| HD_040 | HD | M | 57 |
| HD_041 | HD | M | 25 |
| HD_042 | HD | M | 33 |
| HD_043 | HD | M | 31 |
| HD_044 | HD | M | 28 |
| HD_045 | HD | M | 30 |
| HD_046 | HD | M | 7 |
| HD_047 | HD | M | 16 |
| HD_048 | HD | F | 69 |
| HD_049 | HD | M | 31 |
| HD_050 | HD | M | 22 |
| HD_051 | HD | M | 22 |
| HD_052 | HD | M | 28 |
| HD_053 | HD | F | 21 |
| HD_054 | HD | F | 20 |

**Supplementary Table 2A:** FIGURE 1B_DATA.

| cells | marker | Stimulation | Timing | Count | Mean | SEM | Q25 | Q75 |
| --- | --- | --- | --- | --- | --- | --- | --- | --- |
| CD19BC | p-Btk | pV | unstim | 38 | 0 | 0 | 0 | 0 |
| CD19BC | p-Btk | pV | 5min | 38 | 2.273574 | 0.145285 | 2.00653 | 2.914537 |
| CD19BC | p-Btk | pV | 15min | 38 | 2.373596 | 0.11088 | 1.923358 | 2.855928 |
| CD19BC | p-Btk | pV | 30min | 38 | 2.024052 | 0.13213 | 1.464026 | 2.661647 |
| CD19BC | p-Btk | PMA_I | unstim | 38 | 0 | 0 | 0 | 0 |
| CD19BC | p-Btk | PMA_I | 5min | 37 | -0.01649 | 0.012343 | -0.06358 | 0.039356 |
| CD19BC | p-Btk | PMA_I | 15min | 38 | -0.02283 | 0.012062 | -0.07846 | 0.025278 |
| CD19BC | p-Btk | PMA_I | 30min | 38 | -0.0472 | 0.012291 | -0.08667 | -0.00031 |
| CD19BC | p-Akt | pV | unstim | 38 | 0 | 0 | 0 | 0 |
| CD19BC | p-Akt | pV | 5min | 38 | 0.718068 | 0.0483 | 0.500965 | 0.906914 |
| CD19BC | p-Akt | pV | 15min | 38 | 0.490917 | 0.043612 | 0.296867 | 0.664958 |
| CD19BC | p-Akt | pV | 30min | 38 | 0.348372 | 0.029859 | 0.216094 | 0.48116 |
| CD19BC | p-Akt | PMA_I | unstim | 38 | 0 | 0 | 0 | 0 |
| CD19BC | p-Akt | PMA_I | 5min | 37 | 0.211495 | 0.021138 | 0.136556 | 0.270015 |
| CD19BC | p-Akt | PMA_I | 15min | 38 | 0.250941 | 0.019289 | 0.196823 | 0.304246 |
| CD19BC | p-Akt | PMA_I | 30min | 38 | 0.270189 | 0.016668 | 0.21863 | 0.314068 |
| CD19BC | p-p38 | pV | unstim | 38 | 0 | 0 | 0 | 0 |
| CD19BC | p-p38 | pV | 5min | 38 | 1.258136 | 0.117883 | 0.819228 | 1.768712 |
| CD19BC | p-p38 | pV | 15min | 38 | 1.426109 | 0.090671 | 1.169827 | 1.807279 |
| CD19BC | p-p38 | pV | 30min | 38 | 0.947872 | 0.117058 | 0.254308 | 1.524706 |
| CD19BC | p-p38 | PMA_I | unstim | 38 | 0 | 0 | 0 | 0 |
| CD19BC | p-p38 | PMA_I | 5min | 37 | 0.036513 | 0.013709 | -0.02911 | 0.095405 |
| CD19BC | p-p38 | PMA_I | 15min | 38 | 0.060638 | 0.01592 | -0.01519 | 0.092707 |
| CD19BC | p-p38 | PMA_I | 30min | 38 | 0.060618 | 0.016857 | -0.01459 | 0.086032 |
| CD19BC | p-S6 | pV | unstim | 38 | 0 | 0 | 0 | 0 |
| CD19BC | p-S6 | pV | 5min | 38 | 0.072328 | 0.017978 | 0.044414 | 0.131522 |
| CD19BC | p-S6 | pV | 15min | 38 | -0.02029 | 0.018579 | -0.04808 | 0.047562 |
| CD19BC | p-S6 | pV | 30min | 38 | -0.11805 | 0.021195 | -0.15264 | -0.03074 |
| CD19BC | p-S6 | PMA_I | unstim | 38 | 0 | 0 | 0 | 0 |
| CD19BC | p-S6 | PMA_I | 5min | 37 | 0.303564 | 0.033238 | 0.127362 | 0.448741 |
| CD19BC | p-S6 | PMA_I | 15min | 38 | 0.920045 | 0.052097 | 0.764278 | 1.170388 |
| CD19BC | p-S6 | PMA_I | 30min | 38 | 1.219392 | 0.040854 | 1.045915 | 1.459723 |
| CD19BC | p-p65 | pV | unstim | 0 | NA | NA | NA | NA |
| CD19BC | p-p65 | pV | 5min | 0 | NA | NA | NA | NA |
| CD19BC | p-p65 | pV | 15min | 0 | NA | NA | NA | NA |
| CD19BC | p-p65 | pV | 30min | 0 | NA | NA | NA | NA |
| CD19BC | p-p65 | PMA_I | unstim | 0 | NA | NA | NA | NA |
| CD19BC | p-p65 | PMA_I | 5min | 0 | NA | NA | NA | NA |
| CD19BC | p-p65 | PMA_I | 15min | 0 | NA | NA | NA | NA |
| CD19BC | p-p65 | PMA_I | 30min | 0 | NA | NA | NA | NA |
| CD3TC | p-Btk | pV | unstim | 38 | 0 | 0 | 0 | 0 |
| CD3TC | p-Btk | pV | 5min | 38 | 2.498522 | 0.090224 | 2.073547 | 2.825223 |
| CD3TC | p-Btk | pV | 15min | 38 | 2.270216 | 0.100033 | 2.01092 | 2.704716 |
| CD3TC | p-Btk | pV | 30min | 38 | 1.794085 | 0.105975 | 1.391697 | 2.197719 |
| CD3TC | p-Btk | PMA_I | unstim | 38 | 0 | 0 | 0 | 0 |
| CD3TC | p-Btk | PMA_I | 5min | 37 | 0.018196 | 0.01679 | -0.05553 | 0.079121 |
| CD3TC | p-Btk | PMA_I | 15min | 38 | -0.02574 | 0.017772 | -0.06689 | 0.036232 |
| CD3TC | p-Btk | PMA_I | 30min | 38 | -0.08131 | 0.01696 | -0.12052 | -0.01491 |
| CD3TC | p-Akt | pV | unstim | 38 | 0 | 0 | 0 | 0 |
| CD3TC | p-Akt | pV | 5min | 38 | 0.942795 | 0.035019 | 0.818472 | 1.060625 |
| CD3TC | p-Akt | pV | 15min | 38 | 0.742582 | 0.038482 | 0.589674 | 0.926197 |
| CD3TC | p-Akt | pV | 30min | 38 | 0.559411 | 0.046902 | 0.336883 | 0.777193 |
| CD3TC | p-Akt | PMA_I | unstim | 38 | 0 | 0 | 0 | 0 |
| CD3TC | p-Akt | PMA_I | 5min | 37 | 0.158495 | 0.01343 | 0.111932 | 0.176974 |
| CD3TC | p-Akt | PMA_I | 15min | 38 | 0.195249 | 0.013332 | 0.139175 | 0.24268 |
| CD3TC | p-Akt | PMA_I | 30min | 38 | 0.242709 | 0.014667 | 0.16969 | 0.294515 |
| CD3TC | p-p38 | pV | unstim | 38 | 0 | 0 | 0 | 0 |
| CD3TC | p-p38 | pV | 5min | 38 | 1.34711 | 0.092335 | 0.977142 | 1.706193 |
| CD3TC | p-p38 | pV | 15min | 38 | 1.36031 | 0.061604 | 1.04682 | 1.656151 |
| CD3TC | p-p38 | pV | 30min | 38 | 1.027791 | 0.082634 | 0.693565 | 1.333588 |
| CD3TC | p-p38 | PMA_I | unstim | 38 | 0 | 0 | 0 | 0 |
| CD3TC | p-p38 | PMA_I | 5min | 37 | 0.068636 | 0.011622 | 0.020606 | 0.11263 |
| CD3TC | p-p38 | PMA_I | 15min | 38 | 0.097422 | 0.011798 | 0.053843 | 0.137947 |
| CD3TC | p-p38 | PMA_I | 30min | 38 | 0.093533 | 0.012301 | 0.04247 | 0.129849 |
| CD3TC | p-S6 | pV | unstim | 38 | 0 | 0 | 0 | 0 |
| CD3TC | p-S6 | pV | 5min | 38 | 0.146036 | 0.012379 | 0.117688 | 0.183402 |
| CD3TC | p-S6 | pV | 15min | 38 | 0.11769 | 0.01477 | 0.069206 | 0.190624 |
| CD3TC | p-S6 | pV | 30min | 38 | 0.045632 | 0.009561 | 0.020437 | 0.080949 |
| CD3TC | p-S6 | PMA_I | unstim | 38 | 0 | 0 | 0 | 0 |
| CD3TC | p-S6 | PMA_I | 5min | 37 | 0.267614 | 0.029775 | 0.138737 | 0.414897 |
| CD3TC | p-S6 | PMA_I | 15min | 38 | 0.803008 | 0.064721 | 0.584409 | 1.063206 |
| CD3TC | p-S6 | PMA_I | 30min | 38 | 1.335535 | 0.057699 | 1.182661 | 1.61345 |
| CD3TC | p-p65 | pV | unstim | 38 | 0 | 0 | 0 | 0 |
| CD3TC | p-p65 | pV | 5min | 38 | 0.318524 | 0.019647 | 0.230402 | 0.400815 |
| CD3TC | p-p65 | pV | 15min | 38 | 0.219537 | 0.016755 | 0.15396 | 0.294157 |
| CD3TC | p-p65 | pV | 30min | 38 | 0.064736 | 0.016663 | -0.00597 | 0.161726 |
| CD3TC | p-p65 | PMA_I | unstim | 38 | 0 | 0 | 0 | 0 |
| CD3TC | p-p65 | PMA_I | 5min | 37 | 0.090147 | 0.017657 | 0.016103 | 0.163597 |
| CD3TC | p-p65 | PMA_I | 15min | 38 | 0.31424 | 0.02707 | 0.215627 | 0.438557 |
| CD3TC | p-p65 | PMA_I | 30min | 38 | 0.292865 | 0.018537 | 0.210526 | 0.36784 |
| NK | p-Btk | pV | unstim | 38 | 0 | 0 | 0 | 0 |
| NK | p-Btk | pV | 5min | 38 | 1.069402 | 0.153768 | 0.035867 | 1.852495 |
| NK | p-Btk | pV | 15min | 38 | 0.674569 | 0.099311 | 0.070122 | 1.230438 |
| NK | p-Btk | pV | 30min | 38 | 0.442357 | 0.092558 | 0.011086 | 0.988805 |
| NK | p-Btk | PMA_I | unstim | 38 | 0 | 0 | 0 | 0 |
| NK | p-Btk | PMA_I | 5min | 37 | -0.01674 | 0.01851 | -0.03768 | 0.029794 |
| NK | p-Btk | PMA_I | 15min | 38 | -0.09304 | 0.020974 | -0.15996 | -0.0077 |
| NK | p-Btk | PMA_I | 30min | 38 | -0.18965 | 0.027615 | -0.26763 | -0.05403 |
| NK | p-Akt | pV | unstim | 38 | 0 | 0 | 0 | 0 |
| NK | p-Akt | pV | 5min | 38 | 0.34266 | 0.044937 | 0.105824 | 0.543911 |
| NK | p-Akt | pV | 15min | 38 | 0.200874 | 0.027701 | 0.039406 | 0.34868 |
| NK | p-Akt | pV | 30min | 38 | 0.125519 | 0.02326 | 0.049903 | 0.194071 |
| NK | p-Akt | PMA_I | unstim | 38 | 0 | 0 | 0 | 0 |
| NK | p-Akt | PMA_I | 5min | 37 | 0.062521 | 0.016343 | -0.01183 | 0.106965 |
| NK | p-Akt | PMA_I | 15min | 38 | 0.009819 | 0.019746 | -0.06338 | 0.071609 |
| NK | p-Akt | PMA_I | 30min | 38 | -0.06293 | 0.018803 | -0.11167 | -0.01597 |
| NK | p-p38 | pV | unstim | 38 | 0 | 0 | 0 | 0 |
| NK | p-p38 | pV | 5min | 38 | 0.691974 | 0.10025 | 0.102641 | 1.212164 |
| NK | p-p38 | pV | 15min | 38 | 0.593836 | 0.085822 | 0.134931 | 0.886302 |
| NK | p-p38 | pV | 30min | 38 | 0.442797 | 0.083056 | 0.064704 | 0.712068 |
| NK | p-p38 | PMA_I | unstim | 38 | 0 | 0 | 0 | 0 |
| NK | p-p38 | PMA_I | 5min | 37 | 0.059173 | 0.013798 | 0.005626 | 0.081454 |
| NK | p-p38 | PMA_I | 15min | 38 | 0.055458 | 0.016393 | 0.000598 | 0.10105 |
| NK | p-p38 | PMA_I | 30min | 38 | 0.029757 | 0.015429 | -0.04932 | 0.103789 |
| NK | p-S6 | pV | unstim | 38 | 0 | 0 | 0 | 0 |
| NK | p-S6 | pV | 5min | 38 | 0.006107 | 0.010965 | -0.01992 | 0.055098 |
| NK | p-S6 | pV | 15min | 38 | -0.0198 | 0.011348 | -0.04106 | 0.028666 |
| NK | p-S6 | pV | 30min | 38 | -0.05552 | 0.012174 | -0.09193 | 0.003509 |
| NK | p-S6 | PMA_I | unstim | 38 | 0 | 0 | 0 | 0 |
| NK | p-S6 | PMA_I | 5min | 37 | 0.062549 | 0.015378 | -0.00343 | 0.106679 |
| NK | p-S6 | PMA_I | 15min | 38 | 0.117784 | 0.03982 | -0.02413 | 0.182248 |
| NK | p-S6 | PMA_I | 30min | 38 | 0.289059 | 0.076433 | -0.03189 | 0.502173 |
| NK | p-p65 | pV | unstim | 38 | 0 | 0 | 0 | 0 |
| NK | p-p65 | pV | 5min | 38 | 0.160122 | 0.030575 | 0.005973 | 0.33795 |
| NK | p-p65 | pV | 15min | 38 | 0.01949 | 0.023764 | -0.07818 | 0.133574 |
| NK | p-p65 | pV | 30min | 38 | -0.05932 | 0.015566 | -0.10887 | 0.008201 |
| NK | p-p65 | PMA_I | unstim | 38 | 0 | 0 | 0 | 0 |
| NK | p-p65 | PMA_I | 5min | 37 | 0.059656 | 0.022384 | -0.0168 | 0.108775 |
| NK | p-p65 | PMA_I | 15min | 38 | 0.066986 | 0.02582 | -0.03184 | 0.120929 |
| NK | p-p65 | PMA_I | 30min | 38 | 0.017416 | 0.019782 | -0.02658 | 0.077731 |
| classM | p-Btk | pV | unstim | 38 | 0 | 0 | 0 | 0 |
| classM | p-Btk | pV | 5min | 37 | 0.813314 | 0.090094 | 0.456009 | 1.27836 |
| classM | p-Btk | pV | 15min | 37 | 0.883714 | 0.079487 | 0.500585 | 1.28654 |
| classM | p-Btk | pV | 30min | 38 | 1.273454 | 0.085426 | 0.919865 | 1.642775 |
| classM | p-Btk | PMA_I | unstim | 38 | 0 | 0 | 0 | 0 |
| classM | p-Btk | PMA_I | 5min | 37 | -0.13513 | 0.027908 | -0.28519 | -0.01915 |
| classM | p-Btk | PMA_I | 15min | 38 | -0.27178 | 0.034558 | -0.4338 | -0.10914 |
| classM | p-Btk | PMA_I | 30min | 38 | -0.24534 | 0.039678 | -0.39184 | -0.05089 |
| classM | p-Akt | pV | unstim | 38 | 0 | 0 | 0 | 0 |
| classM | p-Akt | pV | 5min | 37 | 0.144001 | 0.021294 | 0.091069 | 0.209456 |
| classM | p-Akt | pV | 15min | 37 | 0.05121 | 0.025296 | -0.06532 | 0.162096 |
| classM | p-Akt | pV | 30min | 38 | -0.05926 | 0.029654 | -0.18398 | 0.056161 |
| classM | p-Akt | PMA_I | unstim | 38 | 0 | 0 | 0 | 0 |
| classM | p-Akt | PMA_I | 5min | 37 | -0.05759 | 0.018775 | -0.13879 | 0.014668 |
| classM | p-Akt | PMA_I | 15min | 38 | -0.07747 | 0.020625 | -0.13566 | -0.01093 |
| classM | p-Akt | PMA_I | 30min | 38 | -0.108 | 0.022159 | -0.18486 | -0.00731 |
| classM | p-p38 | pV | unstim | 38 | 0 | 0 | 0 | 0 |
| classM | p-p38 | pV | 5min | 37 | 0.491679 | 0.059674 | 0.275 | 0.766019 |
| classM | p-p38 | pV | 15min | 37 | 0.447495 | 0.056609 | 0.184734 | 0.717712 |
| classM | p-p38 | pV | 30min | 38 | 0.601708 | 0.056377 | 0.307567 | 0.875176 |
| classM | p-p38 | PMA_I | unstim | 38 | 0 | 0 | 0 | 0 |
| classM | p-p38 | PMA_I | 5min | 37 | 0.118709 | 0.025438 | 0.047332 | 0.204914 |
| classM | p-p38 | PMA_I | 15min | 38 | 0.085171 | 0.028704 | -0.02632 | 0.185941 |
| classM | p-p38 | PMA_I | 30min | 38 | 0.092217 | 0.026726 | -0.01962 | 0.194024 |
| classM | p-S6 | pV | unstim | 38 | 0 | 0 | 0 | 0 |
| classM | p-S6 | pV | 5min | 37 | -0.14815 | 0.049106 | -0.22331 | -0.02215 |
| classM | p-S6 | pV | 15min | 37 | -0.25373 | 0.05351 | -0.40527 | -0.03455 |
| classM | p-S6 | pV | 30min | 38 | -0.35305 | 0.056652 | -0.52095 | -0.13279 |
| classM | p-S6 | PMA_I | unstim | 38 | 0 | 0 | 0 | 0 |
| classM | p-S6 | PMA_I | 5min | 37 | 0.313141 | 0.053958 | 0.069661 | 0.475181 |
| classM | p-S6 | PMA_I | 15min | 38 | 0.556589 | 0.073851 | 0.283948 | 0.849301 |
| classM | p-S6 | PMA_I | 30min | 38 | 0.696822 | 0.09184 | 0.301146 | 1.072088 |
| classM | p-p65 | pV | unstim | 38 | 0 | 0 | 0 | 0 |
| classM | p-p65 | pV | 5min | 37 | -0.21756 | 0.03181 | -0.35532 | -0.03615 |
| classM | p-p65 | pV | 15min | 37 | -0.25516 | 0.030181 | -0.3479 | -0.10013 |
| classM | p-p65 | pV | 30min | 38 | -0.31564 | 0.030258 | -0.42635 | -0.17847 |
| classM | p-p65 | PMA_I | unstim | 38 | 0 | 0 | 0 | 0 |
| classM | p-p65 | PMA_I | 5min | 37 | -0.11152 | 0.019367 | -0.15973 | -0.04195 |
| classM | p-p65 | PMA_I | 15min | 38 | -0.19985 | 0.025844 | -0.30154 | -0.07527 |
| classM | p-p65 | PMA_I | 30min | 38 | -0.3264 | 0.034056 | -0.46303 | -0.17021 |

**Supplementary Table 2B:**
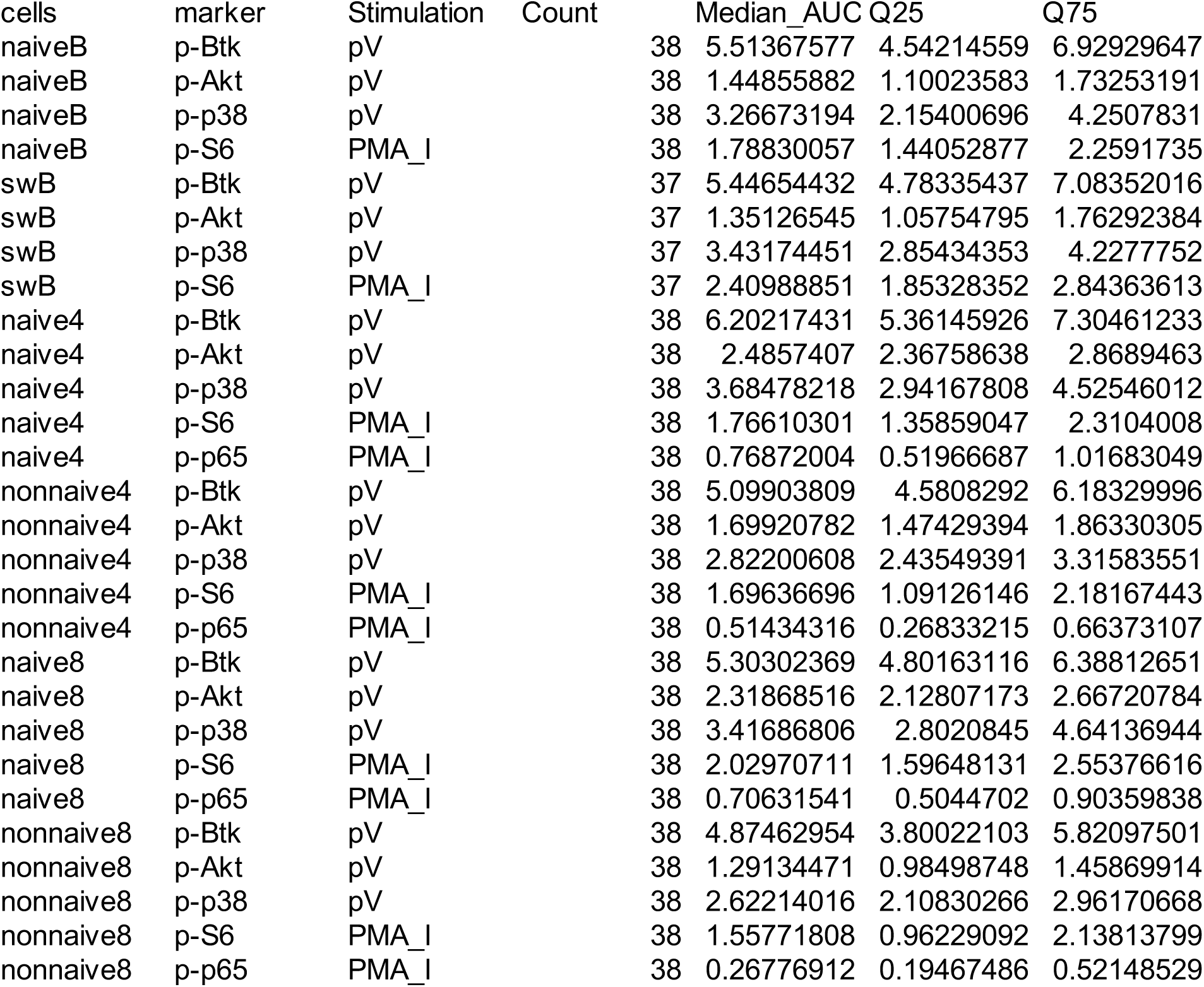
FIGURE 2A_AUC_DATA.

**Supplementary Table 2C:** FIGURE 2A_LINEPLOT_DATA.

| cells | marker | Stimulation | Timing | Count | Mean | SEM | Q25 | Q75 |
| --- | --- | --- | --- | --- | --- | --- | --- | --- |
| naiveB | p-Btk | pV | unstim | 38 | 0 | 0 | 0 | 0 |
| naiveB | p-Btk | pV | 5min | 38 | 2.348876 | 0.132736 | 2.00609 | 2.922822 |
| naiveB | p-Btk | pV | 15min | 38 | 2.398045 | 0.109816 | 1.965414 | 2.880376 |
| naiveB | p-Btk | pV | 30min | 38 | 2.039492 | 0.134098 | 1.401541 | 2.670396 |
| naiveB | p-Akt | pV | unstim | 38 | 0 | 0 | 0 | 0 |
| naiveB | p-Akt | pV | 5min | 38 | 0.750606 | 0.045934 | 0.591966 | 0.913461 |
| naiveB | p-Akt | pV | 15min | 38 | 0.503557 | 0.044662 | 0.306438 | 0.679122 |
| naiveB | p-Akt | pV | 30min | 38 | 0.360986 | 0.033132 | 0.191144 | 0.517808 |
| naiveB | p-p38 | pV | unstim | 38 | 0 | 0 | 0 | 0 |
| naiveB | p-p38 | pV | 5min | 38 | 1.284688 | 0.118546 | 0.744739 | 1.781411 |
| naiveB | p-p38 | pV | 15min | 38 | 1.455459 | 0.09231 | 1.083534 | 1.888605 |
| naiveB | p-p38 | pV | 30min | 38 | 0.945826 | 0.121136 | 0.223056 | 1.458396 |
| naiveB | p-S6 | PMA_I | unstim | 38 | 0 | 0 | 0 | 0 |
| naiveB | p-S6 | PMA_I | 5min | 37 | 0.305875 | 0.030933 | 0.143668 | 0.442142 |
| naiveB | p-S6 | PMA_I | 15min | 38 | 0.889615 | 0.050792 | 0.720368 | 1.127482 |
| naiveB | p-S6 | PMA_I | 30min | 38 | 1.173711 | 0.041101 | 0.972287 | 1.417727 |
| swB | p-Btk | pV | unstim | 38 | 0 | 0 | 0 | 0 |
| swB | p-Btk | pV | 5min | 38 | 2.286829 | 0.13555 | 1.980662 | 2.942988 |
| swB | p-Btk | pV | 15min | 38 | 2.412409 | 0.102806 | 2.090087 | 2.80566 |
| swB | p-Btk | pV | 30min | 37 | 2.096587 | 0.12732 | 1.590012 | 2.657084 |
| swB | p-Akt | pV | unstim | 38 | 0 | 0 | 0 | 0 |
| swB | p-Akt | pV | 5min | 38 | 0.723415 | 0.048891 | 0.525898 | 0.925516 |
| swB | p-Akt | pV | 15min | 38 | 0.512235 | 0.044393 | 0.327123 | 0.679377 |
| swB | p-Akt | pV | 30min | 37 | 0.343926 | 0.024885 | 0.239751 | 0.440213 |
| swB | p-p38 | pV | unstim | 38 | 0 | 0 | 0 | 0 |
| swB | p-p38 | pV | 5min | 38 | 1.376404 | 0.100915 | 1.143668 | 1.807467 |
| swB | p-p38 | pV | 15min | 38 | 1.562804 | 0.079348 | 1.378474 | 1.814296 |
| swB | p-p38 | pV | 30min | 37 | 1.115528 | 0.112722 | 0.446706 | 1.692513 |
| swB | p-S6 | PMA_I | unstim | 38 | 0 | 0 | 0 | 0 |
| swB | p-S6 | PMA_I | 5min | 37 | 0.384205 | 0.042262 | 0.235077 | 0.563048 |
| swB | p-S6 | PMA_I | 15min | 37 | 1.166936 | 0.057001 | 0.954488 | 1.444816 |
| swB | p-S6 | PMA_I | 30min | 37 | 1.524164 | 0.046042 | 1.331837 | 1.79014 |
| naive4 | p-Btk | pV | unstim | 38 | 0 | 0 | 0 | 0 |
| naive4 | p-Btk | pV | 5min | 38 | 2.724154 | 0.102096 | 2.245622 | 3.079467 |
| naive4 | p-Btk | pV | 15min | 38 | 2.560832 | 0.116545 | 2.226529 | 2.970756 |
| naive4 | p-Btk | pV | 30min | 38 | 2.183574 | 0.116046 | 1.641302 | 2.642872 |
| naive4 | p-Akt | pV | unstim | 38 | 0 | 0 | 0 | 0 |
| naive4 | p-Akt | pV | 5min | 38 | 1.195935 | 0.04015 | 1.028503 | 1.366778 |
| naive4 | p-Akt | pV | 15min | 38 | 1.00439 | 0.049559 | 0.786036 | 1.23473 |
| naive4 | p-Akt | pV | 30min | 38 | 0.726961 | 0.051081 | 0.462137 | 0.938241 |
| naive4 | p-p38 | pV | unstim | 38 | 0 | 0 | 0 | 0 |
| naive4 | p-p38 | pV | 5min | 38 | 1.504436 | 0.101283 | 1.125676 | 1.981519 |
| naive4 | p-p38 | pV | 15min | 38 | 1.609193 | 0.070549 | 1.310403 | 1.974804 |
| naive4 | p-p38 | pV | 30min | 38 | 1.277907 | 0.089039 | 0.807601 | 1.697351 |
| naive4 | p-S6 | PMA_I | unstim | 38 | 0 | 0 | 0 | 0 |
| naive4 | p-S6 | PMA_I | 5min | 37 | 0.277801 | 0.028055 | 0.133091 | 0.432097 |
| naive4 | p-S6 | PMA_I | 15min | 38 | 0.824266 | 0.0653 | 0.557664 | 1.124154 |
| naive4 | p-S6 | PMA_I | 30min | 38 | 1.393173 | 0.054705 | 1.240291 | 1.650134 |
| naive4 | p-p65 | PMA_I | unstim | 38 | 0 | 0 | 0 | 0 |
| naive4 | p-p65 | PMA_I | 5min | 37 | 0.12137 | 0.022406 | 0.008532 | 0.220724 |
| naive4 | p-p65 | PMA_I | 15min | 38 | 0.405122 | 0.033176 | 0.307804 | 0.548649 |
| naive4 | p-p65 | PMA_I | 30min | 38 | 0.402622 | 0.019953 | 0.3293 | 0.488003 |
| nonnaive4 | p-Btk | pV | unstim | 38 | 0 | 0 | 0 | 0 |
| nonnaive4 | p-Btk | pV | 5min | 38 | 2.347314 | 0.088044 | 1.904067 | 2.641878 |
| nonnaive4 | p-Btk | pV | 15min | 38 | 2.104544 | 0.103324 | 1.82518 | 2.393865 |
| nonnaive4 | p-Btk | pV | 30min | 38 | 1.703454 | 0.108431 | 1.157801 | 2.066565 |
| nonnaive4 | p-Akt | pV | unstim | 38 | 0 | 0 | 0 | 0 |
| nonnaive4 | p-Akt | pV | 5min | 38 | 0.773773 | 0.035623 | 0.646199 | 0.875252 |
| nonnaive4 | p-Akt | pV | 15min | 38 | 0.628214 | 0.039674 | 0.477608 | 0.802854 |
| nonnaive4 | p-Akt | pV | 30min | 38 | 0.495843 | 0.039262 | 0.343956 | 0.634843 |
| nonnaive4 | p-p38 | pV | unstim | 38 | 0 | 0 | 0 | 0 |
| nonnaive4 | p-p38 | pV | 5min | 38 | 1.227656 | 0.083869 | 0.857168 | 1.578717 |
| nonnaive4 | p-p38 | pV | 15min | 38 | 1.19533 | 0.062385 | 0.87412 | 1.500156 |
| nonnaive4 | p-p38 | pV | 30min | 38 | 0.949004 | 0.07737 | 0.655674 | 1.17781 |
| nonnaive4 | p-S6 | PMA_I | unstim | 38 | 0 | 0 | 0 | 0 |
| nonnaive4 | p-S6 | PMA_I | 5min | 37 | 0.266439 | 0.034016 | 0.087595 | 0.344913 |
| nonnaive4 | p-S6 | PMA_I | 15min | 38 | 0.793302 | 0.067464 | 0.550933 | 1.080731 |
| nonnaive4 | p-S6 | PMA_I | 30min | 38 | 1.221576 | 0.078546 | 0.91193 | 1.652744 |
| nonnaive4 | p-p65 | PMA_I | unstim | 38 | 0 | 0 | 0 | 0 |
| nonnaive4 | p-p65 | PMA_I | 5min | 37 | 0.081989 | 0.016252 | 0.016183 | 0.115531 |
| nonnaive4 | p-p65 | PMA_I | 15min | 38 | 0.280726 | 0.026662 | 0.149604 | 0.396386 |
| nonnaive4 | p-p65 | PMA_I | 30min | 38 | 0.23954 | 0.021596 | 0.132975 | 0.336916 |
| naive8 | p-Btk | pV | unstim | 38 | 0 | 0 | 0 | 0 |
| naive8 | p-Btk | pV | 5min | 38 | 2.526801 | 0.095391 | 2.09089 | 2.814184 |
| naive8 | p-Btk | pV | 15min | 38 | 2.178371 | 0.127082 | 1.822217 | 2.747086 |
| naive8 | p-Btk | pV | 30min | 38 | 1.757696 | 0.111371 | 1.136998 | 2.248558 |
| naive8 | p-Akt | pV | unstim | 38 | 0 | 0 | 0 | 0 |
| naive8 | p-Akt | pV | 5min | 38 | 1.183404 | 0.043988 | 1.019349 | 1.303398 |
| naive8 | p-Akt | pV | 15min | 38 | 0.904277 | 0.054634 | 0.694783 | 1.134656 |
| naive8 | p-Akt | pV | 30min | 38 | 0.664554 | 0.051295 | 0.395346 | 0.907684 |
| naive8 | p-p38 | pV | unstim | 38 | 0 | 0 | 0 | 0 |
| naive8 | p-p38 | pV | 5min | 38 | 1.517338 | 0.088401 | 1.202368 | 1.860335 |
| naive8 | p-p38 | pV | 15min | 38 | 1.480071 | 0.071788 | 1.108477 | 1.758176 |
| naive8 | p-p38 | pV | 30min | 38 | 1.12097 | 0.088863 | 0.766305 | 1.580475 |
| naive8 | p-S6 | PMA_I | unstim | 38 | 0 | 0 | 0 | 0 |
| naive8 | p-S6 | PMA_I | 5min | 37 | 0.338263 | 0.032631 | 0.156924 | 0.473244 |
| naive8 | p-S6 | PMA_I | 15min | 38 | 0.93458 | 0.0655 | 0.632392 | 1.209201 |
| naive8 | p-S6 | PMA_I | 30min | 38 | 1.479687 | 0.05237 | 1.319815 | 1.702638 |
| naive8 | p-p65 | PMA_I | unstim | 38 | 0 | 0 | 0 | 0 |
| naive8 | p-p65 | PMA_I | 5min | 37 | 0.081866 | 0.019729 | -0.00197 | 0.163428 |
| naive8 | p-p65 | PMA_I | 15min | 38 | 0.399304 | 0.032279 | 0.270074 | 0.545167 |
| naive8 | p-p65 | PMA_I | 30min | 38 | 0.386233 | 0.020231 | 0.328631 | 0.480588 |
| nonnaive8 | p-Btk | pV | unstim | 38 | 0 | 0 | 0 | 0 |
| nonnaive8 | p-Btk | pV | 5min | 38 | 2.1798 | 0.116648 | 1.810188 | 2.544114 |
| nonnaive8 | p-Btk | pV | 15min | 38 | 1.888358 | 0.121588 | 1.659643 | 2.243617 |
| nonnaive8 | p-Btk | pV | 30min | 38 | 1.388899 | 0.124803 | 0.985825 | 1.822869 |
| nonnaive8 | p-Akt | pV | unstim | 38 | 0 | 0 | 0 | 0 |
| nonnaive8 | p-Akt | pV | 5min | 38 | 0.604283 | 0.029398 | 0.466965 | 0.743656 |
| nonnaive8 | p-Akt | pV | 15min | 38 | 0.444 | 0.032659 | 0.341869 | 0.591934 |
| nonnaive8 | p-Akt | pV | 30min | 38 | 0.339445 | 0.035059 | 0.176541 | 0.461162 |
| nonnaive8 | p-p38 | pV | unstim | 38 | 0 | 0 | 0 | 0 |
| nonnaive8 | p-p38 | pV | 5min | 38 | 1.143901 | 0.085846 | 0.854404 | 1.423418 |
| nonnaive8 | p-p38 | pV | 15min | 38 | 1.06983 | 0.075812 | 0.685492 | 1.433823 |
| nonnaive8 | p-p38 | pV | 30min | 38 | 0.807769 | 0.089929 | 0.46241 | 1.136138 |
| nonnaive8 | p-S6 | PMA_I | unstim | 38 | 0 | 0 | 0 | 0 |
| nonnaive8 | p-S6 | PMA_I | 5min | 37 | 0.236441 | 0.034509 | 0.070425 | 0.349535 |
| nonnaive8 | p-S6 | PMA_I | 15min | 38 | 0.729716 | 0.084359 | 0.330876 | 1.116381 |
| nonnaive8 | p-S6 | PMA_I | 30min | 38 | 1.216644 | 0.096441 | 0.852636 | 1.563109 |
| nonnaive8 | p-p65 | PMA_I | unstim | 38 | 0 | 0 | 0 | 0 |
| nonnaive8 | p-p65 | PMA_I | 5min | 37 | 0.068242 | 0.017218 | -0.00234 | 0.107103 |
| nonnaive8 | p-p65 | PMA_I | 15min | 38 | 0.202432 | 0.028355 | 0.08513 | 0.329791 |
| nonnaive8 | p-p65 | PMA_I | 30min | 38 | 0.139444 | 0.018068 | 0.053789 | 0.201883 |

**Supplementary Table 2D:** SF2_MFI_DATA.

| cells | marker | Stimulation | Timing | Count | Median | Q25 | Q75 | p.adj | p.adj.signif |
| --- | --- | --- | --- | --- | --- | --- | --- | --- | --- |
| CD19BC | p-Btk | pV | unstim | 38 | 0.271228 | 0.120908 | 0.50427 | NA | NA |
| CD19BC | p-Btk | pV | 5min | 38 | 2.738207 | 2.312843 | 3.244524 | 8.60E-18 | **** |
| CD19BC | p-Btk | pV | 15min | 38 | 2.866527 | 2.478843 | 3.359421 | 8.00E-22 | **** |
| CD19BC | p-Btk | pV | 30min | 38 | 2.385788 | 2.022321 | 3.071144 | 5.40E-17 | **** |
| CD19BC | p-Btk | PMA_I | unstim | 38 | 0.488728 | 0.31658 | 0.747 | NA | NA |
| CD19BC | p-Btk | PMA_I | 5min | 37 | 0.427996 | 0.253837 | 0.640875 | 0.28 | ns |
| CD19BC | p-Btk | PMA_I | 15min | 38 | 0.417037 | 0.246313 | 0.684194 | 0.34 | ns |
| CD19BC | p-Btk | PMA_I | 30min | 38 | 0.416383 | 0.228371 | 0.657265 | 0.19 | ns |
| CD19BC | p-Akt | pV | unstim | 38 | 0.15817 | 0.097397 | 0.244795 | NA | NA |
| CD19BC | p-Akt | pV | 5min | 38 | 0.862789 | 0.663555 | 1.091038 | 2.70E-15 | **** |
| CD19BC | p-Akt | pV | 15min | 38 | 0.665598 | 0.459588 | 0.806097 | 2.40E-11 | **** |
| CD19BC | p-Akt | pV | 30min | 38 | 0.524691 | 0.332629 | 0.631502 | 8.20E-09 | **** |
| CD19BC | p-Akt | PMA_I | unstim | 38 | 0.140581 | 0.036291 | 0.195013 | NA | NA |
| CD19BC | p-Akt | PMA_I | 5min | 37 | 0.354346 | 0.210609 | 0.436294 | 5.80E-08 | **** |
| CD19BC | p-Akt | PMA_I | 15min | 38 | 0.376545 | 0.27357 | 0.509594 | 1.70E-09 | **** |
| CD19BC | p-Akt | PMA_I | 30min | 38 | 0.389866 | 0.285896 | 0.50159 | 1.60E-10 | **** |
| CD19BC | p-p38 | pV | unstim | 38 | 0.604348 | 0.396778 | 0.786447 | NA | NA |
| CD19BC | p-p38 | pV | 5min | 38 | 1.923421 | 1.519191 | 2.369437 | 1.50E-13 | **** |
| CD19BC | p-p38 | pV | 15min | 38 | 2.115113 | 1.851958 | 2.406978 | 1.50E-18 | **** |
| CD19BC | p-p38 | pV | 30min | 38 | 1.666109 | 0.999484 | 2.155507 | 1.70E-09 | **** |
| CD19BC | p-p38 | PMA_I | unstim | 38 | 0.607182 | 0.444831 | 0.815031 | NA | NA |
| CD19BC | p-p38 | PMA_I | 5min | 37 | 0.627945 | 0.497002 | 0.850302 | 0.68 | ns |
| CD19BC | p-p38 | PMA_I | 15min | 38 | 0.710165 | 0.510138 | 0.892591 | 0.34 | ns |
| CD19BC | p-p38 | PMA_I | 30min | 38 | 0.723002 | 0.514107 | 0.861664 | 0.33 | ns |
| CD19BC | p-S6 | pV | unstim | 38 | 0.308841 | 0.258513 | 0.378662 | NA | NA |
| CD19BC | p-S6 | pV | 5min | 38 | 0.414225 | 0.332742 | 0.482981 | 0.055 | * |
| CD19BC | p-S6 | pV | 15min | 38 | 0.314205 | 0.269359 | 0.405287 | 0.62 | ns |
| CD19BC | p-S6 | pV | 30min | 38 | 0.23421 | 0.182531 | 0.277612 | 0.00018 | **** |
| CD19BC | p-S6 | PMA_I | unstim | 38 | 0.323934 | 0.269267 | 0.359887 | NA | NA |
| CD19BC | p-S6 | PMA_I | 5min | 37 | 0.568017 | 0.444475 | 0.847718 | 4.50E-07 | **** |
| CD19BC | p-S6 | PMA_I | 15min | 38 | 1.369581 | 1.067228 | 1.518609 | 1.50E-18 | **** |
| CD19BC | p-S6 | PMA_I | 30min | 38 | 1.631633 | 1.379029 | 1.755446 | 1.70E-28 | **** |
| CD3TC | p-Btk | pV | unstim | 38 | 0.010655 | -0.16942 | 0.189084 | NA | NA |
| CD3TC | p-Btk | pV | 5min | 38 | 2.64364 | 2.204592 | 3.204963 | 1.60E-26 | **** |
| CD3TC | p-Btk | pV | 15min | 38 | 2.391149 | 2.092833 | 2.76162 | 1.60E-26 | **** |
| CD3TC | p-Btk | pV | 30min | 38 | 1.945777 | 1.407341 | 2.445378 | 3.20E-17 | **** |
| CD3TC | p-Btk | PMA_I | unstim | 38 | 0.139759 | 0.002656 | 0.418002 | NA | NA |
| CD3TC | p-Btk | PMA_I | 5min | 37 | 0.158848 | -0.04658 | 0.332231 | 0.68 | ns |
| CD3TC | p-Btk | PMA_I | 15min | 38 | 0.107824 | -0.06497 | 0.325471 | 0.33 | ns |
| CD3TC | p-Btk | PMA_I | 30min | 38 | 0.048516 | -0.11116 | 0.215896 | 0.1 | ns |
| CD3TC | p-Akt | pV | unstim | 38 | 0.349057 | 0.287653 | 0.42256 | NA | NA |
| CD3TC | p-Akt | pV | 5min | 38 | 1.243354 | 1.11191 | 1.405173 | 1.00E-25 | **** |
| CD3TC | p-Akt | pV | 15min | 38 | 1.090947 | 0.829932 | 1.230779 | 7.50E-19 | **** |
| CD3TC | p-Akt | pV | 30min | 38 | 0.808688 | 0.598793 | 1.155619 | 1.50E-10 | **** |
| CD3TC | p-Akt | PMA_I | unstim | 38 | 0.314752 | 0.230942 | 0.381472 | NA | NA |
| CD3TC | p-Akt | PMA_I | 5min | 37 | 0.459578 | 0.383694 | 0.545981 | 8.00E-06 | **** |
| CD3TC | p-Akt | PMA_I | 15min | 38 | 0.494991 | 0.435702 | 0.55942 | 6.70E-08 | **** |
| CD3TC | p-Akt | PMA_I | 30min | 38 | 0.556033 | 0.48251 | 0.62626 | 1.20E-09 | **** |
| CD3TC | p-p38 | pV | unstim | 38 | 0.239279 | 0.140478 | 0.376216 | NA | NA |
| CD3TC | p-p38 | pV | 5min | 38 | 1.628554 | 1.189016 | 2.025218 | 2.90E-17 | **** |
| CD3TC | p-p38 | pV | 15min | 38 | 1.741819 | 1.355292 | 1.998596 | 6.00E-21 | **** |
| CD3TC | p-p38 | pV | 30min | 38 | 1.330399 | 0.866053 | 1.68215 | 1.90E-13 | **** |
| CD3TC | p-p38 | PMA_I | unstim | 38 | 0.271519 | 0.166632 | 0.425022 | NA | NA |
| CD3TC | p-p38 | PMA_I | 5min | 37 | 0.340626 | 0.180362 | 0.464075 | 0.32 | ns |
| CD3TC | p-p38 | PMA_I | 15min | 38 | 0.37513 | 0.235825 | 0.493048 | 0.084 | ns |
| CD3TC | p-p38 | PMA_I | 30min | 38 | 0.379504 | 0.237646 | 0.502469 | 0.088 | ns |
| CD3TC | p-S6 | pV | unstim | 38 | 0.166392 | 0.143864 | 0.214065 | NA | NA |
| CD3TC | p-S6 | pV | 5min | 38 | 0.312127 | 0.252963 | 0.387449 | 1.90E-08 | **** |
| CD3TC | p-S6 | pV | 15min | 38 | 0.277663 | 0.217494 | 0.374667 | 9.10E-06 | **** |
| CD3TC | p-S6 | pV | 30min | 38 | 0.21762 | 0.171423 | 0.258314 | 0.046 | * |
| CD3TC | p-S6 | PMA_I | unstim | 38 | 0.151139 | 0.136805 | 0.203669 | NA | NA |
| CD3TC | p-S6 | PMA_I | 5min | 37 | 0.365049 | 0.283503 | 0.578039 | 7.50E-09 | **** |
| CD3TC | p-S6 | PMA_I | 15min | 38 | 0.99445 | 0.715163 | 1.286328 | 3.00E-13 | **** |
| CD3TC | p-S6 | PMA_I | 30min | 38 | 1.535402 | 1.33973 | 1.770011 | 1.10E-21 | **** |
| CD3TC | p-p65 | pV | unstim | 38 | 0.417931 | 0.292288 | 0.574677 | NA | NA |
| CD3TC | p-p65 | pV | 5min | 38 | 0.735288 | 0.601471 | 0.897743 | 1.30E-08 | **** |
| CD3TC | p-p65 | pV | 15min | 38 | 0.621571 | 0.530389 | 0.767042 | 1.10E-05 | **** |
| CD3TC | p-p65 | pV | 30min | 38 | 0.516876 | 0.289865 | 0.59923 | 0.15 | ns |
| CD3TC | p-p65 | PMA_I | unstim | 38 | 0.394746 | 0.268094 | 0.576982 | NA | NA |
| CD3TC | p-p65 | PMA_I | 5min | 37 | 0.47529 | 0.3173 | 0.681576 | 0.07 | * |
| CD3TC | p-p65 | PMA_I | 15min | 38 | 0.693166 | 0.546018 | 0.907558 | 2.10E-07 | **** |
| CD3TC | p-p65 | PMA_I | 30min | 38 | 0.673973 | 0.588727 | 0.847847 | 7.20E-09 | **** |
| NK | p-Btk | pV | unstim | 38 | 0.287505 | 0.184351 | 0.518523 | NA | NA |
| NK | p-Btk | pV | 5min | 38 | 1.873478 | 0.428225 | 2.453973 | 5.90E-08 | **** |
| NK | p-Btk | pV | 15min | 38 | 1.156092 | 0.459653 | 1.801373 | 6.30E-08 | **** |
| NK | p-Btk | pV | 30min | 38 | 0.557205 | 0.345935 | 1.592317 | 1.20E-05 | **** |
| NK | p-Btk | PMA_I | unstim | 38 | 0.501397 | 0.402125 | 0.828429 | NA | NA |
| NK | p-Btk | PMA_I | 5min | 37 | 0.48771 | 0.365286 | 0.650173 | 0.3 | ns |
| NK | p-Btk | PMA_I | 15min | 38 | 0.387778 | 0.280541 | 0.530513 | 0.026 | * |
| NK | p-Btk | PMA_I | 30min | 38 | 0.284015 | 0.221223 | 0.411913 | 0.00013 | **** |
| NK | p-Akt | pV | unstim | 38 | 0.456055 | 0.31258 | 0.572097 | NA | NA |
| NK | p-Akt | pV | 5min | 38 | 0.844336 | 0.347809 | 1.018393 | 0.0019 | ** |
| NK | p-Akt | pV | 15min | 38 | 0.615893 | 0.372991 | 0.829629 | 0.072 | * |
| NK | p-Akt | pV | 30min | 38 | 0.513457 | 0.340806 | 0.695084 | 0.36 | ns |
| NK | p-Akt | PMA_I | unstim | 38 | 0.406031 | 0.269042 | 0.580835 | NA | NA |
| NK | p-Akt | PMA_I | 5min | 37 | 0.481449 | 0.282932 | 0.677522 | 0.37 | ns |
| NK | p-Akt | PMA_I | 15min | 38 | 0.411272 | 0.13 | 0.618994 | 0.94 | ns |
| NK | p-Akt | PMA_I | 30min | 38 | 0.323662 | 0.116977 | 0.509088 | 0.35 | ns |
| NK | p-p38 | pV | unstim | 38 | 0.200322 | -0.0204 | 0.348908 | NA | NA |
| NK | p-p38 | pV | 5min | 38 | 0.810516 | 0.322955 | 1.396146 | 1.10E-07 | **** |
| NK | p-p38 | pV | 15min | 38 | 0.710625 | 0.354366 | 1.024571 | 5.90E-08 | **** |
| NK | p-p38 | pV | 30min | 38 | 0.413343 | 0.225801 | 0.873832 | 4.00E-05 | **** |
| NK | p-p38 | PMA_I | unstim | 38 | 0.193001 | 0.073992 | 0.382277 | NA | NA |
| NK | p-p38 | PMA_I | 5min | 37 | 0.237041 | 0.115799 | 0.394658 | 0.42 | ns |
| NK | p-p38 | PMA_I | 15min | 38 | 0.243086 | 0.11989 | 0.333417 | 0.42 | ns |
| NK | p-p38 | PMA_I | 30min | 38 | 0.193029 | 0.095657 | 0.342492 | 0.76 | ns |
| NK | p-S6 | pV | unstim | 38 | 0.203509 | 0.123669 | 0.283026 | NA | NA |
| NK | p-S6 | pV | 5min | 38 | 0.211671 | 0.089269 | 0.296654 | 0.94 | ns |
| NK | p-S6 | pV | 15min | 38 | 0.189367 | 0.08742 | 0.263175 | 0.35 | ns |
| NK | p-S6 | pV | 30min | 38 | 0.146077 | 0.07631 | 0.19722 | 0.023 | * |
| NK | p-S6 | PMA_I | unstim | 38 | 0.197701 | 0.111646 | 0.235188 | NA | NA |
| NK | p-S6 | PMA_I | 5min | 37 | 0.265069 | 0.149424 | 0.364765 | 0.04 | * |
| NK | p-S6 | PMA_I | 15min | 38 | 0.226249 | 0.089041 | 0.417102 | 0.034 | * |
| NK | p-S6 | PMA_I | 30min | 38 | 0.19246 | 0.089613 | 0.852439 | 0.0019 | ** |
| NK | p-p65 | pV | unstim | 38 | 0.29737 | 0.209891 | 0.361365 | NA | NA |
| NK | p-p65 | pV | 5min | 38 | 0.461239 | 0.318006 | 0.665482 | 2.00E-04 | **** |
| NK | p-p65 | pV | 15min | 38 | 0.319787 | 0.222246 | 0.43367 | 0.27 | ns |
| NK | p-p65 | pV | 30min | 38 | 0.257029 | 0.162435 | 0.319555 | 0.37 | ns |
| NK | p-p65 | PMA_I | unstim | 38 | 0.293693 | 0.224642 | 0.360034 | NA | NA |
| NK | p-p65 | PMA_I | 5min | 37 | 0.322977 | 0.2357 | 0.472428 | 0.12 | ns |
| NK | p-p65 | PMA_I | 15min | 38 | 0.35417 | 0.269922 | 0.420893 | 0.084 | ns |
| NK | p-p65 | PMA_I | 30min | 38 | 0.315246 | 0.275353 | 0.365967 | 0.55 | ns |
| classM | p-Btk | pV | unstim | 38 | 0.712002 | 0.559831 | 1.028372 | NA | NA |
| classM | p-Btk | pV | 5min | 37 | 1.942649 | 1.380062 | 2.381747 | 1.90E-12 | **** |
| classM | p-Btk | pV | 15min | 37 | 2.030524 | 1.538566 | 2.314629 | 1.30E-14 | **** |
| classM | p-Btk | pV | 30min | 38 | 2.463315 | 1.946019 | 2.790941 | 2.20E-20 | **** |
| classM | p-Btk | PMA_I | unstim | 38 | 1.140903 | 0.909076 | 1.503597 | NA | NA |
| classM | p-Btk | PMA_I | 5min | 37 | 0.915498 | 0.756672 | 1.28761 | 0.028 | * |
| classM | p-Btk | PMA_I | 15min | 38 | 0.798471 | 0.681164 | 1.140249 | 0.00014 | **** |
| classM | p-Btk | PMA_I | 30min | 38 | 0.84257 | 0.676631 | 1.108941 | 0.00038 | *** |
| classM | p-Akt | pV | unstim | 38 | 1.667377 | 1.524817 | 1.901181 | NA | NA |
| classM | p-Akt | pV | 5min | 37 | 1.719594 | 1.564236 | 1.932451 | 0.43 | ns |
| classM | p-Akt | pV | 15min | 37 | 1.628294 | 1.481404 | 1.881588 | 0.68 | ns |
| classM | p-Akt | pV | 30min | 38 | 1.542488 | 1.369112 | 1.793074 | 0.1 | ns |
| classM | p-Akt | PMA_I | unstim | 38 | 1.62565 | 1.484899 | 1.832627 | NA | NA |
| classM | p-Akt | PMA_I | 5min | 37 | 1.536507 | 1.406104 | 1.781614 | 0.31 | ns |
| classM | p-Akt | PMA_I | 15min | 38 | 1.559961 | 1.358015 | 1.78191 | 0.2 | ns |
| classM | p-Akt | PMA_I | 30min | 38 | 1.539897 | 1.342381 | 1.741976 | 0.079 | ns |
| classM | p-p38 | pV | unstim | 38 | 1.36243 | 1.182889 | 1.549659 | NA | NA |
| classM | p-p38 | pV | 5min | 37 | 1.955428 | 1.650606 | 2.11133 | 2.20E-06 | **** |
| classM | p-p38 | pV | 15min | 37 | 1.899582 | 1.536964 | 2.185223 | 5.50E-06 | **** |
| classM | p-p38 | pV | 30min | 38 | 2.079594 | 1.780251 | 2.313334 | 6.10E-10 | **** |
| classM | p-p38 | PMA_I | unstim | 38 | 1.495912 | 1.307252 | 1.699098 | NA | NA |
| classM | p-p38 | PMA_I | 5min | 37 | 1.561178 | 1.405704 | 1.762747 | 0.46 | ns |
| classM | p-p38 | PMA_I | 15min | 38 | 1.585359 | 1.318331 | 1.688411 | 0.68 | ns |
| classM | p-p38 | PMA_I | 30min | 38 | 1.596226 | 1.349328 | 1.705674 | 0.64 | ns |
| classM | p-S6 | pV | unstim | 38 | 1.65122 | 1.322944 | 1.888667 | NA | NA |
| classM | p-S6 | pV | 5min | 37 | 1.48331 | 1.203371 | 1.802727 | 0.38 | ns |
| classM | p-S6 | pV | 15min | 37 | 1.389835 | 1.025511 | 1.736739 | 0.079 | ns |
| classM | p-S6 | pV | 30min | 38 | 1.344219 | 0.93621 | 1.509122 | 0.0092 | ** |
| classM | p-S6 | PMA_I | unstim | 38 | 1.616526 | 1.325291 | 1.848168 | NA | NA |
| classM | p-S6 | PMA_I | 5min | 37 | 1.938252 | 1.65539 | 2.102403 | 0.00052 | *** |
| classM | p-S6 | PMA_I | 15min | 38 | 2.147228 | 1.929022 | 2.392744 | 3.10E-09 | **** |
| classM | p-S6 | PMA_I | 30min | 38 | 2.283933 | 2.038354 | 2.465518 | 9.40E-12 | **** |
| classM | p-p65 | pV | unstim | 38 | 1.029756 | 0.78544 | 1.167253 | NA | NA |
| classM | p-p65 | pV | 5min | 37 | 0.790937 | 0.654989 | 0.924814 | 0.035 | * |
| classM | p-p65 | pV | 15min | 37 | 0.78798 | 0.583179 | 0.924674 | 0.0075 | ** |
| classM | p-p65 | pV | 30min | 38 | 0.73378 | 0.535848 | 0.850767 | 0.00041 | *** |
| classM | p-p65 | PMA_I | unstim | 38 | 1.064661 | 0.887795 | 1.217759 | NA | NA |
| classM | p-p65 | PMA_I | 5min | 37 | 0.947782 | 0.773045 | 1.042248 | 0.22 | ns |
| classM | p-p65 | PMA_I | 15min | 38 | 0.831034 | 0.695119 | 0.94496 | 0.011 | ** |
| classM | p-p65 | PMA_I | 30min | 38 | 0.712524 | 0.61072 | 0.825536 | 6.80E-06 | **** |

**Supplementary Table 2E:**
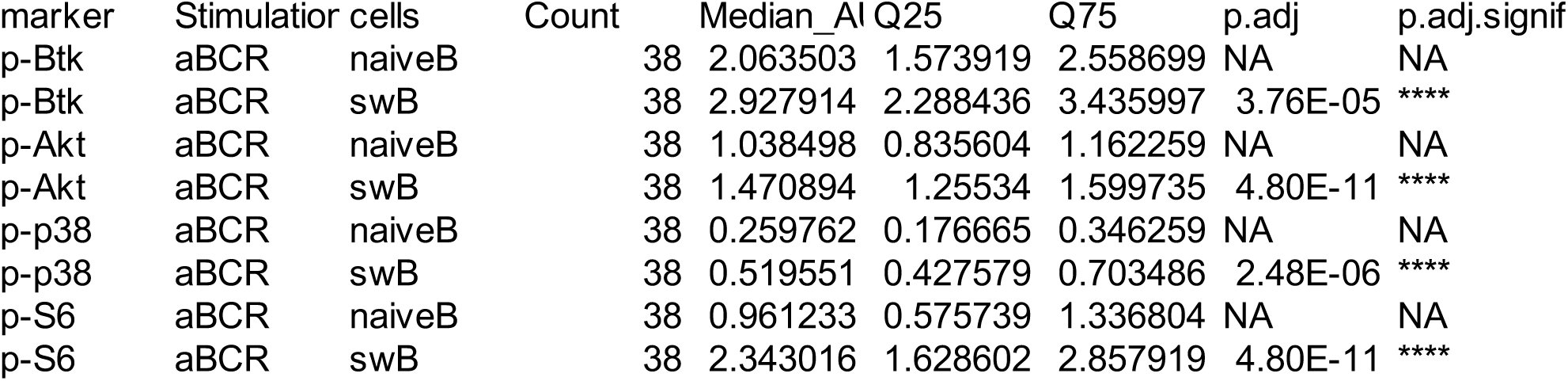
FIGURE 3A_AUC_DATA.

**Supplementary Table 2F:**
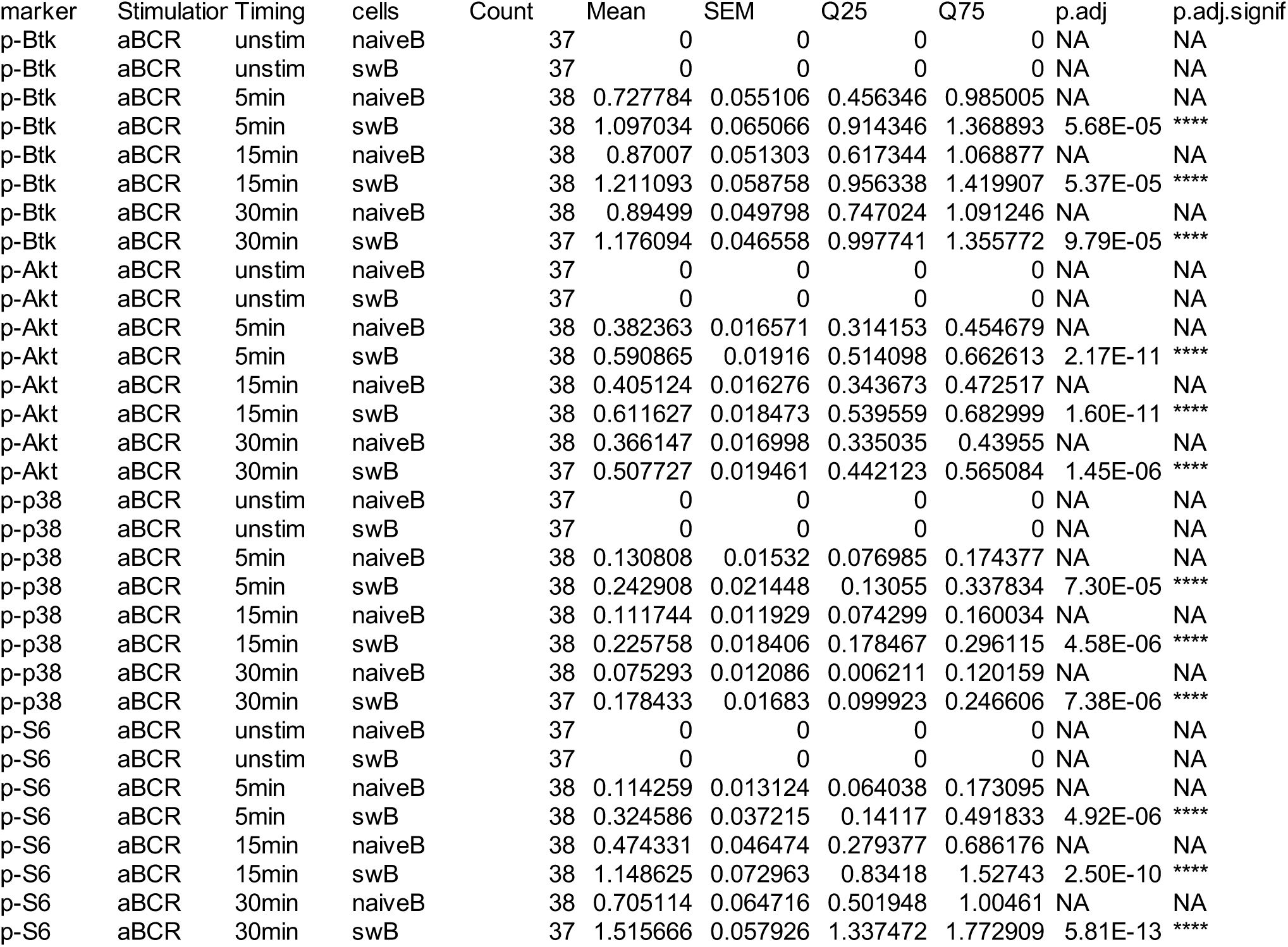
FIGURE 3A_LINEPLOT_DATA.

**Supplementary Table 2G:**
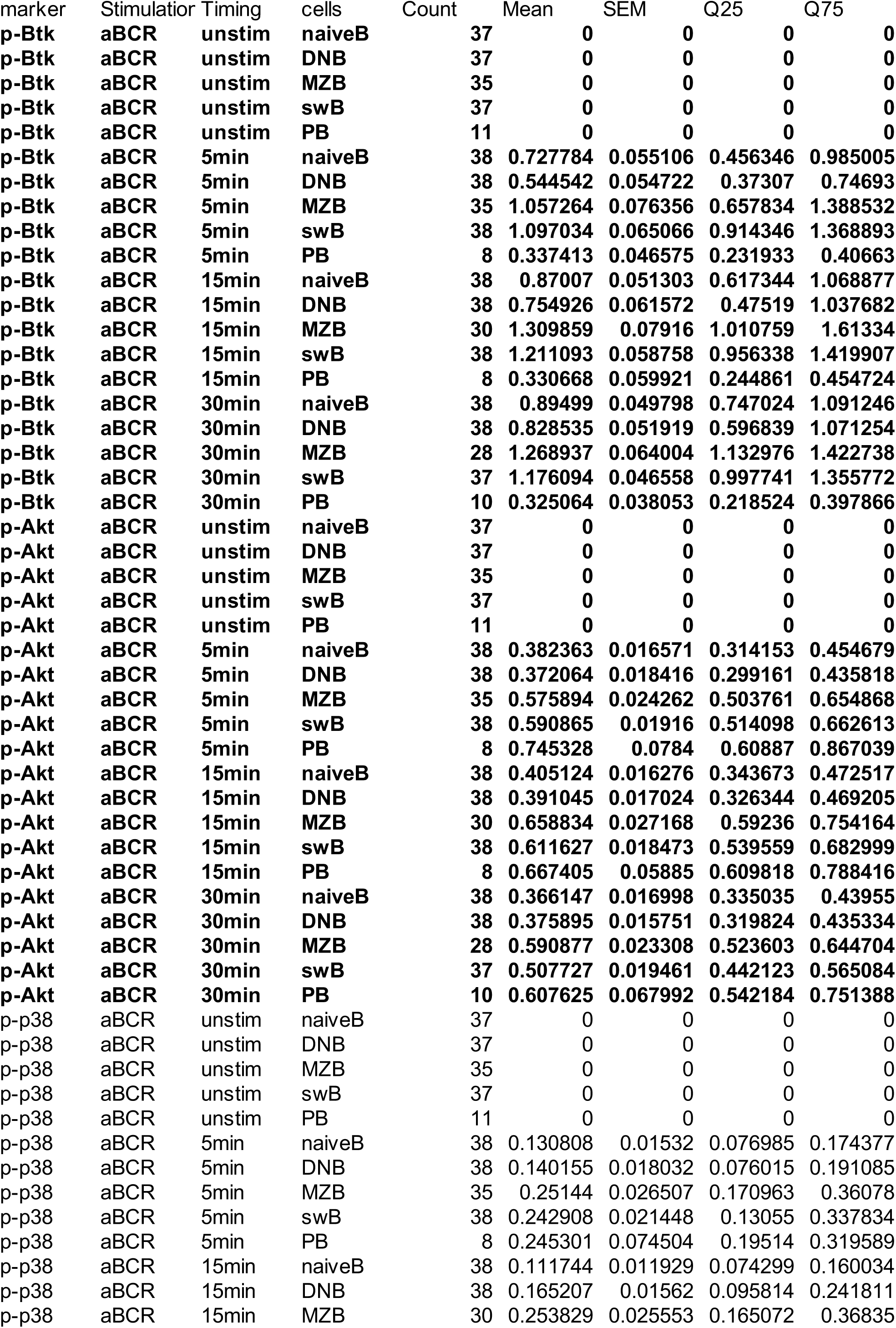
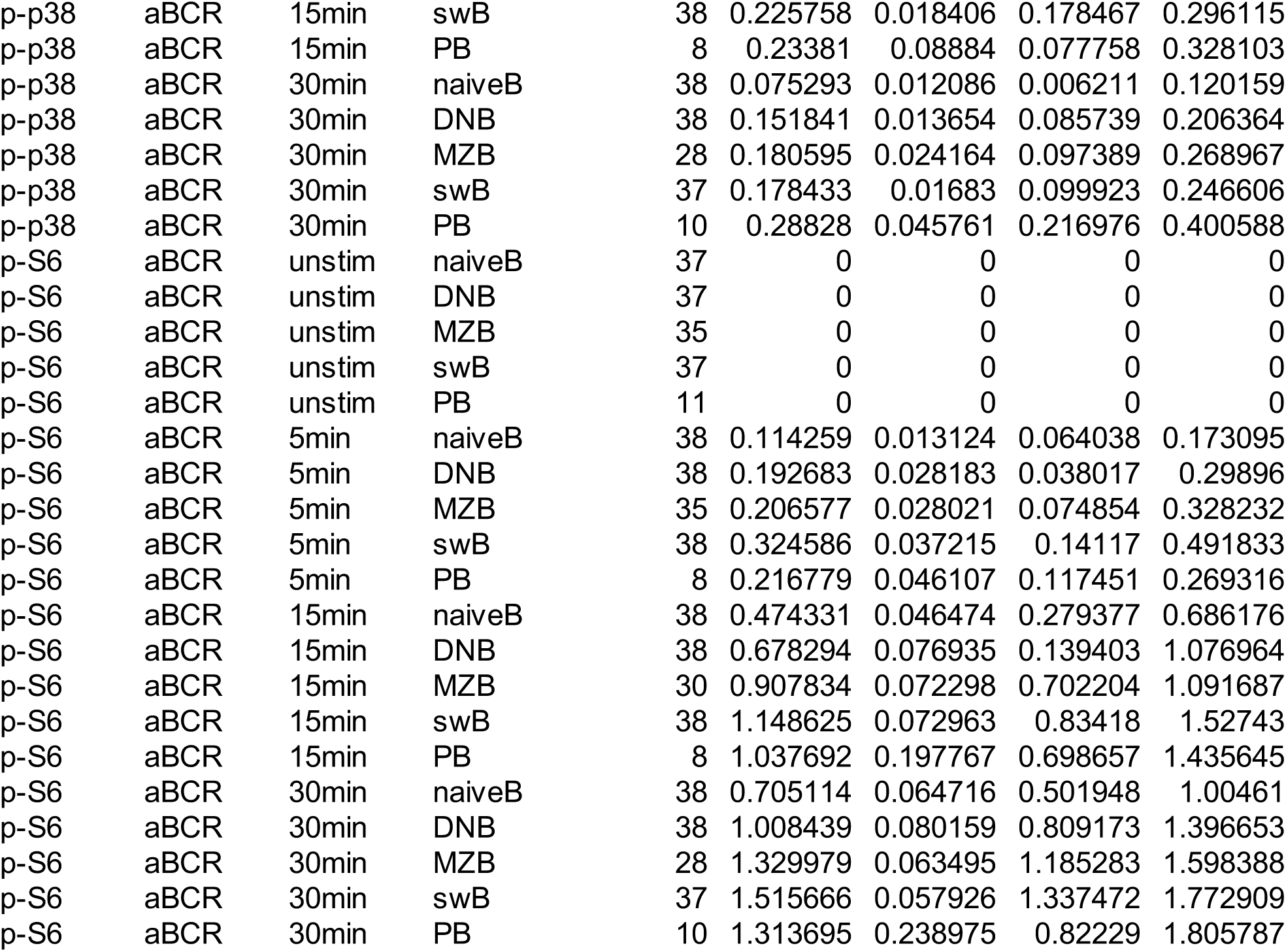
FIGURE 3BC_DATA.

| marker | Stimulation | Timing | cells | Count | Mean | SEM | Q25 | Q75 |
| --- | --- | --- | --- | --- | --- | --- | --- | --- |
| p-Btk | aBCR | unstim | naiveB | 37 | 0 | 0 | 0 | 0 |
| p-Btk | aBCR | unstim | DNB | 37 | 0 | 0 | 0 | 0 |
| p-Btk | aBCR | unstim | MZB | 35 | 0 | 0 | 0 | 0 |
| p-Btk | aBCR | unstim | swB | 37 | 0 | 0 | 0 | 0 |
| p-Btk | aBCR | unstim | PB | 11 | 0 | 0 | 0 | 0 |
| p-Btk | aBCR | 5min | naiveB | 38 | 0.727784 | 0.055106 | 0.456346 | 0.985005 |
| p-Btk | aBCR | 5min | DNB | 38 | 0.544542 | 0.054722 | 0.37307 | 0.74693 |
| p-Btk | aBCR | 5min | MZB | 35 | 1.057264 | 0.076356 | 0.657834 | 1.388532 |
| p-Btk | aBCR | 5min | swB | 38 | 1.097034 | 0.065066 | 0.914346 | 1.368893 |
| p-Btk | aBCR | 5min | PB | 8 | 0.337413 | 0.046575 | 0.231933 | 0.40663 |
| p-Btk | aBCR | 15min | naiveB | 38 | 0.87007 | 0.051303 | 0.617344 | 1.068877 |
| p-Btk | aBCR | 15min | DNB | 38 | 0.754926 | 0.061572 | 0.47519 | 1.037682 |
| p-Btk | aBCR | 15min | MZB | 30 | 1.309859 | 0.07916 | 1.010759 | 1.61334 |
| p-Btk | aBCR | 15min | swB | 38 | 1.211093 | 0.058758 | 0.956338 | 1.419907 |
| p-Btk | aBCR | 15min | PB | 8 | 0.330668 | 0.059921 | 0.244861 | 0.454724 |
| p-Btk | aBCR | 30min | naiveB | 38 | 0.89499 | 0.049798 | 0.747024 | 1.091246 |
| p-Btk | aBCR | 30min | DNB | 38 | 0.828535 | 0.051919 | 0.596839 | 1.071254 |
| p-Btk | aBCR | 30min | MZB | 28 | 1.268937 | 0.064004 | 1.132976 | 1.422738 |
| p-Btk | aBCR | 30min | swB | 37 | 1.176094 | 0.046558 | 0.997741 | 1.355772 |
| p-Btk | aBCR | 30min | PB | 10 | 0.325064 | 0.038053 | 0.218524 | 0.397866 |
| p-Akt | aBCR | unstim | naiveB | 37 | 0 | 0 | 0 | 0 |
| p-Akt | aBCR | unstim | DNB | 37 | 0 | 0 | 0 | 0 |
| p-Akt | aBCR | unstim | MZB | 35 | 0 | 0 | 0 | 0 |
| p-Akt | aBCR | unstim | swB | 37 | 0 | 0 | 0 | 0 |
| p-Akt | aBCR | unstim | PB | 11 | 0 | 0 | 0 | 0 |
| p-Akt | aBCR | 5min | naiveB | 38 | 0.382363 | 0.016571 | 0.314153 | 0.454679 |
| p-Akt | aBCR | 5min | DNB | 38 | 0.372064 | 0.018416 | 0.299161 | 0.435818 |
| p-Akt | aBCR | 5min | MZB | 35 | 0.575894 | 0.024262 | 0.503761 | 0.654868 |
| p-Akt | aBCR | 5min | swB | 38 | 0.590865 | 0.01916 | 0.514098 | 0.662613 |
| p-Akt | aBCR | 5min | PB | 8 | 0.745328 | 0.0784 | 0.60887 | 0.867039 |
| p-Akt | aBCR | 15min | naiveB | 38 | 0.405124 | 0.016276 | 0.343673 | 0.472517 |
| p-Akt | aBCR | 15min | DNB | 38 | 0.391045 | 0.017024 | 0.326344 | 0.469205 |
| p-Akt | aBCR | 15min | MZB | 30 | 0.658834 | 0.027168 | 0.59236 | 0.754164 |
| p-Akt | aBCR | 15min | swB | 38 | 0.611627 | 0.018473 | 0.539559 | 0.682999 |
| p-Akt | aBCR | 15min | PB | 8 | 0.667405 | 0.05885 | 0.609818 | 0.788416 |
| p-Akt | aBCR | 30min | naiveB | 38 | 0.366147 | 0.016998 | 0.335035 | 0.43955 |
| p-Akt | aBCR | 30min | DNB | 38 | 0.375895 | 0.015751 | 0.319824 | 0.435334 |
| p-Akt | aBCR | 30min | MZB | 28 | 0.590877 | 0.023308 | 0.523603 | 0.644704 |
| p-Akt | aBCR | 30min | swB | 37 | 0.507727 | 0.019461 | 0.442123 | 0.565084 |
| p-Akt | aBCR | 30min | PB | 10 | 0.607625 | 0.067992 | 0.542184 | 0.751388 |
| p-p38 | aBCR | unstim | naiveB | 37 | 0 | 0 | 0 | 0 |
| p-p38 | aBCR | unstim | DNB | 37 | 0 | 0 | 0 | 0 |
| p-p38 | aBCR | unstim | MZB | 35 | 0 | 0 | 0 | 0 |
| p-p38 | aBCR | unstim | swB | 37 | 0 | 0 | 0 | 0 |
| p-p38 | aBCR | unstim | PB | 11 | 0 | 0 | 0 | 0 |
| p-p38 | aBCR | 5min | naiveB | 38 | 0.130808 | 0.01532 | 0.076985 | 0.174377 |
| p-p38 | aBCR | 5min | DNB | 38 | 0.140155 | 0.018032 | 0.076015 | 0.191085 |
| p-p38 | aBCR | 5min | MZB | 35 | 0.25144 | 0.026507 | 0.170963 | 0.36078 |
| p-p38 | aBCR | 5min | swB | 38 | 0.242908 | 0.021448 | 0.13055 | 0.337834 |
| p-p38 | aBCR | 5min | PB | 8 | 0.245301 | 0.074504 | 0.19514 | 0.319589 |
| p-p38 | aBCR | 15min | naiveB | 38 | 0.111744 | 0.011929 | 0.074299 | 0.160034 |
| p-p38 | aBCR | 15min | DNB | 38 | 0.165207 | 0.01562 | 0.095814 | 0.241811 |
| p-p38 | aBCR | 15min | MZB | 30 | 0.253829 | 0.025553 | 0.165072 | 0.36835 |
| p-p38 | aBCR | 15min | swB | 38 | 0.225758 | 0.018406 | 0.178467 | 0.296115 |
| p-p38 | aBCR | 15min | PB | 8 | 0.23381 | 0.08884 | 0.077758 | 0.328103 |
| p-p38 | aBCR | 30min | naiveB | 38 | 0.075293 | 0.012086 | 0.006211 | 0.120159 |
| p-p38 | aBCR | 30min | DNB | 38 | 0.151841 | 0.013654 | 0.085739 | 0.206364 |
| p-p38 | aBCR | 30min | MZB | 28 | 0.180595 | 0.024164 | 0.097389 | 0.268967 |
| p-p38 | aBCR | 30min | swB | 37 | 0.178433 | 0.01683 | 0.099923 | 0.246606 |
| p-p38 | aBCR | 30min | PB | 10 | 0.28828 | 0.045761 | 0.216976 | 0.400588 |
| p-S6 | aBCR | unstim | naiveB | 37 | 0 | 0 | 0 | 0 |
| p-S6 | aBCR | unstim | DNB | 37 | 0 | 0 | 0 | 0 |
| p-S6 | aBCR | unstim | MZB | 35 | 0 | 0 | 0 | 0 |
| p-S6 | aBCR | unstim | swB | 37 | 0 | 0 | 0 | 0 |
| p-S6 | aBCR | unstim | PB | 11 | 0 | 0 | 0 | 0 |
| p-S6 | aBCR | 5min | naiveB | 38 | 0.114259 | 0.013124 | 0.064038 | 0.173095 |
| p-S6 | aBCR | 5min | DNB | 38 | 0.192683 | 0.028183 | 0.038017 | 0.29896 |
| p-S6 | aBCR | 5min | MZB | 35 | 0.206577 | 0.028021 | 0.074854 | 0.328232 |
| p-S6 | aBCR | 5min | swB | 38 | 0.324586 | 0.037215 | 0.14117 | 0.491833 |
| p-S6 | aBCR | 5min | PB | 8 | 0.216779 | 0.046107 | 0.117451 | 0.269316 |
| p-S6 | aBCR | 15min | naiveB | 38 | 0.474331 | 0.046474 | 0.279377 | 0.686176 |
| p-S6 | aBCR | 15min | DNB | 38 | 0.678294 | 0.076935 | 0.139403 | 1.076964 |
| p-S6 | aBCR | 15min | MZB | 30 | 0.907834 | 0.072298 | 0.702204 | 1.091687 |
| p-S6 | aBCR | 15min | swB | 38 | 1.148625 | 0.072963 | 0.83418 | 1.52743 |
| p-S6 | aBCR | 15min | PB | 8 | 1.037692 | 0.197767 | 0.698657 | 1.435645 |
| p-S6 | aBCR | 30min | naiveB | 38 | 0.705114 | 0.064716 | 0.501948 | 1.00461 |
| p-S6 | aBCR | 30min | DNB | 38 | 1.008439 | 0.080159 | 0.809173 | 1.396653 |
| p-S6 | aBCR | 30min | MZB | 28 | 1.329979 | 0.063495 | 1.185283 | 1.598388 |
| p-S6 | aBCR | 30min | swB | 37 | 1.515666 | 0.057926 | 1.337472 | 1.772909 |
| p-S6 | aBCR | 30min | PB | 10 | 1.313695 | 0.238975 | 0.82229 | 1.805787 |

**Supplementary Table 2H:** FIGURE 3BC_STATS.

| marker | Stimulation | Timing | Celltype_1 | Celltype_2 | p.adj | p.adj.signif |
| --- | --- | --- | --- | --- | --- | --- |
| **p-Btk** | **T105** | **5min** | **naiveB** | **DNB** | **0.22** | **ns** |
| **p-Btk** | **T105** | **5min** | **naiveB** | **MZB** | **0.0024** | **\*\*** |
| **p-Btk** | **T105** | **5min** | **naiveB** | **swB** | **0.00031** | **\*\*\*** |
| **p-Btk** | **T105** | **5min** | **naiveB** | **PB** | **0.064** | **ns** |
| **p-Btk** | **T105** | **5min** | **DNB** | **MZB** | **3.40E-07** | **\*\*\*\*** |
| **p-Btk** | **T105** | **5min** | **DNB** | **swB** | **1.80E-08** | **\*\*\*\*** |
| **p-Btk** | **T105** | **5min** | **DNB** | **PB** | **0.62** | **ns** |
| **p-Btk** | **T105** | **5min** | **MZB** | **swB** | **0.99** | **ns** |
| **p-Btk** | **T105** | **5min** | **MZB** | **PB** | **2.50E-05** | **\*\*\*\*** |
| **p-Btk** | **T105** | **5min** | **swB** | **PB** | **6.40E-06** | **\*\*\*\*** |
| p-Btk | T105 | 15min | naiveB | DNB | 0.64 | ns |
| p-Btk | T105 | 15min | naiveB | MZB | 2.10E-05 | **** |
| p-Btk | T105 | 15min | naiveB | swB | 7.00E-04 | *** |
| p-Btk | T105 | 15min | naiveB | PB | 0.0019 | ** |
| p-Btk | T105 | 15min | DNB | MZB | 4.60E-08 | **** |
| p-Btk | T105 | 15min | DNB | swB | 2.00E-06 | **** |
| p-Btk | T105 | 15min | DNB | PB | 0.027 | * |
| p-Btk | T105 | 15min | MZB | swB | 0.8 | ns |
| p-Btk | T105 | 15min | MZB | PB | 3.20E-09 | **** |
| p-Btk | T105 | 15min | swB | PB | 5.20E-08 | **** |
| p-Btk | T105 | 30min | naiveB | DNB | 0.87 | ns |
| p-Btk | T105 | 30min | naiveB | MZB | 1.90E-05 | **** |
| p-Btk | T105 | 30min | naiveB | swB | 0.00088 | *** |
| p-Btk | T105 | 30min | naiveB | PB | 4.20E-06 | **** |
| p-Btk | T105 | 30min | DNB | MZB | 3.30E-07 | **** |
| p-Btk | T105 | 30min | DNB | swB | 1.80E-05 | **** |
| p-Btk | T105 | 30min | DNB | PB | 6.40E-05 | **** |
| p-Btk | T105 | 30min | MZB | swB | 0.74 | ns |
| p-Btk | T105 | 30min | MZB | PB | 3.20E-13 | **** |
| p-Btk | T105 | 30min | swB | PB | 6.80E-12 | **** |
| **p-Akt** | **T105** | **5min** | **naiveB** | **DNB** | **1** | **ns** |
| **p-Akt** | **T105** | **5min** | **naiveB** | **MZB** | **8.70E-09** | **\*\*\*\*** |
| **p-Akt** | **T105** | **5min** | **naiveB** | **swB** | **2.60E-10** | **\*\*\*\*** |
| **p-Akt** | **T105** | **5min** | **naiveB** | **PB** | **9.20E-11** | **\*\*\*\*** |
| **p-Akt** | **T105** | **5min** | **DNB** | **MZB** | **1.40E-09** | **\*\*\*\*** |
| **p-Akt** | **T105** | **5min** | **DNB** | **swB** | **3.70E-11** | **\*\*\*\*** |
| **p-Akt** | **T105** | **5min** | **DNB** | **PB** | **2.80E-11** | **\*\*\*\*** |
| **p-Akt** | **T105** | **5min** | **MZB** | **swB** | **0.99** | **ns** |
| **p-Akt** | **T105** | **5min** | **MZB** | **PB** | **0.007** | **\*\*** |
| **p-Akt** | **T105** | **5min** | **swB** | **PB** | **0.017** | **\*** |
| p-Akt | T105 | 15min | naiveB | DNB | 0.99 | ns |
| p-Akt | T105 | 15min | naiveB | MZB | 1.10E-13 | **** |
| p-Akt | T105 | 15min | naiveB | swB | 4.50E-11 | **** |
| p-Akt | T105 | 15min | naiveB | PB | 8.10E-07 | **** |
| p-Akt | T105 | 15min | DNB | MZB | 5.20E-14 | **** |
| p-Akt | T105 | 15min | DNB | swB | 2.60E-12 | **** |
| p-Akt | T105 | 15min | DNB | PB | 1.90E-07 | **** |
| p-Akt | T105 | 15min | MZB | swB | 0.49 | ns |
| p-Akt | T105 | 15min | MZB | PB | 1 | ns |
| p-Akt | T105 | 15min | swB | PB | 0.75 | ns |
| p-Akt | T105 | 30min | naiveB | DNB | 1 | ns |
| p-Akt | T105 | 30min | naiveB | MZB | 4.40E-11 | **** |
| p-Akt | T105 | 30min | naiveB | swB | 9.00E-06 | **** |
| p-Akt | T105 | 30min | naiveB | PB | 6.90E-07 | **** |
| p-Akt | T105 | 30min | DNB | MZB | 2.60E-10 | **** |
| p-Akt | T105 | 30min | DNB | swB | 4.20E-05 | **** |
| p-Akt | T105 | 30min | DNB | PB | 2.00E-06 | **** |
| p-Akt | T105 | 30min | MZB | swB | 0.048 | * |
| p-Akt | T105 | 30min | MZB | PB | 1 | ns |
| p-Akt | T105 | 30min | swB | PB | 0.14 | ns |
| p-p38 | T105 | 5min | naiveB | DNB | 1 | ns |
| p-p38 | T105 | 5min | naiveB | MZB | 0.0011 | ** |
| p-p38 | T105 | 5min | naiveB | swB | 0.0023 | ** |
| p-p38 | T105 | 5min | naiveB | PB | 0.16 | ns |
| p-p38 | T105 | 5min | DNB | MZB | 0.0033 | ** |
| p-p38 | T105 | 5min | DNB | swB | 0.0067 | ** |
| p-p38 | T105 | 5min | DNB | PB | 0.24 | ns |
| p-p38 | T105 | 5min | MZB | swB | 1 | ns |
| p-p38 | T105 | 5min | MZB | PB | 1 | ns |
| p-p38 | T105 | 5min | swB | PB | 1 | ns |
| p-p38 | T105 | 15min | naiveB | DNB | 0.28 | ns |
| p-p38 | T105 | 15min | naiveB | MZB | 2.00E-05 | **** |
| p-p38 | T105 | 15min | naiveB | swB | 0.00039 | *** |
| p-p38 | T105 | 15min | naiveB | PB | 0.063 | ns |
| p-p38 | T105 | 15min | DNB | MZB | 0.02 | * |
| p-p38 | T105 | 15min | DNB | swB | 0.17 | ns |
| p-p38 | T105 | 15min | DNB | PB | 0.56 | ns |
| p-p38 | T105 | 15min | MZB | swB | 0.86 | ns |
| p-p38 | T105 | 15min | MZB | PB | 0.99 | ns |
| p-p38 | T105 | 15min | swB | PB | 1 | ns |
| p-p38 | T105 | 30min | naiveB | DNB | 0.0099 | ** |
| p-p38 | T105 | 30min | naiveB | MZB | 0.00043 | *** |
| p-p38 | T105 | 30min | naiveB | swB | 0.00017 | *** |
| p-p38 | T105 | 30min | naiveB | PB | 1.80E-07 | **** |
| p-p38 | T105 | 30min | DNB | MZB | 0.78 | ns |
| p-p38 | T105 | 30min | DNB | swB | 0.78 | ns |
| p-p38 | T105 | 30min | DNB | PB | 0.0018 | ** |
| p-p38 | T105 | 30min | MZB | swB | 1 | ns |
| p-p38 | T105 | 30min | MZB | PB | 0.034 | * |
| p-p38 | T105 | 30min | swB | PB | 0.021 | * |
| p-S6 | T105 | 5min | naiveB | DNB | 0.26 | ns |
| p-S6 | T105 | 5min | naiveB | MZB | 0.14 | ns |
| p-S6 | T105 | 5min | naiveB | swB | 2.30E-06 | **** |
| p-S6 | T105 | 5min | naiveB | PB | 0.53 | ns |
| p-S6 | T105 | 5min | DNB | MZB | 1 | ns |
| p-S6 | T105 | 5min | DNB | swB | 0.0077 | ** |
| p-S6 | T105 | 5min | DNB | PB | 1 | ns |
| p-S6 | T105 | 5min | MZB | swB | 0.028 | * |
| p-S6 | T105 | 5min | MZB | PB | 1 | ns |
| p-S6 | T105 | 5min | swB | PB | 0.48 | ns |
| p-S6 | T105 | 15min | naiveB | DNB | 0.21 | ns |
| p-S6 | T105 | 15min | naiveB | MZB | 0.00035 | *** |
| p-S6 | T105 | 15min | naiveB | swB | 6.60E-10 | **** |
| p-S6 | T105 | 15min | naiveB | PB | 0.006 | ** |
| p-S6 | T105 | 15min | DNB | MZB | 0.17 | ns |
| p-S6 | T105 | 15min | DNB | swB | 2.30E-05 | **** |
| p-S6 | T105 | 15min | DNB | PB | 0.18 | ns |
| p-S6 | T105 | 15min | MZB | swB | 0.13 | ns |
| p-S6 | T105 | 15min | MZB | PB | 0.94 | ns |
| p-S6 | T105 | 15min | swB | PB | 0.96 | ns |
| p-S6 | T105 | 30min | naiveB | DNB | 0.023 | * |
| p-S6 | T105 | 30min | naiveB | MZB | 4.50E-07 | **** |
| p-S6 | T105 | 30min | naiveB | swB | 2.40E-12 | **** |
| p-S6 | T105 | 30min | naiveB | PB | 0.0012 | ** |
| p-S6 | T105 | 30min | DNB | MZB | 0.028 | * |
| p-S6 | T105 | 30min | DNB | swB | 1.30E-05 | **** |
| p-S6 | T105 | 30min | DNB | PB | 0.28 | ns |
| p-S6 | T105 | 30min | MZB | swB | 0.43 | ns |
| p-S6 | T105 | 30min | MZB | PB | 1 | ns |
| p-S6 | T105 | 30min | swB | PB | 0.69 | ns |

**Supplementary Table 2I:** FIGURE 3EF_AUC_DATA.

| marker | Stimulation | cells | Count | Median_AI | Q25 | Q75 | p.adj | p.adj.signif |
| --- | --- | --- | --- | --- | --- | --- | --- | --- |
| p-ITK | aTCR | naive4 | 38 | 1.399297 | 1.056442 | 1.831197 | NA | NA |
| p-ITK | aTCR | nonnaive4 | 38 | 1.137418 | 0.868259 | 1.385741 | 0.0075 | ** |
| p-ITK | aTCR | naive8 | 38 | 0.882749 | 0.649251 | 1.100488 | NA | NA |
| p-ITK | aTCR | nonnaive8 | 38 | 0.776246 | 0.621721 | 0.945542 | 0.431 | ns |

**Supplementary Table 2J:**
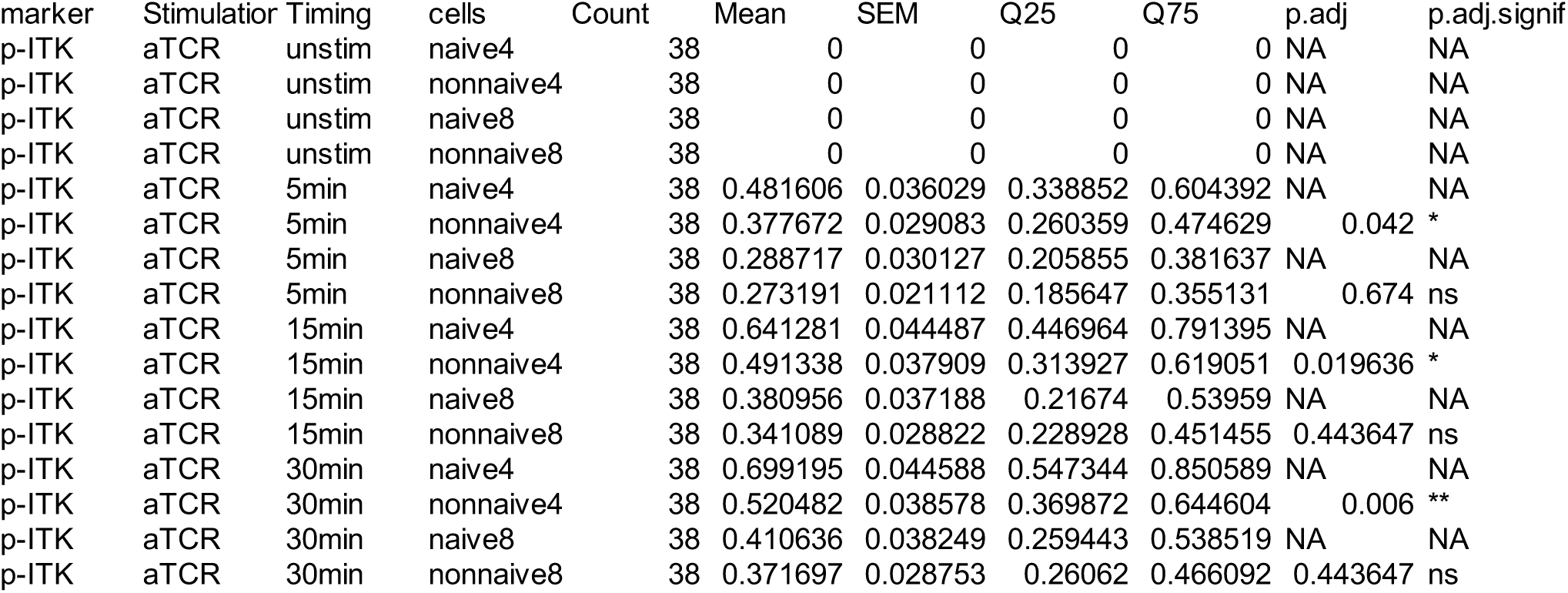
FIGURE 3EF_LINEPLOT_DATA.

**Supplementary Table 2K:**
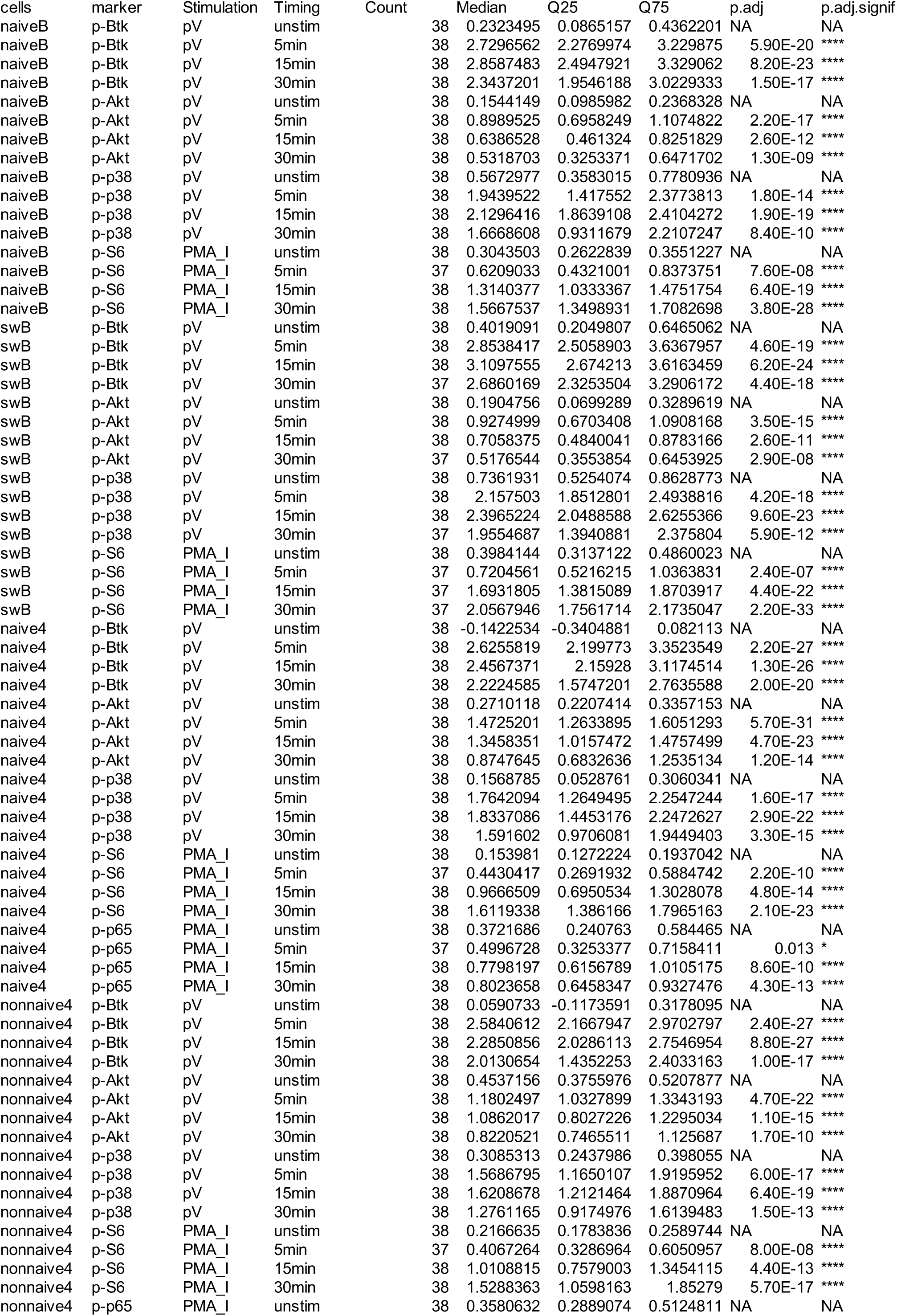
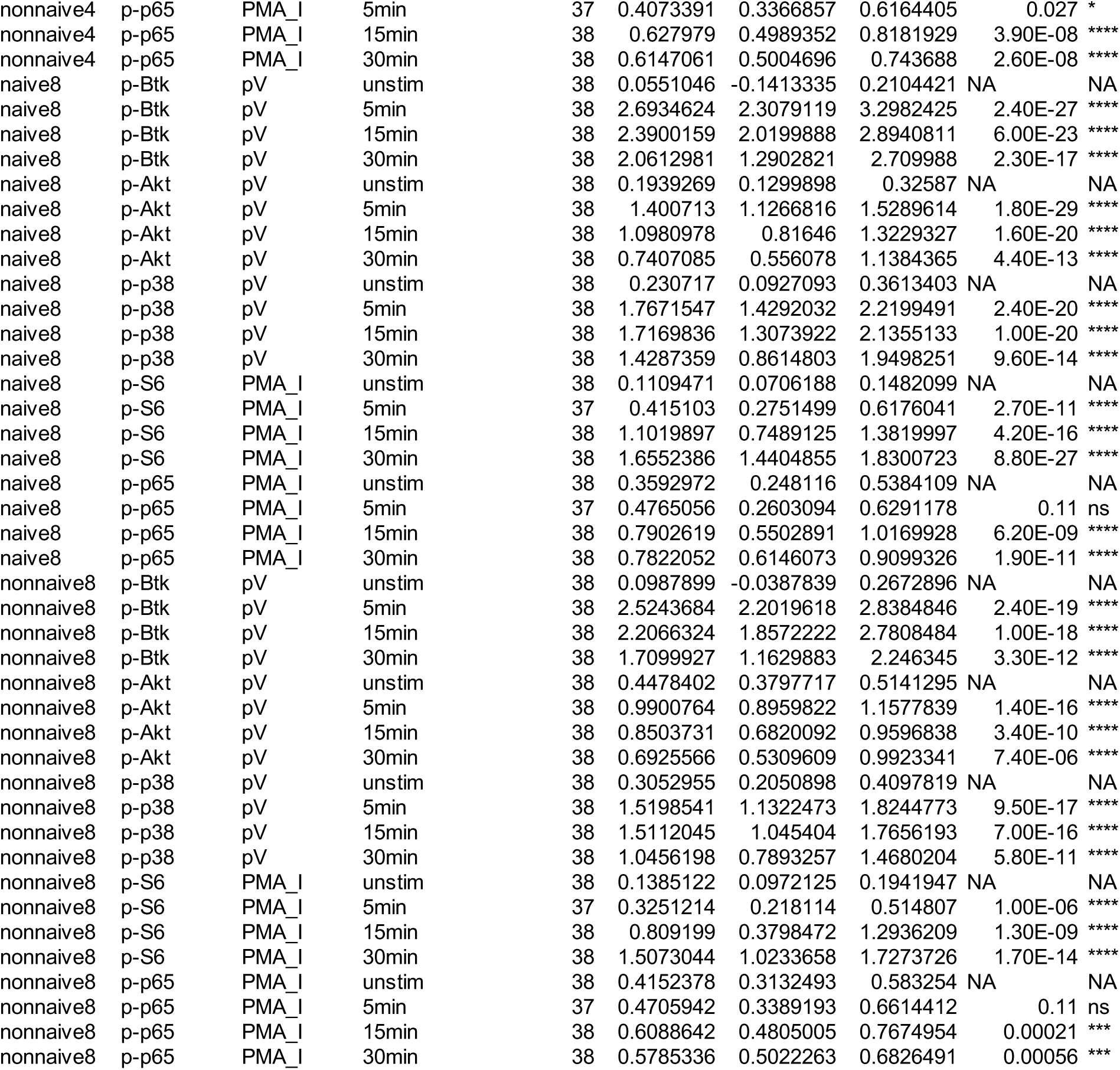
SF4_MFI_DATA.

**Supplementary Table 2L:**
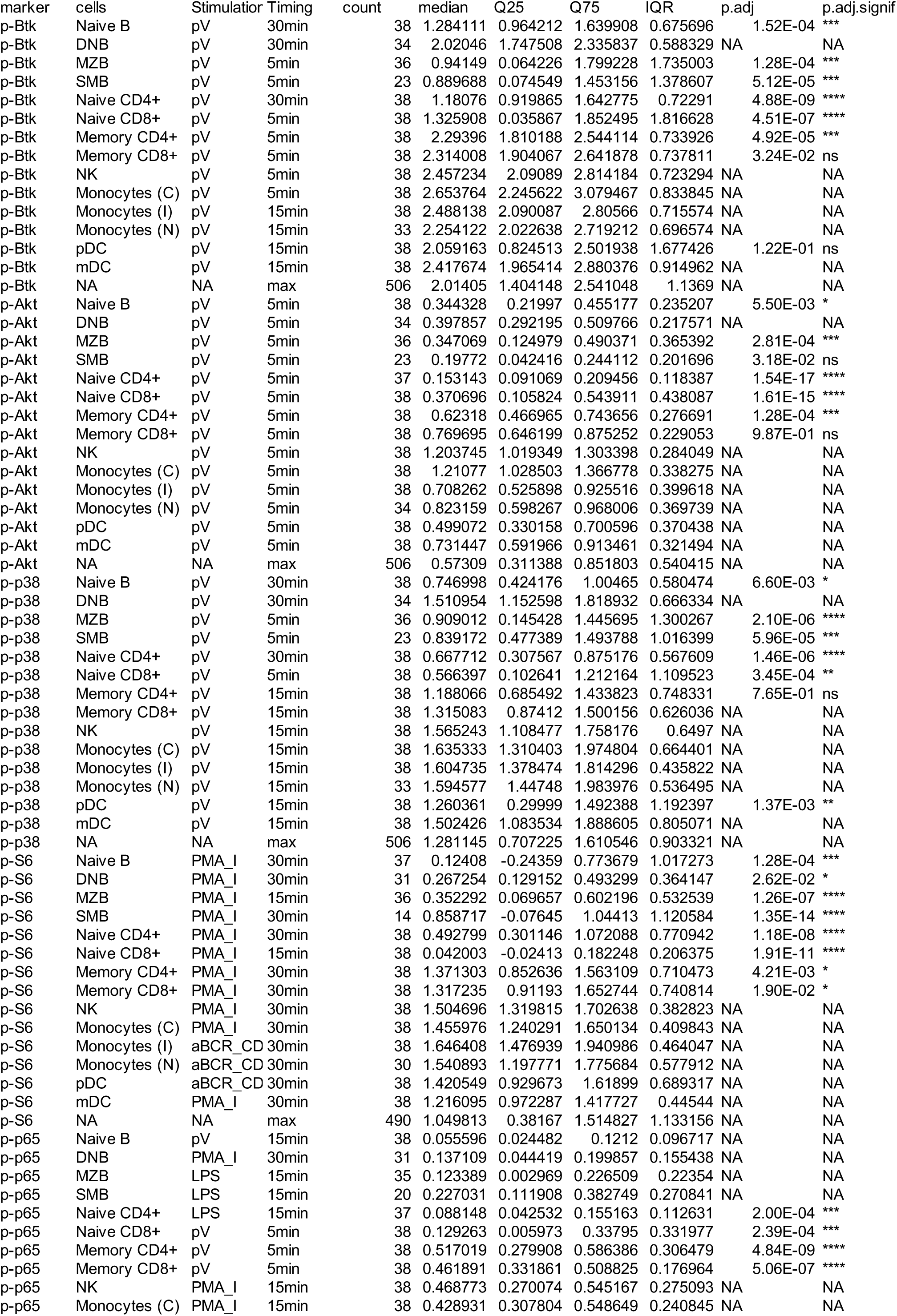
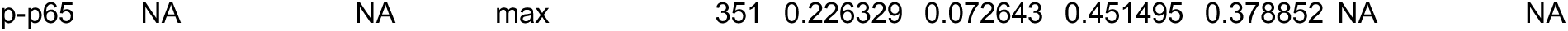
FIGURE4A_DATA.

**Supplementary Table 2M:**
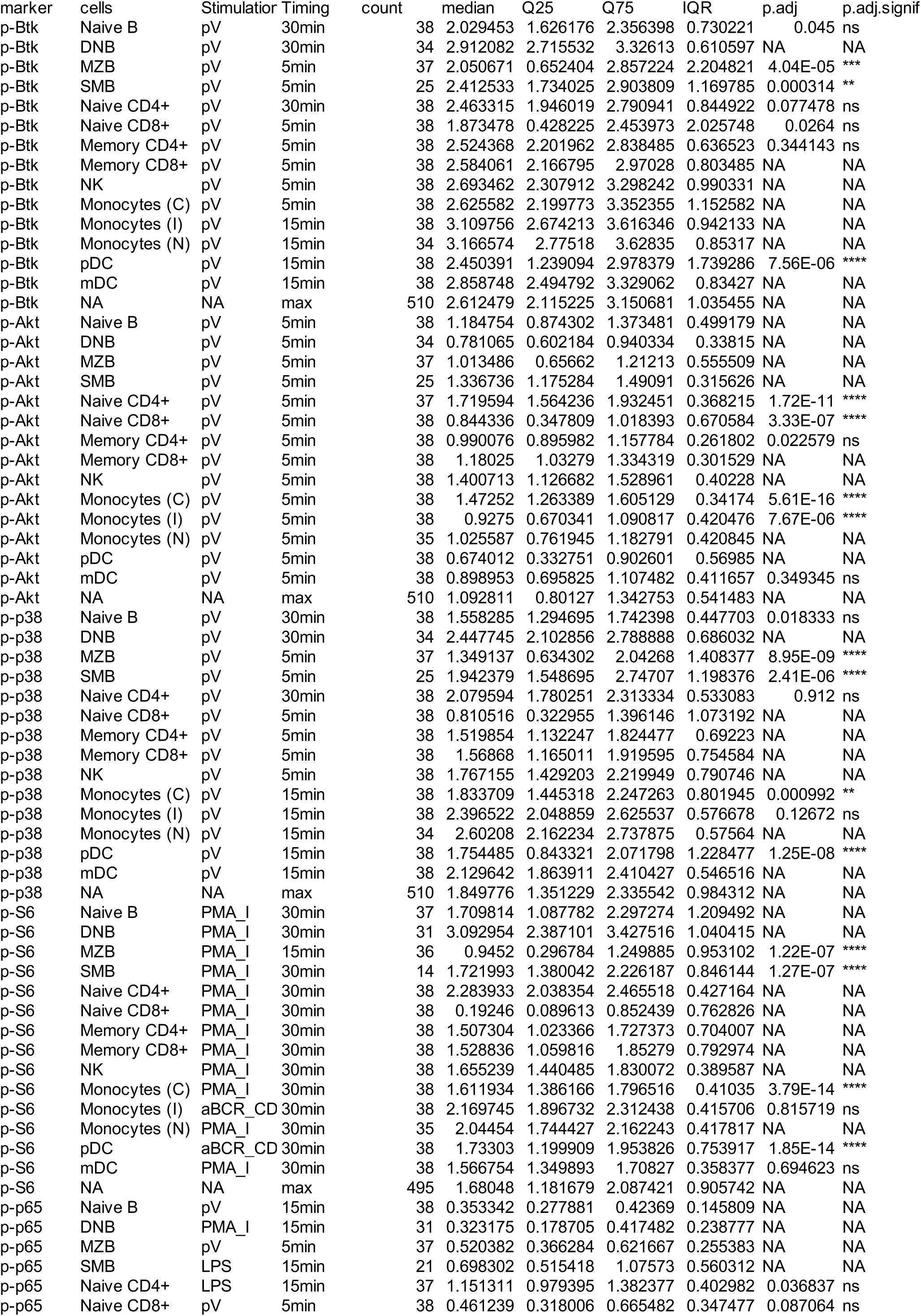
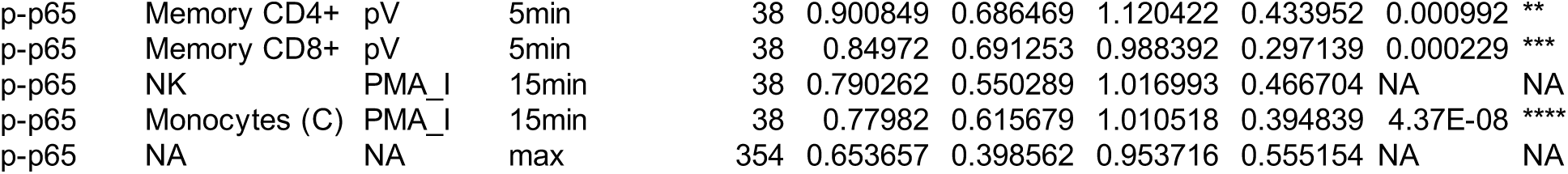
FIGURE4B_DATA.

**Supplementary Table 2N:** FIGURE5B_DATA.

| cells | marker | Stimulation | Timing | Agegroup | Count | Mean | SEM | Q25 | Q75 |
| --- | --- | --- | --- | --- | --- | --- | --- | --- | --- |
| naiveB | p-BTK | aBCR | unstim | 0-18 | 10 | 0 | 0 | 0 | 0 |
| naiveB | p-BTK | aBCR | unstim | 18-55 | 37 | 0 | 0 | 0 | 0 |
| naiveB | p-BTK | aBCR | unstim | 56+ | 6 | 0 | 0 | 0 | 0 |
| naiveB | p-BTK | aBCR | 5min | 0-18 | 10 | 1.188088 | 0.105214 | 0.918892 | 1.345357 |
| naiveB | p-BTK | aBCR | 5min | 18-55 | 38 | 0.727784 | 0.055106 | 0.456346 | 0.985005 |
| naiveB | p-BTK | aBCR | 5min | 56+ | 6 | 0.751254 | 0.152145 | 0.711516 | 0.790418 |
| naiveB | p-BTK | aBCR | 15min | 0-18 | 8 | 1.076332 | 0.089034 | 0.945853 | 1.229517 |
| naiveB | p-BTK | aBCR | 15min | 18-55 | 38 | 0.87007 | 0.051303 | 0.617344 | 1.068877 |
| naiveB | p-BTK | aBCR | 15min | 56+ | 6 | 0.996891 | 0.134425 | 0.939518 | 1.16914 |
| naiveB | p-BTK | aBCR | 30min | 0-18 | 10 | 1.036654 | 0.100784 | 0.832944 | 1.194809 |
| naiveB | p-BTK | aBCR | 30min | 18-55 | 38 | 0.89499 | 0.049798 | 0.747024 | 1.091246 |
| naiveB | p-BTK | aBCR | 30min | 56+ | 6 | 0.978331 | 0.122498 | 0.773054 | 1.177658 |
| naiveB | p-Akt | aBCR | unstim | 0-18 | 10 | 0 | 0 | 0 | 0 |
| naiveB | p-Akt | aBCR | unstim | 18-55 | 37 | 0 | 0 | 0 | 0 |
| naiveB | p-Akt | aBCR | unstim | 56+ | 6 | 0 | 0 | 0 | 0 |
| naiveB | p-Akt | aBCR | 5min | 0-18 | 10 | 0.531555 | 0.0312 | 0.468492 | 0.61004 |
| naiveB | p-Akt | aBCR | 5min | 18-55 | 38 | 0.382363 | 0.016571 | 0.314153 | 0.454679 |
| naiveB | p-Akt | aBCR | 5min | 56+ | 6 | 0.459519 | 0.074983 | 0.419135 | 0.516971 |
| naiveB | p-Akt | aBCR | 15min | 0-18 | 8 | 0.493822 | 0.024036 | 0.451876 | 0.525705 |
| naiveB | p-Akt | aBCR | 15min | 18-55 | 38 | 0.405124 | 0.016276 | 0.343673 | 0.472517 |
| naiveB | p-Akt | aBCR | 15min | 56+ | 6 | 0.481517 | 0.068158 | 0.416172 | 0.537804 |
| naiveB | p-Akt | aBCR | 30min | 0-18 | 10 | 0.403668 | 0.025531 | 0.375063 | 0.439228 |
| naiveB | p-Akt | aBCR | 30min | 18-55 | 38 | 0.366147 | 0.016998 | 0.335035 | 0.43955 |
| naiveB | p-Akt | aBCR | 30min | 56+ | 6 | 0.407988 | 0.050703 | 0.366306 | 0.45566 |
| naiveB | p-p38 | aBCR | unstim | 0-18 | 10 | 0 | 0 | 0 | 0 |
| naiveB | p-p38 | aBCR | unstim | 18-55 | 37 | 0 | 0 | 0 | 0 |
| naiveB | p-p38 | aBCR | unstim | 56+ | 6 | 0 | 0 | 0 | 0 |
| naiveB | p-p38 | aBCR | 5min | 0-18 | 10 | 0.151652 | 0.030097 | 0.078053 | 0.216324 |
| naiveB | p-p38 | aBCR | 5min | 18-55 | 38 | 0.130808 | 0.01532 | 0.076985 | 0.174377 |
| naiveB | p-p38 | aBCR | 5min | 56+ | 6 | 0.125701 | 0.03334 | 0.143568 | 0.167114 |
| naiveB | p-p38 | aBCR | 15min | 0-18 | 8 | 0.111183 | 0.02314 | 0.063288 | 0.149616 |
| naiveB | p-p38 | aBCR | 15min | 18-55 | 38 | 0.111744 | 0.011929 | 0.074299 | 0.160034 |
| naiveB | p-p38 | aBCR | 15min | 56+ | 6 | 0.108636 | 0.037609 | 0.094215 | 0.154605 |
| naiveB | p-p38 | aBCR | 30min | 0-18 | 10 | 0.065649 | 0.019725 | 0.030299 | 0.08777 |
| naiveB | p-p38 | aBCR | 30min | 18-55 | 38 | 0.075293 | 0.012086 | 0.006211 | 0.120159 |
| naiveB | p-p38 | aBCR | 30min | 56+ | 6 | 0.068026 | 0.037904 | 0.036317 | 0.117201 |
| naiveB | p-S6 | aBCR | unstim | 0-18 | 10 | 0 | 0 | 0 | 0 |
| naiveB | p-S6 | aBCR | unstim | 18-55 | 37 | 0 | 0 | 0 | 0 |
| naiveB | p-S6 | aBCR | unstim | 56+ | 6 | 0 | 0 | 0 | 0 |
| naiveB | p-S6 | aBCR | 5min | 0-18 | 10 | 0.268203 | 0.034805 | 0.220437 | 0.356091 |
| naiveB | p-S6 | aBCR | 5min | 18-55 | 38 | 0.114259 | 0.013124 | 0.064038 | 0.173095 |
| naiveB | p-S6 | aBCR | 5min | 56+ | 6 | 0.093706 | 0.034782 | 0.086095 | 0.14684 |
| naiveB | p-S6 | aBCR | 15min | 0-18 | 8 | 0.731553 | 0.106311 | 0.637174 | 0.908359 |
| naiveB | p-S6 | aBCR | 15min | 18-55 | 38 | 0.474331 | 0.046474 | 0.279377 | 0.686176 |
| naiveB | p-S6 | aBCR | 15min | 56+ | 6 | 0.572727 | 0.150071 | 0.369474 | 0.87246 |
| naiveB | p-S6 | aBCR | 30min | 0-18 | 10 | 1.105647 | 0.122865 | 0.919728 | 1.289523 |
| naiveB | p-S6 | aBCR | 30min | 18-55 | 38 | 0.705114 | 0.064716 | 0.501948 | 1.00461 |
| naiveB | p-S6 | aBCR | 30min | 56+ | 6 | 0.662249 | 0.139795 | 0.393694 | 0.867924 |
| swB | p-BTK | aBCR | unstim | 0-18 | 9 | 0 | 0 | 0 | 0 |
| swB | p-BTK | aBCR | unstim | 18-55 | 37 | 0 | 0 | 0 | 0 |
| swB | p-BTK | aBCR | unstim | 56+ | 6 | 0 | 0 | 0 | 0 |
| swB | p-BTK | aBCR | 5min | 0-18 | 9 | 1.178913 | 0.102513 | 0.931362 | 1.38818 |
| swB | p-BTK | aBCR | 5min | 18-55 | 38 | 1.097034 | 0.065066 | 0.914346 | 1.368893 |
| swB | p-BTK | aBCR | 5min | 56+ | 6 | 1.160466 | 0.142744 | 1.03477 | 1.390897 |
| swB | p-BTK | aBCR | 15min | 0-18 | 8 | 1.080167 | 0.093282 | 0.907503 | 1.280667 |
| swB | p-BTK | aBCR | 15min | 18-55 | 38 | 1.211093 | 0.058758 | 0.956338 | 1.419907 |
| swB | p-BTK | aBCR | 15min | 56+ | 6 | 1.322791 | 0.077656 | 1.239624 | 1.471534 |
| swB | p-BTK | aBCR | 30min | 0-18 | 9 | 0.936522 | 0.095104 | 0.650702 | 1.084685 |
| swB | p-BTK | aBCR | 30min | 18-55 | 37 | 1.176094 | 0.046558 | 0.997741 | 1.355772 |
| swB | p-BTK | aBCR | 30min | 56+ | 6 | 1.227649 | 0.061928 | 1.141729 | 1.269574 |
| swB | p-Akt | aBCR | unstim | 0-18 | 9 | 0 | 0 | 0 | 0 |
| swB | p-Akt | aBCR | unstim | 18-55 | 37 | 0 | 0 | 0 | 0 |
| swB | p-Akt | aBCR | unstim | 56+ | 6 | 0 | 0 | 0 | 0 |
| swB | p-Akt | aBCR | 5min | 0-18 | 9 | 0.699768 | 0.058051 | 0.604943 | 0.811719 |
| swB | p-Akt | aBCR | 5min | 18-55 | 38 | 0.590865 | 0.01916 | 0.514098 | 0.662613 |
| swB | p-Akt | aBCR | 5min | 56+ | 6 | 0.6609 | 0.082858 | 0.591859 | 0.716886 |
| swB | p-Akt | aBCR | 15min | 0-18 | 8 | 0.633995 | 0.044688 | 0.564691 | 0.68393 |
| swB | p-Akt | aBCR | 15min | 18-55 | 38 | 0.611627 | 0.018473 | 0.539559 | 0.682999 |
| swB | p-Akt | aBCR | 15min | 56+ | 6 | 0.642057 | 0.071595 | 0.52713 | 0.710514 |
| swB | p-Akt | aBCR | 30min | 0-18 | 9 | 0.43331 | 0.048242 | 0.397659 | 0.527934 |
| swB | p-Akt | aBCR | 30min | 18-55 | 37 | 0.507727 | 0.019461 | 0.442123 | 0.565084 |
| swB | p-Akt | aBCR | 30min | 56+ | 6 | 0.531319 | 0.039416 | 0.477306 | 0.579775 |
| swB | p-p38 | aBCR | unstim | 0-18 | 9 | 0 | 0 | 0 | 0 |
| swB | p-p38 | aBCR | unstim | 18-55 | 37 | 0 | 0 | 0 | 0 |
| swB | p-p38 | aBCR | unstim | 56+ | 6 | 0 | 0 | 0 | 0 |
| swB | p-p38 | aBCR | 5min | 0-18 | 9 | 0.195849 | 0.036262 | 0.148835 | 0.262586 |
| swB | p-p38 | aBCR | 5min | 18-55 | 38 | 0.242908 | 0.021448 | 0.13055 | 0.337834 |
| swB | p-p38 | aBCR | 5min | 56+ | 6 | 0.256253 | 0.044877 | 0.209303 | 0.308501 |
| swB | p-p38 | aBCR | 15min | 0-18 | 8 | 0.170284 | 0.032154 | 0.100041 | 0.202609 |
| swB | p-p38 | aBCR | 15min | 18-55 | 38 | 0.225758 | 0.018406 | 0.178467 | 0.296115 |
| swB | p-p38 | aBCR | 15min | 56+ | 6 | 0.283218 | 0.060413 | 0.211237 | 0.395953 |
| swB | p-p38 | aBCR | 30min | 0-18 | 9 | 0.103881 | 0.041293 | 0.043018 | 0.167833 |
| swB | p-p38 | aBCR | 30min | 18-55 | 37 | 0.178433 | 0.01683 | 0.099923 | 0.246606 |
| swB | p-p38 | aBCR | 30min | 56+ | 6 | 0.207836 | 0.054584 | 0.118231 | 0.301348 |
| swB | p-S6 | aBCR | unstim | 0-18 | 9 | 0 | 0 | 0 | 0 |
| swB | p-S6 | aBCR | unstim | 18-55 | 37 | 0 | 0 | 0 | 0 |
| swB | p-S6 | aBCR | unstim | 56+ | 6 | 0 | 0 | 0 | 0 |
| swB | p-S6 | aBCR | 5min | 0-18 | 9 | 0.439457 | 0.063079 | 0.453233 | 0.588387 |
| swB | p-S6 | aBCR | 5min | 18-55 | 38 | 0.324586 | 0.037215 | 0.14117 | 0.491833 |
| swB | p-S6 | aBCR | 5min | 56+ | 6 | 0.344709 | 0.050108 | 0.321672 | 0.330449 |
| swB | p-S6 | aBCR | 15min | 0-18 | 8 | 1.271341 | 0.152824 | 0.904855 | 1.652162 |
| swB | p-S6 | aBCR | 15min | 18-55 | 38 | 1.148625 | 0.072963 | 0.83418 | 1.52743 |
| swB | p-S6 | aBCR | 15min | 56+ | 6 | 1.246245 | 0.137216 | 1.208046 | 1.449414 |
| swB | p-S6 | aBCR | 30min | 0-18 | 9 | 1.545963 | 0.13851 | 1.351108 | 1.908117 |
| swB | p-S6 | aBCR | 30min | 18-55 | 37 | 1.515666 | 0.057926 | 1.337472 | 1.772909 |
| swB | p-S6 | aBCR | 30min | 56+ | 6 | 1.378819 | 0.114468 | 1.363224 | 1.470904 |

**Supplementary Table 2O:**
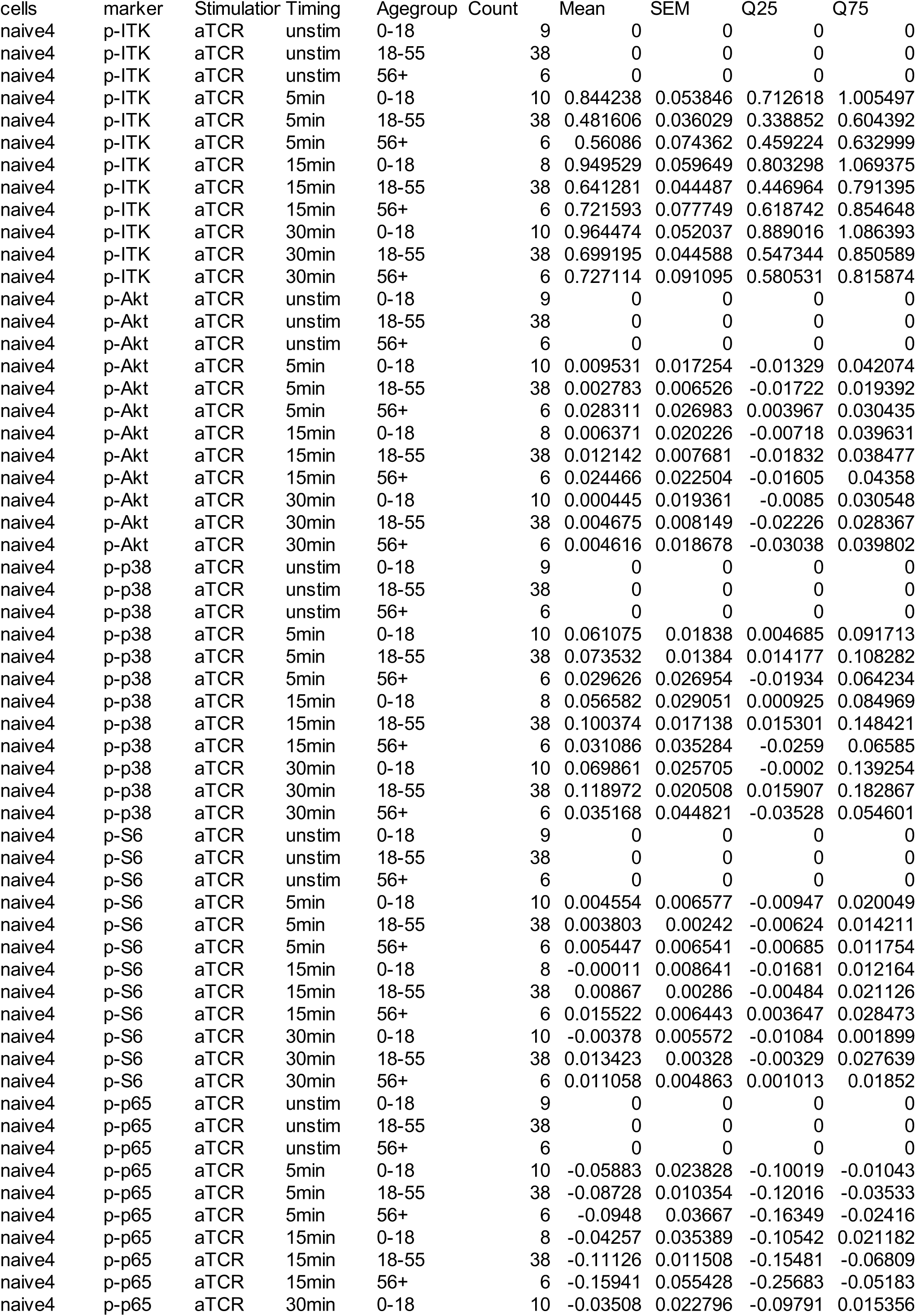
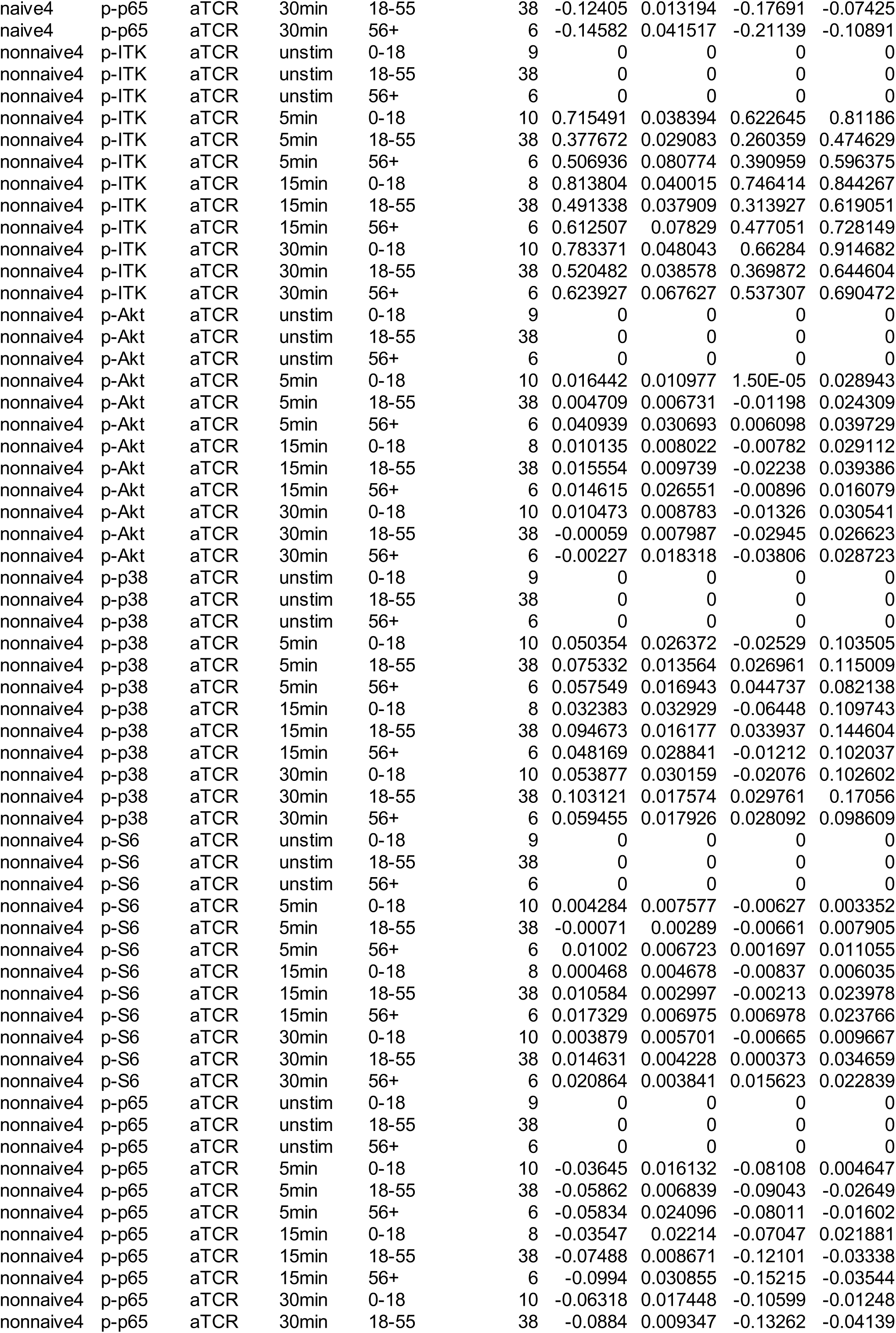
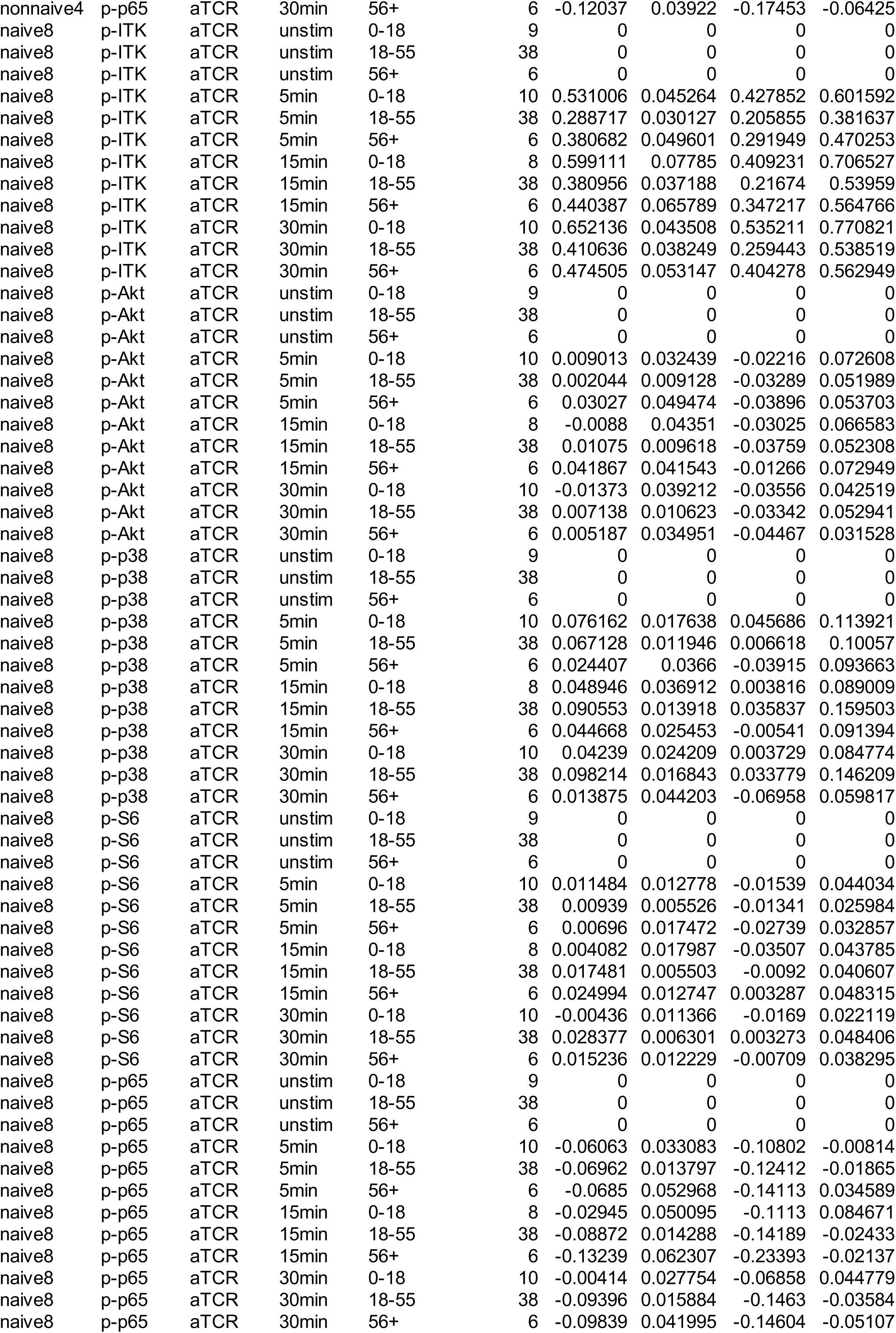
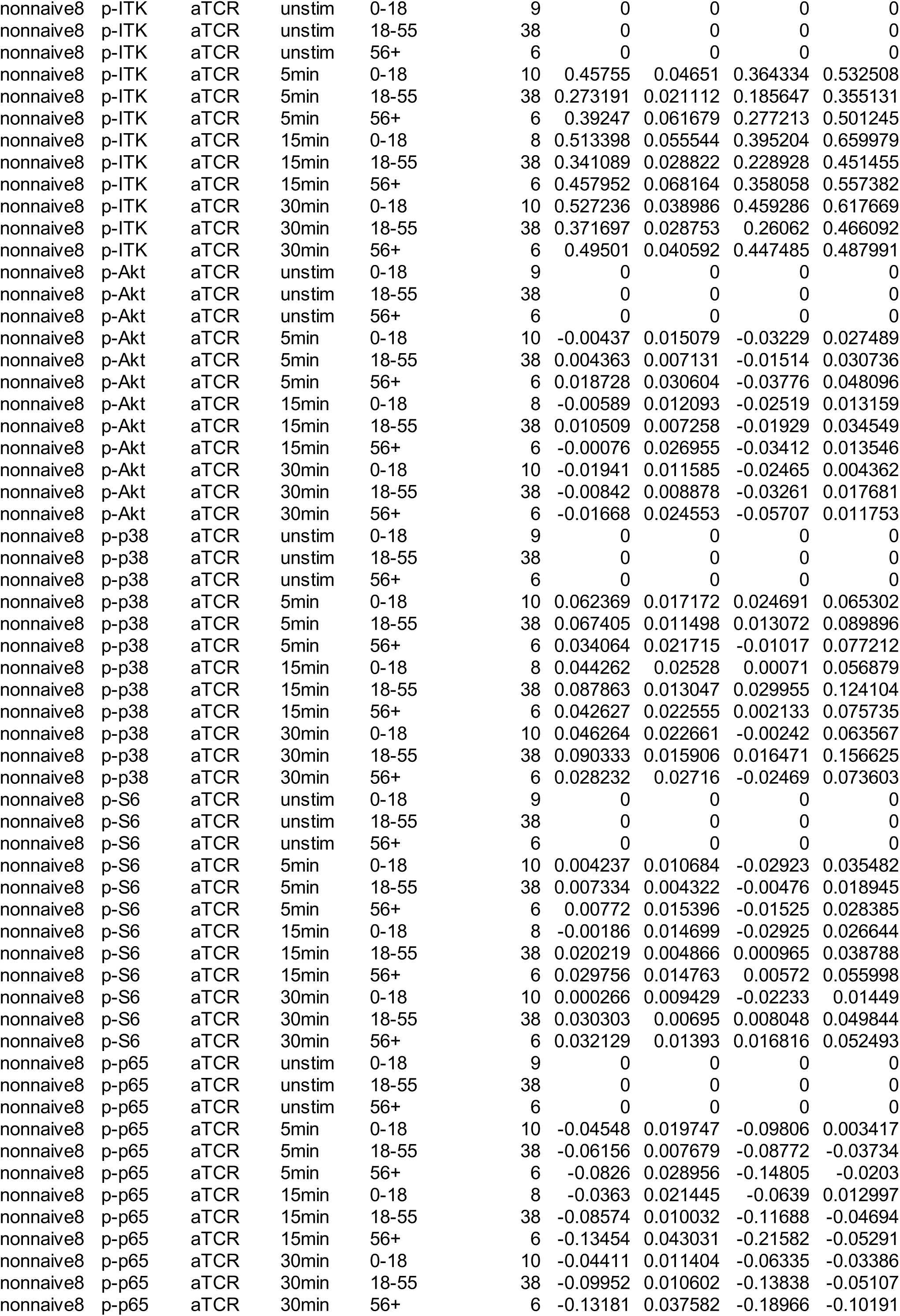
FIGURE5CD_DATA.

